# Fast and Accurate Genomic Analyses using Genome Graphs

**DOI:** 10.1101/194530

**Authors:** Goran Rakocevic, Vladimir Semenyuk, James Spencer, John Browning, Ivan Johnson, Vladan Arsenijevic, Jelena Nadj, Kaushik Ghose, Maria C. Suciu, Sun-Gou Ji, Gülfem Demir, Lizao Li, Berke Ç. Toptaş, Alexey Dolgoborodov, Björn Pollex, Iosif Spulber, Irina Glotova, Péter Kómár, Andrew Stachyra, Yilong Li, Milos Popovic, Wan-Ping Lee, Morten Källberg, Amit Jain, Deniz Kural

## Abstract

The human reference genome serves as the foundation for genomics by providing a scaffold for alignment of sequencing reads, but currently only reflects a single consensus haplotype, which impairs read alignment and downstream analysis accuracy. Reference genome structures incorporating known genetic variation have been shown to improve the accuracy of genomic analyses, but have so far remained computationally prohibitive for routine large-scale use. Here we present a graph genome implementation that enables read alignment across 2,800 diploid genomes encompassing 12.6 million SNPs and 4.0 million indels. Our Graph Genome Pipeline requires 6.5 hours to process a 30x coverage WGS sample on a system with 36 CPU cores compared with 11 hours required by the GATK Best Practices pipeline. Using complementary benchmarking experiments based on real and simulated data, we show that using a graph genome reference improves read mapping sensitivity and produces a 0.5% increase in variant calling recall, or about 20,000 additional variants being detected per sample, while variant calling specificity is unaffected. Structural variations (SVs) incorporated into a graph genome can be genotyped accurately under a unified framework. Finally, we show that iterative augmentation of graph genomes yields incremental gains in variant calling accuracy. Our implementation is a significant advance towards fulfilling the promise of graph genomes to radically enhance the scalability and accuracy of genomic analyses.

## Introduction

Completion of the first draft of the human reference genome^1,2^ was a landmark achievement in human genetics, establishing a standardized coordinate system for annotating genomic elements and comparing individual human genomes. The reference genome serves as a scaffold for mapping and assembling short DNA sequencing reads into longer consensus contigs, thus underpinning the quality of all ensuing analyses and ultimately the ability to draw conclusions of clinical significance from DNA sequencing.

The current human reference genome is represented as a linear haploid DNA sequence^3^. This structure poses theoretical limitations due to the prevalence of genetic diversity in human populations: any given human genome has on average 3.5-4.0 million single nucleotide polymorphisms (SNPs) or small insertions and deletions (INDELs) and around 2,500 large structural variations (SVs) compared with the reference genome^4,5^, including sizable genomic segments missing from the reference^6^. This genetic divergence may cause sequencing reads to map incorrectly or fail to map altogether^7,8^, particularly when they span SV breakpoints. Read mapping accuracy thus varies significantly across genomic regions in a given sample, and across genetically diverged samples. Misplaced reads may in turn result in both missed true variants (false negatives) and incorrectly reported false variants (false positives), as well as hamper other applications that rely on accurate read mapping, including RNA-sequencing analysis, chromatin immunoprecipitation-sequencing (ChIP-seq) analysis and copy number variation detection^7,8^. Identifying SVs is particularly challenging: despite the large number of SVs already characterized^5^, most methods for genotyping SVs still rely on detecting complex combinations of abnormal read alignment patterns to detect SVs9,10, although more recent algorithms such as BayesTyper11 and Graphtyper12 can take into account known SVs.

The linear reference genome represents just one of the possible alleles at any given site. Since mapping algorithms are based on the similarity to the reference sequence, reads originating from an alternative allele have a higher chance of being mismapped, an effect referred to as “reference bias”. Using population-specific major allele references^11,13–15^ would mitigate this effect but not eliminate it, since the majority of the variants with respect to the standard human reference genome have a <50% allele frequency of any given population (Supplementary Fig. 1). Moreover, having distinct population-specific reference genomes with different coordinate systems would complicate any genomic analyses comprising multiple populations. An alternative approach to capture the genetic diversity in the human reference genome is to augment it with alternate haplotype sequences at genetically diverged loci^16^. This approach has been applied since the previous human reference version, GRCh37^16^, but introduces inefficiency through sequence duplication between the main linear genome and the alternate haplotypes.

Recent large-scale resequencing efforts have comprehensively cataloged common genetic variants^4,17,18^, prompting suggestions to make use of this information through multi-genome references^19,20^, which have been suggested to alleviate reference bias by facilitating read mapping^21,22^. Despite these promising observations, currently available implementations of multi-genome graph references are either orders of magnitude slower than conventional linear reference genome-based methods on human WGS data (one example is BWBBLE^23^) or are intended for use with small genomes^19^ and small regions within large genomes^12,21,22,24^. Graphtyper^12^ is a recently published tool that performs local realignment of reads initially aligned by a linear aligner. This local alignment is performed in ∼50kb sliding windows, which avoids the computational complexity of using a WGS-sized pan-genome, but as a trade-off suffers from reference bias in genomewide read placement and mapping quality determination. The full genome-wide impact of using multi-genome references for human genomic analyses has not been assessed yet, although whole-genome workflows using graph genomes are under active development^25,26^.

Here we present a graph genome pipeline for building, augmenting, storing, querying and variant calling from graph genomes composed of a population of genome sequences. Our algorithms are, to our knowledge, the first graph genome implementation that achieve comparable reference indexing and read alignment speed to BWA-MEM^27^, a widely-used linear reference genome aligner. We show that graph genomes improve the mapping accuracy of next-generation sequencing (NGS) reads on the genome-wide level. Our NGS read alignment and variant calling pipeline, Graph Genome Pipeline, leverages our graph genome data structure and outperforms the state of the art linear reference-genome pipeline^28^ comprising BWA-MEM and GATK HaplotypeCaller^28^, as measured by multiple complementary benchmarks. By including breakpoint-resolved SV polymorphisms into the graph genome, we demonstrate that SVs can be genotyped rapidly and accurately in a unified fashion. As novel genetic variation data are accumulated in graph genomes, incremental improvements in read mapping and variant calling accuracy can be achieved. This will allow our approach to scale and improve with expanding genetic variation catalogs.

### A Computationally Efficient Graph Genome Implementation

We implemented a graph genome data structure that represents genomic sequences on the edges of the graph (Fig. 1a, Supplementary Information). A graph genome is constructed from a population of genome sequences, such that each haploid genome in this population is represented by a sequence path through the graph. To facilitate the use of widely available datasets, we implemented a process to build a graph genome using VCF files indicating genetic variants with respect to a standard linear reference genome, which is provided using a FASTA file. Thus, the standard linear reference genome is just one of the paths through the graph genome. For representational purposes, the linear reference genome path is labeled as the initial edge, and all coordinates of genetic variants are reported with respect to it. This ensures backward compatibility of graph coordinates to linear reference genome coordinates. In practice, a graph genome is built by iteratively adding edges corresponding to a non-reference allele, terminating at nodes corresponding to genomic loci on the initial edge. Insertions are represented as cyclic edges starting and terminating at the same node, but our mapping algorithm enforces acyclic traversal of the graph (Fig. 1a) by requiring that a path traverses a cycle at most once and traverses at most one cycle from each vertex. Genomic features such as tandem repeats, expansions and inversions are represented as insertions or sequence replacements in the graph. Variants are inserted into existing variant branches in a backward compatible manner, enabling variant discovery and representation in genomic regions absent from the linear reference genome^6^ (Fig. 1a). Our graph genome infrastructure supports building and aligning reads against general graph topologies such as hierarchical variation (Fig. 1a, Supplementary Fig. 5). However, unlike VG and HISAT2, we do not support bidirectionality or cycles, since such structures impose additional computational complexity; our directed acyclic graph is able to fully describe all genetic variation encapsulated in the sequential representation of nucleotides comprising chromosomes. For querying a graph genome, we use a hash table that associates short sequences of length *k* (*k*-mers) along all valid paths in the graph with their graph coordinates (Fig. 1b). Uninformative *k*-mers that occur exceptionally frequently are omitted (Supplementary Information).

**Figure 1.**
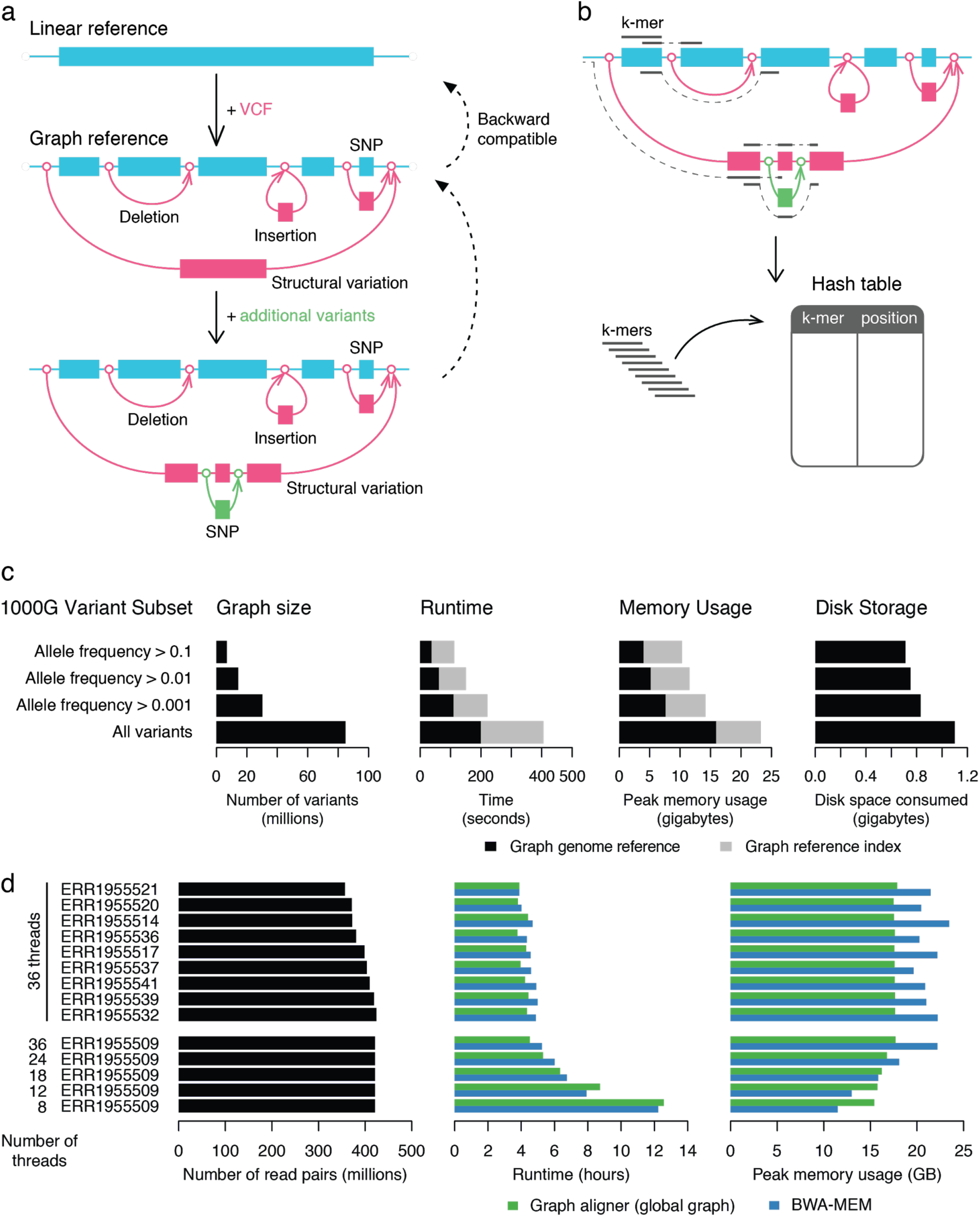
The graph genome architecture and computational resource requirements. (**a**) A graph genome is constructed from a standard linear reference genome FASTA file augmented by a set of genetic variants provided in VCF format. A graph genome can further be augmented with additional genetic variants in a second VCF file, or in the case of variants within variants using Graph Genome Pipeline. The coordinate system of each constructed graph genome is backward compatible with that of the linear reference genome. Each segment between vertices corresponds to an edge in the graph; inserting a variant to the graph can therefore add two (for insertions) or three (other variants) edges to the graph as the original edge is split into separate edges (as needed) at the start and end vertices of the new edge as well as the edge corresponding to the additional variant. Hence the three graphs shown contain 1, 9 and 13 edges respectively. A rendering of these graphs constructed using Graph Genome Pipeline is shown in Supplementary Figure 5. (**b**) A graph genome is indexed by creating a hash table with *k*-mers along all possible paths of the graph as keys and their corresponding graph genome positions as values. These *k*-mer positions can then be used as seeds for aligning sequencing reads against the graph. (**c**) Computational resource requirements of building, indexing and storing graph genomes. The resource usage statistics are provided for the standard linear reference genome (GRCh37), which was augmented with different subsets of the 1000G variants. All tests were performed using a single thread on the Amazon AWS instance type c4.8xlarge. (**d**) Runtime and memory usage for BWA and Graph Aligner using the global graph for 10 randomly selected samples from the Coriell cohort. Both BWA-MEM and Graph Aligner were executed using 36 threads on the Amazon AWS cloud instance type c4.8xlarge.

We implemented a suite of accompanying algorithms optimized for computational efficiency. We used the 1000G Phase 3, Simons Genome Diversity and other variant datasets to construct a graph genome reference that we refer to as the global graph in this manuscript (Supplementary Information, Supplementary Table 1, Supplementary Figs. 2-3). The global graph can be built and indexed in less than ten minutes in total (Fig. 1c). Such a graph reference can be stored in less than 30 gigabytes (GB) of memory or just over 1 GB of disk space (Fig. 1c, Supplementary Information). Loading a stored graph reference into memory takes less than two minutes. Over a tenfold range in the number of variants included, time, memory and disk storage consumption only grew around fourfold, twofold and 50%, respectively (Fig. 1c).

### Improved Read Mapping Accuracy using Graph Genomes

To support genomic analyses on our graph genome implementation, we developed a graph aligner for short reads that uses the *k*-mer index for seeding (Fig. 1b, Supplementary Fig. 5) followed by local read alignment against the graph (Supplementary Information). The read alignments against a path in the graph are projected to the standard reference genome and output to a standard BAM file, with the alignment path along the graph reported using custom annotation tags. Thus, the output format of our graph aligner maintains full compatibility with existing genomics data processing tools. When an unambiguous projection is not possible, for example for reads fully mapped within a long insertion variant, the reads are placed to the closest reference position, so that downstream analysis tools can access these reads conveniently.

We measured the graph aligner runtimes on 10 randomly selected high coverage whole-genome sequencing datasets (Supplementary Table 5). Read alignment against the global graph containing around 16 million variants (Supplementary Table 1) required around 4.5 hours per sample when using 36 threads, and was on average 9% shorter than that of BWA-MEM27 (Fig. 1d). This trend is reversed when only 8 or 12 threads are used, but the runtimes remain comparable (Supplementary Table 6). The peak RAM memory usage of the Graph Aligner was approximately 17.5 GB compared with an average of 21 GB of BWA-MEM (Supplementary Table 6). Overall mapping coverage between Graph Aligner and BWA-MEM are very similar, with small differences in low and high coverage regions (Supplementary Fig. 8).

In order to test the read mapping accuracy of the graph aligner, we simulated sequencing reads from individual sample VCF from the 1000G4 and the GiaB29 projects (Supplementary Information, Supplementary Figure 9). While reads without any variants are mapped equally accurately to the reference genome by the Graph Aligner and BWA-MEM, the Graph Aligner maintains a high mapping rate and accuracy even in reads containing long INDELs relative to the standard linear reference genome (Fig. 2, Supplementary Figs. 10-12). Less than 1% of the reads with >10bp insertions and deletions are mismapped using the 1000G graph, whereas this number is two and three times as large with BWA-MEM, respectively (Fig. 2). Even against a linear reference without variants, our graph aligner is able to align more reads containing INDELs than BWA-MEM (Fig. 2). In conclusion, our graph genome aligner significantly improves read alignment fidelity over BWA-MEM while maintaining comparable runtimes.

**Figure 2.**
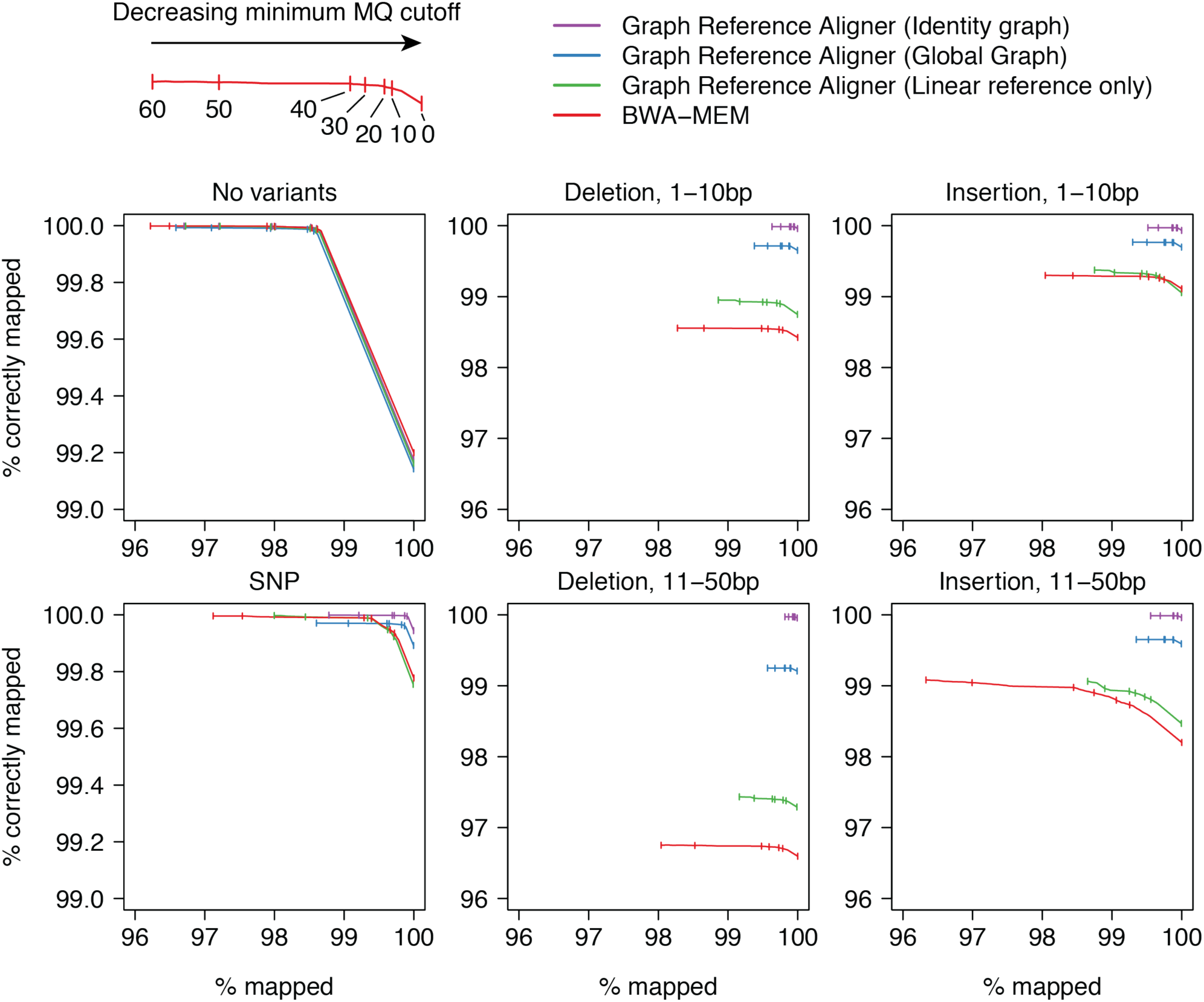
Read mapping accuracy using BWA-MEM and graph genomes. Simulated reads were divided into six categories based on the type of simulated variants they contain relative to the linear reference genome. The percentage of reads mapped correctly was plotted against the percentage of reads mapped for a range of mapping quality (MQ) cutoffs. ‘Identity graph’ refers to a graph genome containing only the genetic variants present in the respective target sample.

### Graph Genome Pipeline Improves Recall in Variant Detection

Graph Genome Pipeline (Supplementary Fig. 4) calls variants, including SVs, using a reassembly variant caller and variant call filters as suggested previously^28^ (Supplementary Information). Generating variant calls from raw FASTQs in the Coriell cohort (29-42x coverage) took on average 6h 19min (σ = 25min) using Graph Genome Pipeline on 36 CPU cores and 20 GB of memory. In comparison, the best practices GATK pipeline using GATK HaplotypeCaller (https://software.broadinstitute.org/gatk/best-practices/, hereafter referred to as BWA-GATK) executed on same hardware required 50 GB of memory and an average of 11h 30min (σ = 3h 16min) of runtime.

We devised four independent and complementary experiments to compare the variant calling accuracy of our Graph Genome Pipeline against that of two commonly used linear pipelines (BWA-GATK and BWA-Freebayes) as well as a recently published graph-based approach (Graphtyper). Furthermore, to separate the impact of the graph aligner and from the variant caller we also benchmarked GATK HaplotypeCaller results derived from Graph Aligner BAMs (Fig. 3b and Supplementary Table 10, which also presents results from BayesTyper). The first benchmarking experiment is based on sequencing data simulation, which provides a known ground truth for all variants throughout the genome, but likely incompletely reproduces the error modalities of real sequencing data. The second benchmarking experiment uses truth data established by the Genome in a Bottle Consortium (GiaB)^29^ for five high coverage whole-genome sequenced samples (50x coverage). These truth data cover only about 70% of the genome considered as “high-confidence” regions by GiaB, likely excluding the ∼30% of the genome that is hardest to align and call variants against. The third variant calling benchmark is based on measuring Mendelian consistency in family trios (Supplementary Figs. 16-18), which is an indirect proxy for variant calling accuracy, but can be conducted on real data throughout the genome. To support this approach, we developed computational methods to resolve variant representation differences in trio comparison and estimate the precision and recall rates of a variant caller using variant calls derived independently from each member of a family trio (Supplementary Information). Finally, we compared the variant calling results to SNP genotyping results from two commonly used SNP array platforms (Supplementary Information, Supplementary Fig. 19, Supplementary Table 8).

**Figure 3.**
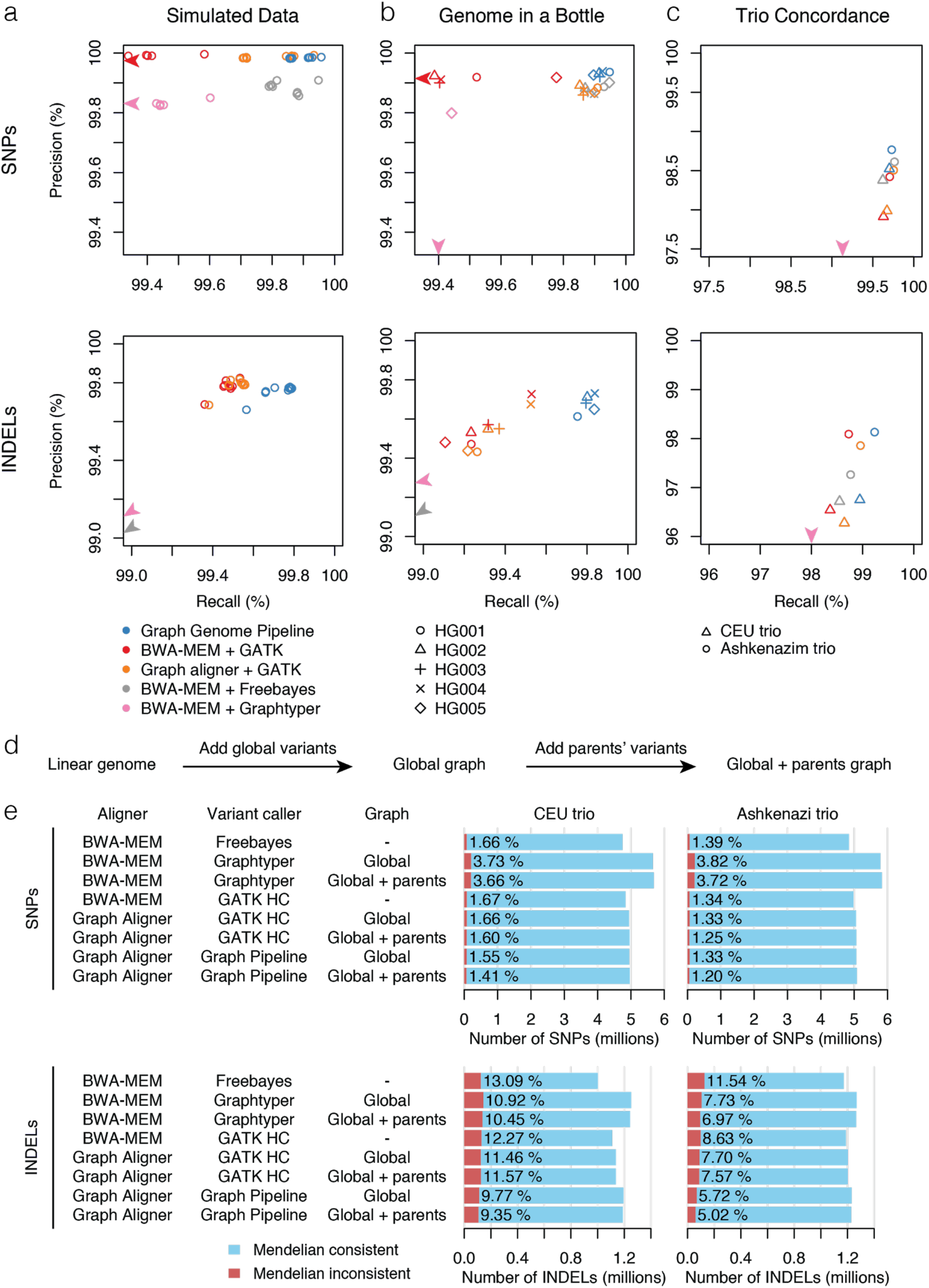
Variant calling benchmarking between Graph Genome Pipeline and BWA-GATK. (**a**) Precision and recall in simulated sequencing data. Every point corresponds to a different simulated individual. (**b**) Precision and recall benchmarking based on five samples for which the GiaB truth sets are available. We show BWA-GATK results with Hard Filtering as that approach performed better out of the two recommended options. Supplementary Table 10 shows BWA-GATK results with VQSR, as well as results from BayesTyper. (**c**) Statistically estimated precision and recall from the Mendelian consistency data derived from two family trios with independently called variants for each family member. (**d**) Schematic representation of the trio concordance analysis. (**e**) Mendelian concordance of two trios analyzed using BWA-GATK, Graph Genome Pipeline with the global graph and Graph Genome Pipeline using a global graph augmented by the variants detected in either parent of each family. The numbers shown are computed after resolving variant representation differences (Supplementary Information). Mendelian concordance rates without variant representation resolution are shown in Supplementary Figure 15 and Supplementary Table 13.

In addition to the standard approach of filtering the false positive variants that we mainly use in this paper, we have also developed a machine learning-based approach (Supplementary Information). While this method yields a significant improvement in the results and outperforms all other pipelines in the PrecisionFDA Truth contest (Supplementary Figs. 6 and 7, Supplementary Table 7), we do not use it as the main indicator of the Graph Genome Pipeline’s performance, as it relies on training using one of the GiaB samples, which are extensively used in most of our benchmarks.

A consistent pattern emerges from the benchmarking experiments. In both SNP and INDEL calling, Graph Genome Pipeline either has an equally good precision with better recall (Figure 3a,b, Supplementary Tables 9 and 10) or better precision with the same recall (Figure 3c, Supplementary Table 11) compared with other pipelines. Graph Genome Pipeline has the lowest rate of Mendelian violations while calls the second highest number of variants after Graphtyper (Fig. 3d,e, Supplementary Figure 15, Supplementary Tables 12 and 13), but Graphtyper has lower precision overall (Fig 3). The gain in SNP calling accuracy is driven by graph alignment, as SNPs called by GATK-HC from Graph Aligner BAMs have a recall similar to those by BWA-GATK (Fig. 3). Interestingly, Graph Genome Pipeline’s good performance in INDEL recall is driven primarily by our graph alignment-aware reassembling variant caller (Fig 3). Overall, Graph Genome Pipeline has a similar precision to BWA-GATK but improves recall by around 0.5%, corresponding to around 20,000 additional true variants being detected when extrapolated genomewide.

The GiaB variant call sets provide an estimate for the practical upper limit in achievable accuracy using the standard linear reference genome, since they are carefully curated from an extensive amount of high quality data generated from a combination of several different sequencing platforms, such as 300x PCR-free Illumina sequencing and high coverage 10x Genomics data, and meta-analyzed across a suite of state-of-the-art bioinformatics tools^29^. Interestingly, among the variants detect by Graph Genome Pipeline but asserted as homozygous reference by GiaB, a substantial proportion (26-42% across four samples, Supplementary Table 2) exhibited strong support from alternative sequencing technologies as well as in terms of Mendelian concordance (Supplementary Fig. 20). These variants are often located in variant-dense regions, half (52%) of them are part of the global graph, and most of the remaining variants (46%) are phased with one or multiple nearby variants present in the graph (Supplementary Fig. 20, Supplementary Table 3). Contrary to the linear reference genome, Graph Aligner is by design able to map reads across known variations without reference bias, which allows it to mitigate the impact of reference bias in repetitive or variant-dense genomic regions. We therefore hypothesized that these variants could be real but missed by all other linear reference genome-based pipelines used by GiaB due to reference bias. We successfully carried out Sanger sequencing at 351 and 598 of these “false FP” variants in two GiaB samples, HG001 and HG002, validating 63.6% and 60% of these variants as real variants missed by GiaB, respectively (Supplementary Tables 2 and 14). While these numbers constitute only 8.7% and 15.5% of the total number of FP calls made by our pipeline, they demonstrate that our graph genome implementation is able to overcome some practical accuracy limitations of linear reference approaches.

### A Unified Framework for SV Calling Using Graph Genome Pipeline

Sequence information of known SVs can be incorporated into a graph genome, allowing reads to be mapped across them. Graph Genome Pipeline is able to align reads across SVs while BWA-MEM fails to do so with both short Illumina reads (Fig. 4a and Supplementary Figs. 21-22) and long PacBio reads, even when PacBio reads are aligned using parameters tuned for PacBio data (Supplementary Fig. 22).

**Figure 4.**
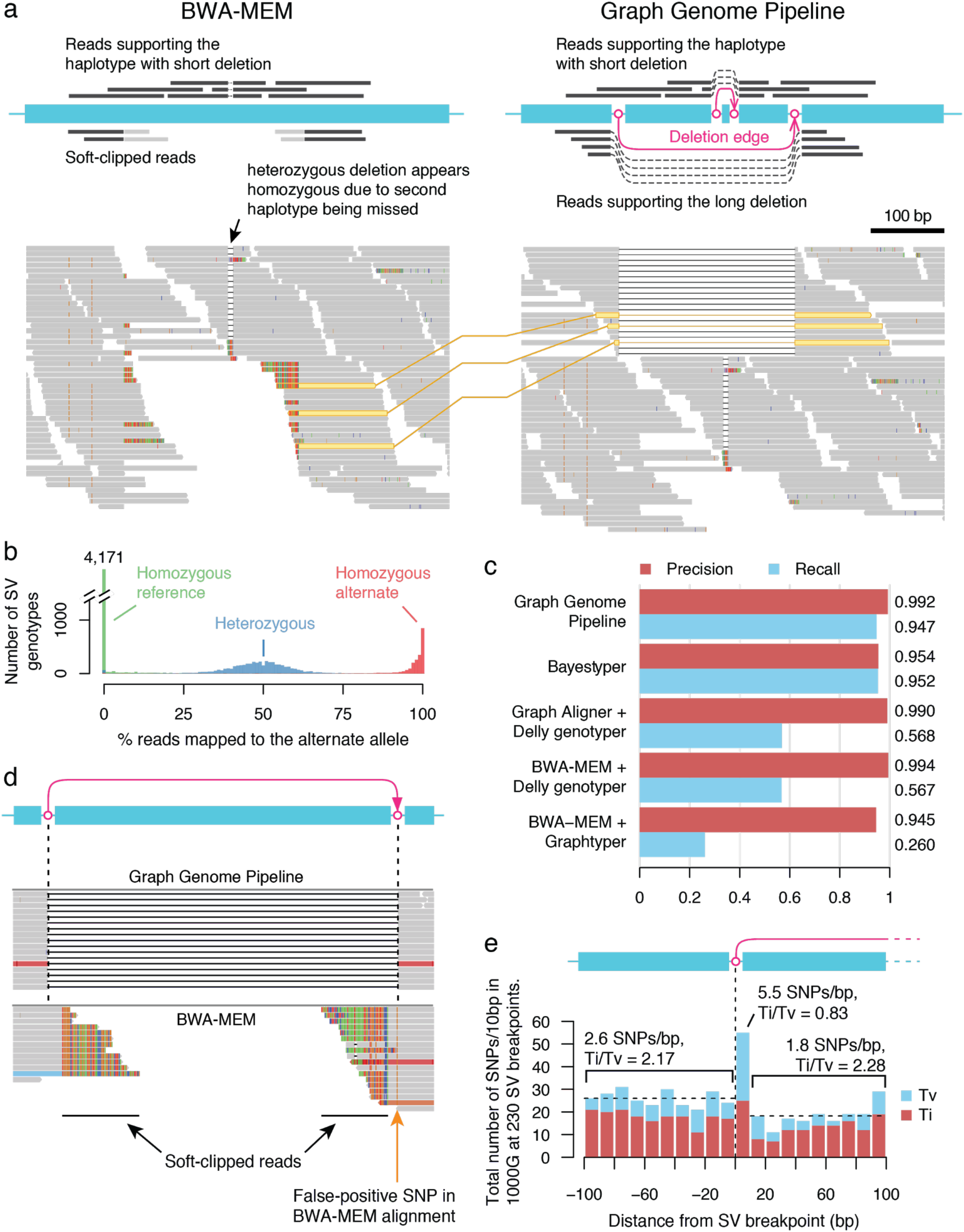
SV genotyping using graph genomes. (**a**) An example Integrative Genomics Viewer^41^ (IGV) screenshot of BWA-MEM and Graph Aligner alignments over an SV at chromosome 4 position 132,524,277-132,524,548 (GRCh38, Database of Genomic Variants ID: esv3602264), which in this sample (HG01628) overlaps with a small deletion variant. BWA-MEM is able to align reads across the small deletion, but not across the SV. Instead, reads on the SV haplotype become soft-clipped (colored vertical bars at the tips of the reads facing the SV). As a consequence of the SV not being called, the small intervening deletion appears homozygous in the read alignment. A schematic of the read alignment pattern is shown above the screenshot. Thanks to this SV being included in the graph genome, Graph Aligner correctly aligns reads across it, revealing the SV haplotype. (**b**) The fraction of reads aligning to an SV branch at an SV breakpoint stratified by the true genotype of each SV in each sample^5^, summarized across 230 SVs and 49 individuals. Reads aligning to the middle of an SV or reference branch are disregarded. (**c**) Benchmarking the graph genome pipeline against three other SV genotyping tools (BayesTyper, Graphtyper, Delly2) across 230 SVs and 49 individuals. BayesTyper and Graph Genome Pipeline use raw FASTQ as input, whereas Graphtyper and Delly2 require aligned reads. Within the curated 230 SVs, BayesTyper and Graph Genome Pipeline outperform Graphtyper and Delly2 on recall (Supplementary Information). All tools perform equally well in precision. (**d**) Example of an alignment that causes a false positive SNP due to misalignment against the linear reference genome. This sample has a homozygous deletion in this region, and Graph Aligner (top) aligns reads successfully across it. BWA-MEM (bottom) fails to align reads across the SV, but since the SV has 20bp of imperfect microhomology at the breakpoint, BWA-MEM aligns the reads on the right hand side over the imperfect microhomology region, resulting in a spurious homozygous G>A SNP call. (**e**) Total number of transition and transversion SNPs reported by 1000G around the breakpoints, aggregated over the 230 SVs and grouped by distance from an SV breakpoint in 10bp bins. I.e. 2.6 SNPs/bp indicates that there are a total of 2.6 SNPs per bp within 100bp upstream of each breakpoint over all 230 SVs. Positive distances (i.e. >0bp) are within a SV, while negative distances (i.e. <0bp) are outside an SV. For example, positions +1 to +10bp correspond to the 10bp closest to an SV breakpoint within a deletion, while positions −1 to −10 bp correspond to the 10bp closest to an SV breakpoint outside a deletion.

To demonstrate that reads spanning SV breakpoints can be used to directly genotype SVs, we manually curated a dataset of 230 high-quality, breakpoint-resolved deletion-type SVs (Supplementary Table 15 and Supplementary Information) and genotyped them across 49 individuals from the Coriell cohort, for which the true SV genotypes are available from the 1000 Genomes Project^5^. Although our SV set does not include any events composed purely of inserted sequence, many of them involve novel sequence insertions at their breakpoints (Supplementary Fig. 3). The fractions of reads spanning SV breakpoints segregates cleanly into three clusters based on SV genotype (Fig. 4b) suggesting that graph genome-based SV genotyping could be accomplished even with a simple read counting-based method (Supplementary Information, Supplementary Table 4). Based on these reads, the Graph Genome Pipeline reassembles SVs (alongside SNPs and INDELs) with an SV genotyping accuracy comparable to current SV callers (Fig 4c, Supplementary Table 4). SV genotyping performance within the 230 SVs is on par with BayesTyper, and although both Delly and Graphtyper have a very high precision, they suffers from low recall (Fig 4c, Supplementary Table 4). However, we also acknowledge that the SV set presented here is devoid of more complex variants such as mobile elements and inversions, and Graph Genome Pipeline’s performance regarding complex SVs is yet to be fully tested.

To compare the SV genotyping performance of the graph aligner to competing technologies, we focused on the GiaB sample HG002, for which both Illumina read and PacBio long read data are publicly available. We manually examined the alignment results from each technology in the 230 SVs in this sample by the graph aligner. BWA-MEM is unable to align Illumina reads across any of these SVs (Fig. 4a, Supplementary Figs. 21 and 22). Similarly, PacBio long read alignment fails in all these SVs, even when aligned with BWA-MEM using parameters tuned for PacBio data (Supplementary Fig. 22). Thus, graph genomes could potentially improve SV genotyping for short read and long read sequencing technologies alike.

Two events among the 230 curated SVs, esv3642033 and esv3638126, are in strong linkage disequilibrium with SNPs significantly associated (P-value < 5×10^−8^) with breast cancer (rs1436904) and obesity class II risk (rs11639988), respectively^30,31^ (Supplementary Fig. 22). We were able to correctly genotype the presence of these two SVs in 48/49 and 49/49 samples, respectively. Thus, graph genome technology may enable integrated methods for genotyping of common SVs, including those with clinical and biological relevance.

### Graph Genomes Prevent Erroneous Variant Calls Around SVs

Structural variations mediated by certain DNA repair mechanisms can exhibit microhomology, including imperfect microhomology, around their breakpoints^32^. If an aligner is not aware of an SV, sequencing reads spanning the SV could become erroneously aligned over a region of imperfect microhomology instead, causing mismatches over the region to be spuriously reported as SNPs and INDELs (Fig. 4d). To quantify this effect in 1000G, we compared the rate of 1000G SNPs around the SV breakpoints with the background rate of SNPs in 1000G (Fig. 4e). These metrics are computed in aggregate over all 230 SVs. Within the deleted portions of the 230 curated deletion SVs combined, the aggregate rate of 1000G SNPs is 1.8/base pair (bp), and these SNPs have a transition-transversion ratio (Ti/Tv) of 2.28, which is expected from real biological SNPs33 (Fig. 4e). In contrast, in the ten first base pairs immediately after an SV breakpoint, the aggregate 1000G SNP rate is increased 3-fold to 5.5/bp, suggesting that 67% of 1000G SNPs called within 10bp of an SV breakpoint are in fact false (Fig. 4d,e). The Ti/Tv of these SNPs is 0.83, deviating significantly from the expected ratio of 2.1. Assuming that spurious SNPs have an expected Ti/Tv of 0.5, the FP SNP rate over this region estimated using Ti/Tv is 79%, reaching a similar value to that estimated using total SNP counts (Supplementary Information). In addition to FP variant calls, we also encountered examples where variants overlapping with an SV erroneously appear homozygous, because BWA-MEM fails to align reads across the SV and thus fails to detect the corresponding SV haplotype (Fig. 4a). Thus, using population variation information in graph genomes can mitigate variant calling and genotyping errors around SVs.

### Incremental Improvement in Variant Calling Recall through Iterative Graph Augmentation

As common genetic variants continue to be cataloged across populations, newly discovered variants can be incrementally added to existing graph genomes to increase the comprehensiveness of the graph while maintaining backward compatibility to samples analyzed using earlier versions of the graph (Fig. 1a). To test whether incremental graph augmentation would improve variant calling, we augmented the global graph with variants detected in ten samples from three super-populations of the Coriell cohort as well as a Qatar genome project cohort^14^ (Fig. 5a,b, Supplementary Table 16), and compared the variant calls obtained using the global and the four augmented global graphs. Augmenting the global graph genome increases the number of known variants (i.e. present in dbSNP) discovered slightly (5% and 10% median increase known SNPs and INDELs discovered, respectively; Fig. 5c,d, Supplementary Table 16). However, the augmented graphs result in almost twice the number of novel (i.e. those not in dbSNP) SNPs and INDELs being called, respectively (Fig. 5c,d, Supplementary Table 16).

**Figure 5.**
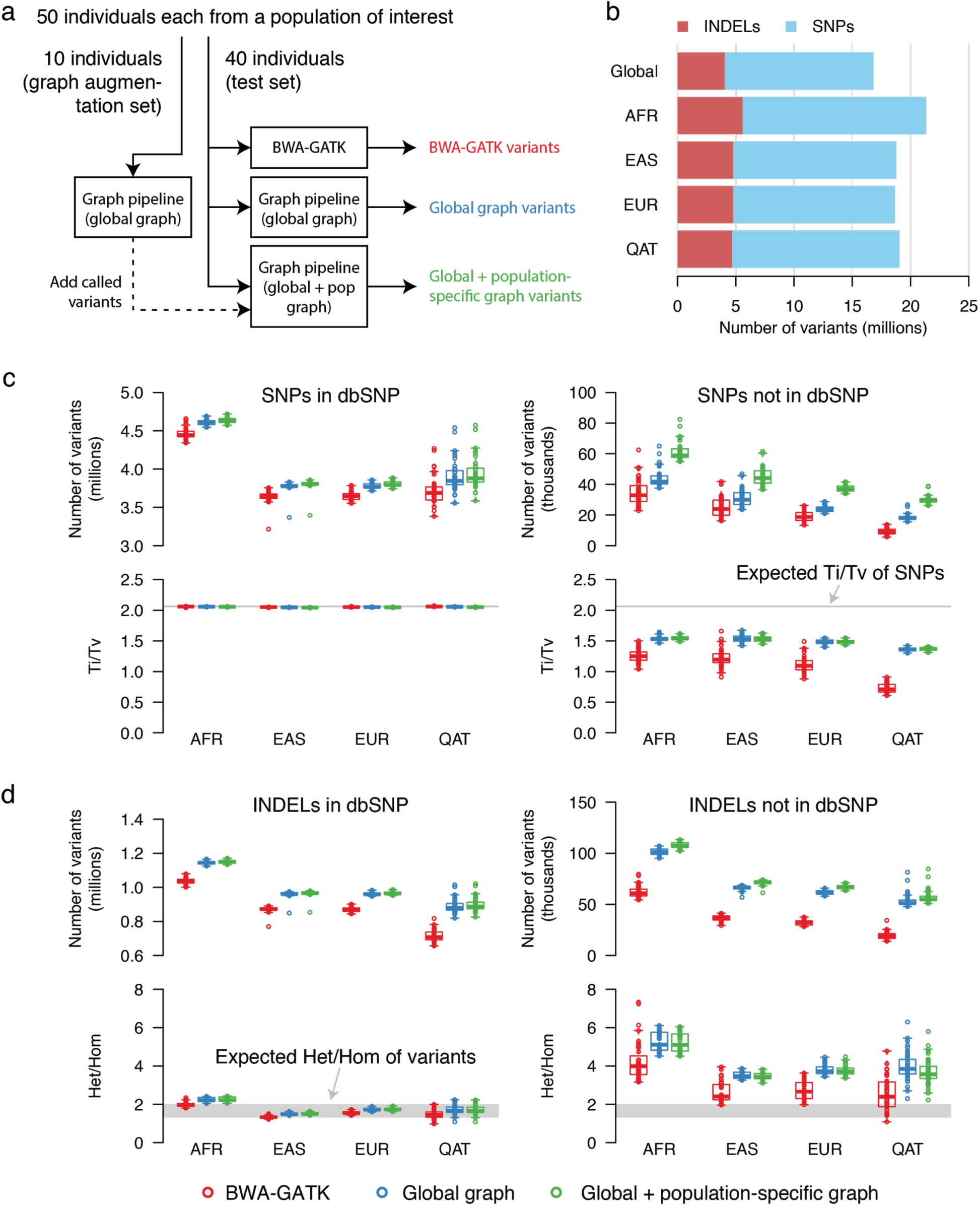
The effect of iteratively augmented graph genomes on variant calling. (**a**) Schematic representation of the graphs generated in the graph augmentation experiment. (**b**) Numbers of SNPs, INDELs and other variant types in the global and augmented graphs. (**c**) Counts and transition/transversion ratios of known and novel SNPs called through the BWA-GATK, global graph and augmented graph pipelines. Grey horizontal lines indicate the expected transition/transversion genome-wide ratio^33^. In box plots, center line, box limits and whiskers indicate the median, upper and lower quartiles and 1.5x interquartile range, respectively. (**d**) Counts and het/hom ratios of known and novel INDELs called through the BWA-GATK, global graph and population-augmented graph pipelines. Horizontal bars indicate the average value of each group. Gray rectangles indicate the expected range of population averages for heterozygous/homozygous alternate ratios^33^. Box plot elements are defined as in (**c**).

We measured the quality of the detected variants indirectly using Ti/Tv and heterozygous-to-homozygous alternate allele ratio (het/hom) for SNPs, and INDELs, respectively^33^. For each tested pipeline, known SNPs and INDELs have a Ti/Tv ratio of 2.04-2.06 and het/hom ratio of 1.3-2.0, respectively (except for INDELs called by the graph pipeline in the African population). These values are within the expected range for common variants33. In contrast, these metrics fall outside the expected range for novel SNPs and INDELs. The Ti/Tv ratios of novel SNPs are closer to the expected Ti/Tv when called by Graph Genome Pipeline compared with BWA-GATK, whereas the ordering is reversed for the het/hom ratios of novel INDELs (Fig. 5c-d). Importantly, while graph augmentation results in more known and novel variants being called, their Ti/Tv and het/hom ratios remain unaffected. Therefore, incremental graph augmentation may improve variant recall with little impact to the quality of of the variants detected.

Graph augmentation had a similar impact in the read alignment and variant calling experiments. Read alignment recall reaches almost 100%, if a target sample is aligned against a graph genome that contains all its actual variants (Fig. 2). Likewise, trio concordance and variant calling recall is further improved, if variant calling in a child is performed using a graph genome augmented with variants detected in the respective parents (Fig. 3c-e, Supplementary Tables 11-13). Thus, incremental augmentation of graph genome references yields cumulative improvements in variant calling recall without an accompanying decrease in precision.

## Discussion

Here we presented, to our knowledge, the first computational infrastructure for building, augmenting, storing and querying multi-genome references at comparable computational requirements to prevailing linear reference-based methods. Our graph genomes are constructed from the linear reference genome and genetic variants provided in standard FASTA and VCF formats, and our graph genome aligner outputs read alignments in the standard BAM format against the coordinate system of the linear reference genome. As a result of the backward compatibility and computational efficiency of our graph genome infrastructure, any existing genomics pipeline can swap in our graph genome algorithms to capitalize on the accompanying improvements in accuracy without requiring any other pipeline or hardware upgrade. This is demonstrated by GATK HaplotypeCaller’s improvement in variant calling accuracy when Graph Aligner, as opposed to BWA-MEM, is used for read alignment.

Our benchmarking experiments demonstrate that using a graph genome reference improves read mapping and variant calling recall, including that of SVs, without a concomitant loss in precision (Figs. 2-5). Our graph aligner is able to readily align reads across breakpoint-resolved SVs included in the graph, unlike linear reference genome-based methods even with long reads (Fig. 4a and Supplementary Fig. 21 and 22). Direct genotyping of SVs using these reads is possible in some cases (Supplementary Table 4). In contrast, existing methods for identifying and genotyping SVs require specifically designed multi-step algorithms^9^. Graph Genome Pipeline allows the identification and genotyping of SVs and small variants in a single unified process, raising the prospect for population-scale SV genetics and association studies.

Currently Graph Genome Pipeline analyzes samples individually, and a version that performs joint variant calling28,34 is under development. Further improvements to variant calling could be achieved by representing haplotypes of small variants and SVs as paths in the graph genome. Information such as allele frequencies of each variant and linkage disequilibrium between them could be incorporated into the graph, providing additional statistical information for read alignment and variant calling. This approach could provide a computationally efficient alternative to joint variant calling in leveraging previously accumulated population genetics information when analyzing a newly sequenced sample. Future efforts from us and others^35^ will be required to assess the benefits of using graph genomes with encoded allele frequency and linkage disequilibrium information.

The potential benefits of an unbiased multi-genome graph reference are not limited to variant calling, but cover the full range of genomics research. Graph genomes provide an unbiased, representative scaffold for read alignment, which is critical to sequencing alignment quantification applications such as RNA-sequencing analysis, ChIP-seq analysis and CNV calling. Our graph genome implementation can also be used to encode information other than whole human genomes. Individual gene families or related microbial strains could be compressed and efficiently searched using our graph genome algorithms. Similarly, the transcriptome could be represented as genomic deletions, allowing RNA-seq reads to be directly aligned across exon-exon junctions^36^. A personalized graph genome could be constructed from a sequenced germline genome, in order to provide an optimized scaffold for somatic variant detection in matched cancer genomes.

The ability to genotype variants, including SVs, efficiently and accurately opens the door for myriad clinical and research applications. A graph built from rearranged B or T cell receptors could allow quantitative tracking of mature B or T cells populations. A somatic graph incorporating somatic mutations and SVs of a treated primary tumor could enable sensitive monitoring of tumor relapse at the primary site or in cell-free DNA. A “precision medicine graph” could be designed and continually updated to include all actionable genetic variants in order to support clinical decision making.

As more genetic variants are accumulated on a reference graph genome, cumulative accuracy gains in genomics analysis can be achieved. This is consistent with recent efforts to establish population-specific reference panels, which have been shown to contribute to increased accuracy in imputation^37–39^ and genetic risk prediction^40^. The current wave of national sequencing projects will extend the catalogs of population-specific genetic variants, which will incrementally improve the prospects for graph genome reference approaches^22,25,26^, including ours. Completion of the first draft of the human reference genome marked the beginning of human genomics. Our computationally efficient and flexible graph genome implementation supports the community for a gradual transition towards a graph-based reference system.

### Data availability

Raw sequencing data for the 150 Coriell WGS samples (Figs. 1, 4 and 5) can be accessed European Nucleotide Archive, Study Accession PRJEB20654, (https://www.ebi.ac.uk/ena/data/view/PRJEB20654). Raw sequencing data for the Qatari samples (Fig. 5) used can be found in the NCBI SRA accessions SRP060765, SRP061943 and SRP061463 (https://www.ncbi.nlm.nih.gov/Traces/study/?acc=SRP060765%2CSRP061943%2CSRP061463&go=go). Genome in a Bottle data (Fig. 3) is available from the NCBI FTP site: ftp://ftp-trace.ncbi.nlm.nih.gov/giab/ftp/data. The Sanger sequencing traces will be deposited to NCBI Trace Archive (Accession ID pending).

### Code availability

Graph Genome Pipeline is available for use on the Seven Bridges Cloud Platform: https://www.sevenbridges.com. Binaries are available upon request. Both options require a license agreement.

## Acknowledgements

We are grateful for the members of the GA4GH Data Workgroup, Benchmarking, and Reference variation initiatives, in particular Dr. Justin Zook, for insightful discussions and ideas. Dr. Maxime Huvet helped refine the treatment and presentation of ideas behind trio-based benchmarking.

Research reported in this publication was supported in part by UK Department of Health grant SBRI Genomics Competition: Enabling Technologies for Genomic Sequence Data Analysis and Interpretation administered by Genomics England.

## 1. Reference Effects in 1000 Genomes Variants

Supplementary Figure 1 shows the frequency of variants that are either fixed or where the non-reference allele is a major allele in each 1000G population.

**Supplementary Figure 1.**
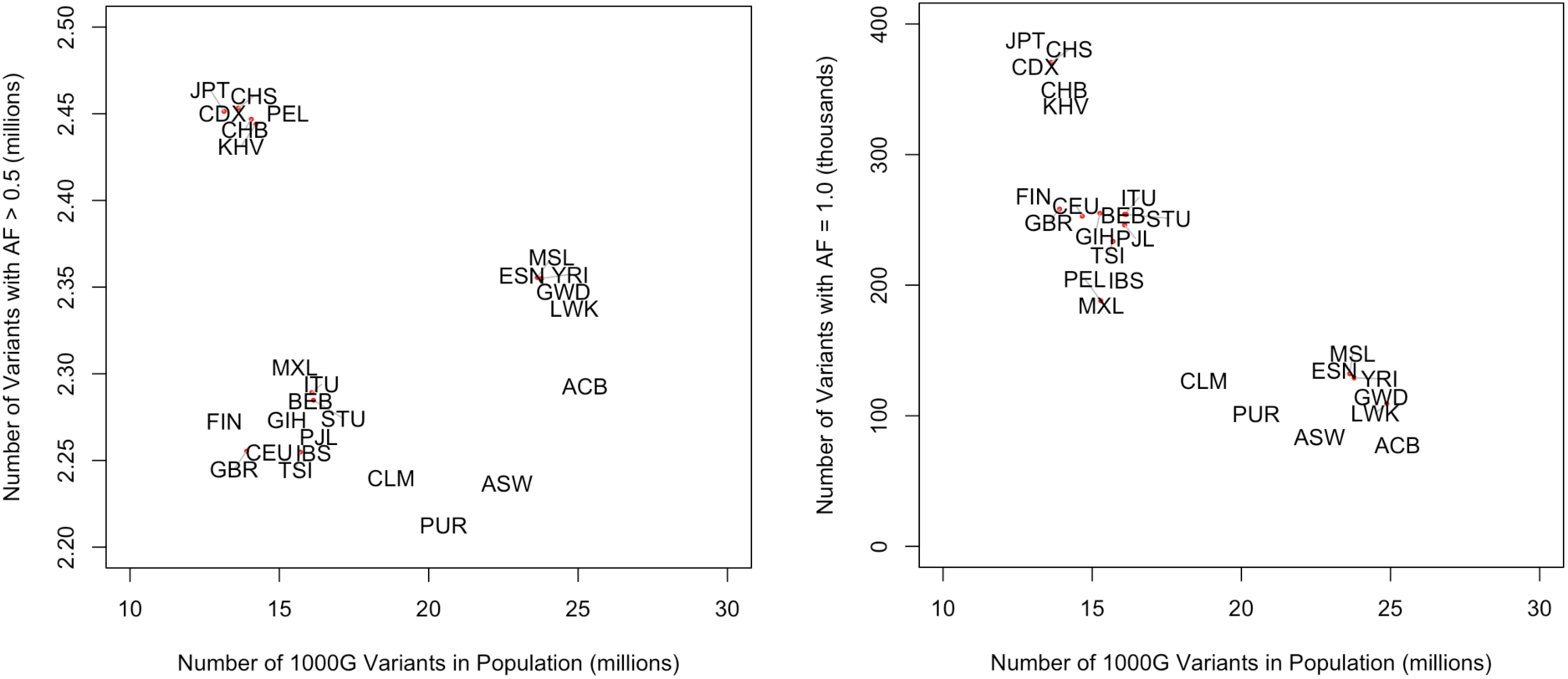
The frequency of major allele (i.e. within-population allele frequency > 0.5; left) and fully fixed variants (right) in each 1000G population. The population labels are as follows. CHB: Han Chinese in Beijing, China; JPT: Japanese in Tokyo, Japan; CHS: Southern Han Chinese; CDX: Chinese Dai in Xishuangbanna, China; KHV: Kinh in Ho Chi Minh City, Vietnam; CEU: Utah Residents (CEPH) with Northern and Western European Ancestry; TSI: Toscani in Italia; FIN: Finnish in Finland; GBR: British in England and Scotland; IBS: Iberian Population in Spain; YRI: Yoruba in Ibadan, Nigeria; LWK: Luhya in Webuye, Kenya; GWD: Gambian in Western Divisions in the Gambia; MSL: Mende in Sierra Leone; ESN: Esan in Nigeria; ASW: Americans of African Ancestry in SW USA; ACB: African Caribbeans in Barbados; MXL: Mexican Ancestry from Los Angeles USA; PUR: Puerto Ricans from Puerto Rico; CLM: Colombians from Medellin, Colombia; PEL: Peruvians from Lima, Peru; GIH: Gujarati Indian from Houston, Texas; PJL: Punjabi from Lahore, Pakistan; BEB: Bengali from Bangladesh; STU: Sri Lankan Tamil from the UK; ITU: Indian Telugu from the UK.

## 2. Graph References

### 2.1. Global Graph Reference

In most of the experiments in this paper we make use of a Global Graph Reference, a graph reference we built from the GRCh37 linear reference genome^16^ and known variants from several publicly available sources (Supplementary Table 1). All variants were passed through a representation normalization step using Bcftools norm (http://samtools.github.io/bcftools) prior to being merged into a single set. The data sources used to build the Global Graph Reference are:

- 1000 Genomes Phase 3 variants^4^, with alternate allele frequency greater than 0.01.
- Simons Genome Diversity Project variants, reprocessed and joint-called using the BWA-GATK Best Practices workflow^17^ and filtered to require an alternate allele occurrence of 10 or greater
- Mills INDELs database^42^
- 1000 Genomes Phase 1 INDEL calls^43^
- A set of 704 long deletion events from the 1000 Genomes Phase 3 data^5^ that have precise breakpoints defined

We set fairly high allelic frequency thresholds for variants that were included in the graph, as we observed a small loss of accuracy when using low frequency variants (Supplementary Table 10). We believe the issue to be caused by the higher false positive rates of lower frequency variants, which has been noted previously44. Furthermore, low allele frequency variants tend to have a lower rate of reproducibility in 1000 Genomes Phase 3 data4, as well as the fact that the 1000 Genomes variant filtering strategy had been modified to be more lenient when filtering out low frequency variants4. The distribution of INDEL sizes in the global graph is shown in Supplementary Figure 2.

All SNPs that fall within short tandem repeat regions are excluded, because such calls often arise from mismapping of INDEL variants or sequencing errors. This list of ≤50bp tandem repeats was compiled by the GA4GH Benchmarking working group^45^. Finally, we applied the following approach to remove variants that are consistently generated through read mapping problems. We simulated 30x coverage 150bp error-free paired-end sequencing data from the standard linear reference genome. We aligned these reads to the initial Global Graph Reference, and called variants. This method yielded 2,026 SNPs and 72 indels that were then excluded from the graph to obtain the final build of the Global Graph Reference. Many of these calls are likely caused by systematic mapping errors in some of the data sources used to build the graph. GRCh37 was chosen over the more recent GRCh38 build primarily because there are more sources of genetic variation data available for the former. The 1000G sample NA12878 was excluded from the data prior to computing the 1000G alternate allele frequencies, since this sample is used in some of the benchmarks in this paper (including GiaB and trio consistency).

**Supplementary Table 1.** Variant counts in the Global Graph reference by category.

| | SNP | Insertion<br>$\leq 30$bp | Deletion $\leq 30$bp | Insertion $> 30$bp | Deletion $> 30$bp |
| --- | --- | --- | --- | --- | --- |
| Count | 12,740,241 | 1,867,108 | 2,138,949 | 22,051 | 28,106 |

**Supplementary Figure 2.**
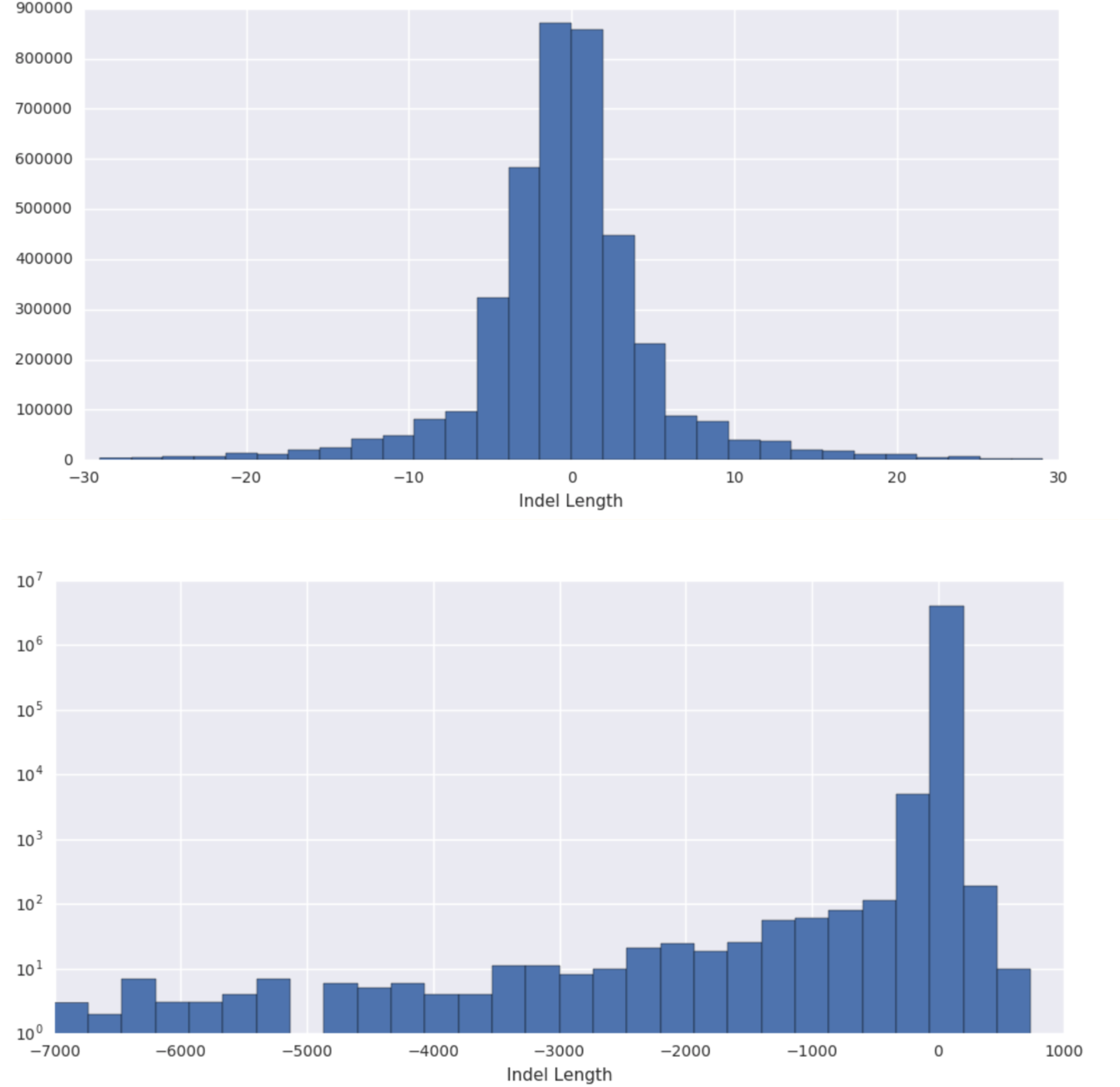
Size distribution of indels in the Global Graph reference for indels shorter than 30bp (top) and longer than 30bp (bottom).

A subset comprising 230 out of the 704 large events (Supplementary Table 15) was carefully curated and used for the SV genotyping experiments described in the paper. The aim was to establish a truth set based on a subset of the 1000 Genomes SV data^5^ with SVs that are most consistent across the literature and can be reassembled using an independent dataset (Simons Genome Diversity Project^17^). The 230 SVs were selected through intensive filtering of the 1000 Genomes SVs against the Database of Genomic Variants’ (DGV)46 Gold Standard (GS) set. This set only includes “loss” and “gain” events and are devoid of more complex variants such as inversions and mobile elements. From the DGV GS set, we only kept SV events supported by at least 2 different studies, 2 different platforms, 2 different algorithms, 500 unique samples (with at least 100 Europeans) and 10% frequency. This left us with 2,133 events out of 114,555 events in the DGV GS set (March 2016). We then intersected the outer range of these 2,133 SVs with SVs reported by the 1000 Genomes SVs that are common (>5% frequency in EUR), in Hardy-Weinberg equilibrium within EUR (Bonferroni corrected P < 0.05), and which have at least one tagging SNP (r2 > 0.9). This led to a set of 298 putative common EUR SVs with at least one tag-SNP. Then, for each SV region, reads were sampled from individuals from the Simons Genome Diversity Project^17^ with tag-SNPs suggesting a homozygous SV and reassembled using TIGRA^47^ per individual. Where available, the Icelandic non-repetitive, non-reference sequences reassembled by PopIns6 were used rather than TIGRA-reassembled contigs. SVs within the HLA region were excluded. These reassembled contigs were then remapped to the reference using a custom algorithm described in Supplementary Information 2.1.1 and small variants 10bp around SV breakpoints were excluded. In the end, 230 reassembled contigs were successfully remapped to unique positions on the reference (Supplementary Figure 3). Each event in the SV set had more deleted than inserted sequence, although many of them did contain at least some novel sequence insertion as part of them (Supplementary Figure 3).

Although the global graph reference is intended to be used as the default graph for any human study, we also provide further evidence that augmented graphs tailored for each study, such as the population graph reference and the pedigree graph reference (Supplementary Information 2.2-2.3), can improve genome inference.

**Supplementary Figure 3.**
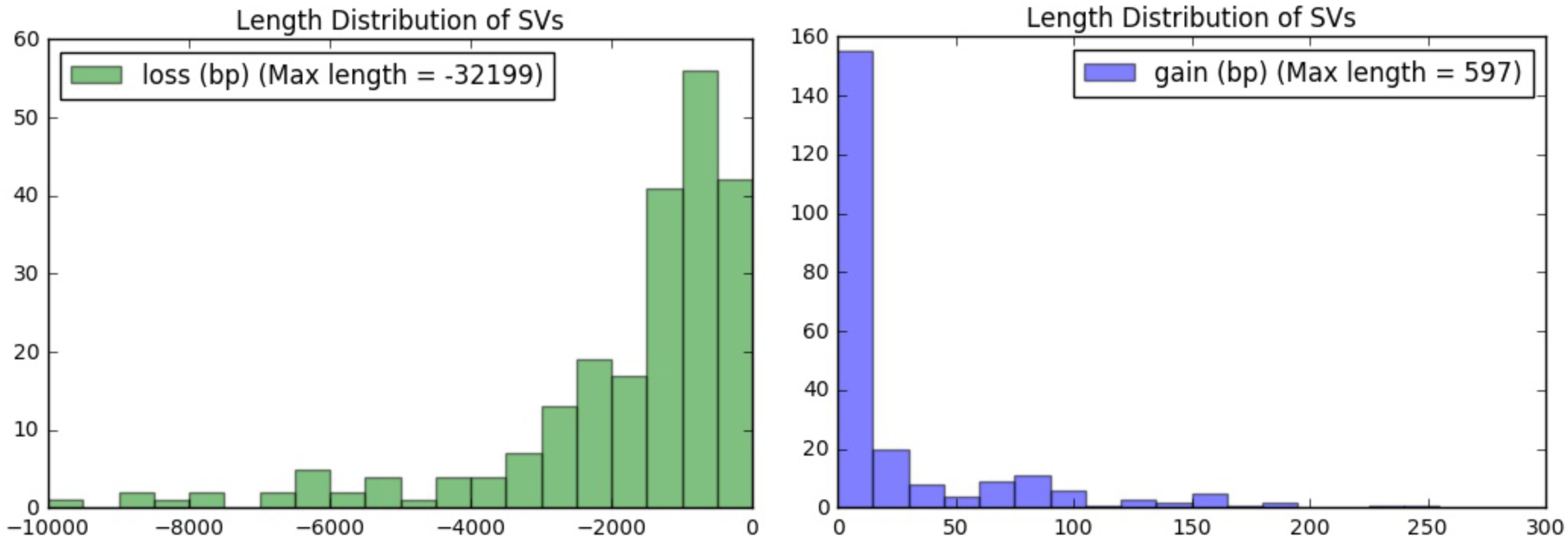
Size distribution of the 230 large deletion events used to build the SV graph. All events are classified as deletions, ranging between 50 and 32,199 bases in length (left). 120 events also feature an insertion of a shorter sequence up to 597 bases in length (right).

#### 2.1.1. Representing SVs in the graph and defining SV breakpoints

Assembled SV contigs were positioned to the graph reference genome using a custom algorithm that addresses challenges unique to graph alignment that current tools like BLAST, BLAT, and AGE do not handle. In particular, we needed to ensure that reads which map perfectly to the assembled contigs still perfectly map in the graph, while being robust against homologies around the breakpoints and errors in the assembled contigs. The main feature is the exact SV branch added to a graph reference is optimized for the intended target sequencing read length.

Using a BED file of the reassembly regions, the assembled contigs from multiple samples for each region, and the target read length as inputs, the algorithm outputs a VCF file for the SVs. This is done through the five following steps:

Establish local reference sequence: for each reassembly region, take the linear reference with 2,000 bp flanks on both sides as the local reference sequence.

Establish initial flanks: for each sample, map its corresponding assembled contig to the local reference sequence using BLAST. Check if the contig splits into two (possibly overlapping but not contained) parts, each with a good gapped alignment to distinct parts of the local reference sequence. If no, then the contig is rejected because either the contig is roughly identical to reference sequence or the SV is too complex. If yes, the two alignments establishes the initial 5’- and 3’-end flanking regions.

Resolve homology and find the optimal flanks: in both initial flanking regions, find the maximal exact match longer than the target read length which is the closest to the center (towards 3’-end for 5’-end flank and vice versa). Check if the 5’- and 3’-end exact matches overlap either in the contigs (deletion with breakpoint homology) or in the reference (insertion with breakpoint homology). If yes, remove the exact overlapped parts in the 3’-end. This establishes the optimal flanks.

Single-sample call: compare the contig with the local reference sequence between the optimal flanks and generate the VCF data for this sample in this region.

Global consensus: for each region, output the VCF with variant position and alleles determined through majority vote over all samples.

Graph representation of SVs were defined by setting the target read length to 100bp.

#### 2.1.2. Genome Graph Containing Low Frequency Variants

Our global graph has a relatively high minimum allele frequency cut-off (1%) on the variants included from the 1000 Genome and ∼2% (at least 10 alternate alleles genotyped) on the variants included from the Simons Genome Diversity datasets. We have also explored the possibility of building a graph using low frequency variants. For this purpose we constructed the graph using the same data sources as the above described global graph, but using 1000 Genomes Phase 3 variants with a minimum allele frequency threshold of 0.1% and Simons Genome Diversity variants with the minimum alternate allele occurrence threshold of 3 (out of the 280 samples in the Simons dataset). From this graph we excluded variants from regions of very high variation density (≥50 variant positions within a 200bp window), which are typically associated with short tandem repeats, as such regions introduced a significant deterioration in alignment speed. The final low allele frequency graph contains 35,444,012 variants.

Comparing the results from the benchmarks on the 5 GiaB we noticed a slight drop in variant calling accuracy using the low allele frequency graph (Supplementary Table 10). We believe this to be associated with the increased false positive rates amongst the lower frequency variants4,44, which may introduce alignment artefacts if added to the graph. Alignment against the low frequency graph was about 15% slower on average (mean 15.7%, std 4% measured on the 5 GiaB samples, Supplementary Table 17).

### 2.2. Population Graph Reference

We constructed the population graph references by first identifying variants in 10 individuals drawn from each population using our Global Graph Reference-based pipeline (see Supplementary Information section 3). From the detected variants, we removed those that were preexisting in the Global Graph Reference, and added the remaining novel variants to the Global Graph Reference (Figure 5a in the main text, Supplementary Table 16). This augmented Graph Reference is used to analyse further samples from the same population (Figure 5a in the main text, Supplementary Table 16).

### 2.3. Pedigree Graph Reference

The Pedigree Graph Reference of a family is built by calling the parent variants using the Global Graph Reference-based Graph Genome Pipeline and adding them to the Global Graph Reference. This Pedigree Graph Reference is presumed to contain the inherited variants as well as a very small number (∼100) of de novo mutations in the child. It is subsequently used in trio-based variant calling benchmarking.

## 3. Graph Reference Based Genome Determination Pipeline

The analysis steps in the Graph Genome Pipeline are shown in Supplementary Figure 4.

**Supplementary Figure 4.**
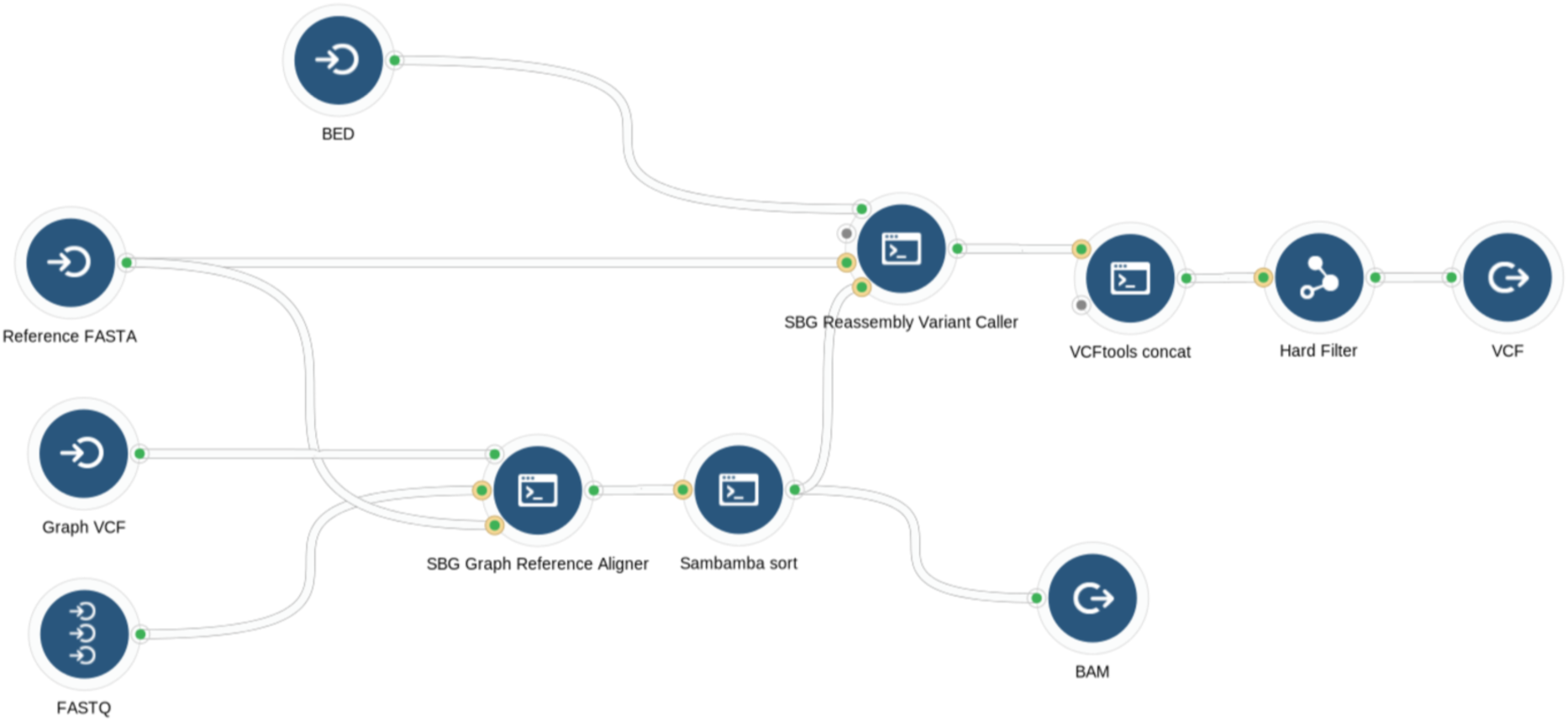
Schematic of the pipeline incorporating the Graph Reference Genome-based aligner and variant caller.

### 3.1. Graph Aligner

#### 3.1.1. Graph representation

We use adjacency lists to store the set of connected vertices for each vertex in the graph. Each vertex stores references to the set of incoming and outgoing edges. The sequence data is stored separately in a buffer; edges hence need only contain an offset address in the buffer and length and so inserting a new edge (which involves splitting an existing edge) is a fast operation.

The graph is efficiently serialised by handling the sequence data and graph structure separately. The sequence data is compressed in blocks using an n-bit encoding, where n is determined according to the alphabet size of the data in the block; n=2 for a sequence containing {A,C,T,G}. The structure of the graph is serialised by storing a reference to the sequence represented by the i-th edge along with the start and end loci at which the edge was inserted into the graph containing (i-1) edges; in other words the set of graph manipulations are stored in the same order as they were used to generate the graph. The index is not serialized.

Given these approaches, the graph containing the complete 1000G variant set occupies 23.8GB of RAM, comprising of 13.3 GB for the graph structure, 3 GB for the sequence data and 7.5 GB for the index, and (when serialised) 1.1 GB of disk space, of which 63% is used for sequence data and the remainder for the graph structure.

#### 3.1.2. Global Search

A hash-based search index is used for efficiently finding reads in a graph reference. A hash value is calculated for nucleotide sequences of length *k* (*k*-mer) in the graph reference. Each search index entry contains a list of start positions of *k*-mers (*k*-mer loci) that have the same hash value. The *k*-mers are determined by sequentially traversing the graph and indexing sequences of length *k* starting at every *s*-th position, where s is the indexing step and 1≤ *s* ≤ *k* (Supplementary Figure 5). Variant edges are similarly indexed at every s-th position until the k-mer would start on the parent edge. For the purposes of indexing, the position on an alternate branch is counted starting from the beginning of the graph and taking the reference path in all branching points preceding the variant branch(es) of interest (Supplementary Figure 5). A *k*-mer is limited to follow up to 16 edges in the graph, where the limit of 16 follows from the observation that such regions of the graph act as attractors. Entries in the index that contain more than a certain number of loci (hash list size threshold) are removed to prioritize seeding based upon informative *k*-mers (i.e. *k*-mers containing less common nucleotide sequences), speeding up the search and reducing memory requirements. The results presented in this paper were calculated for *k*=21 and *s*=7. This combination of settings was found to deliver a good trade-off between index size and performance.

**Supplementary Figure 5.**
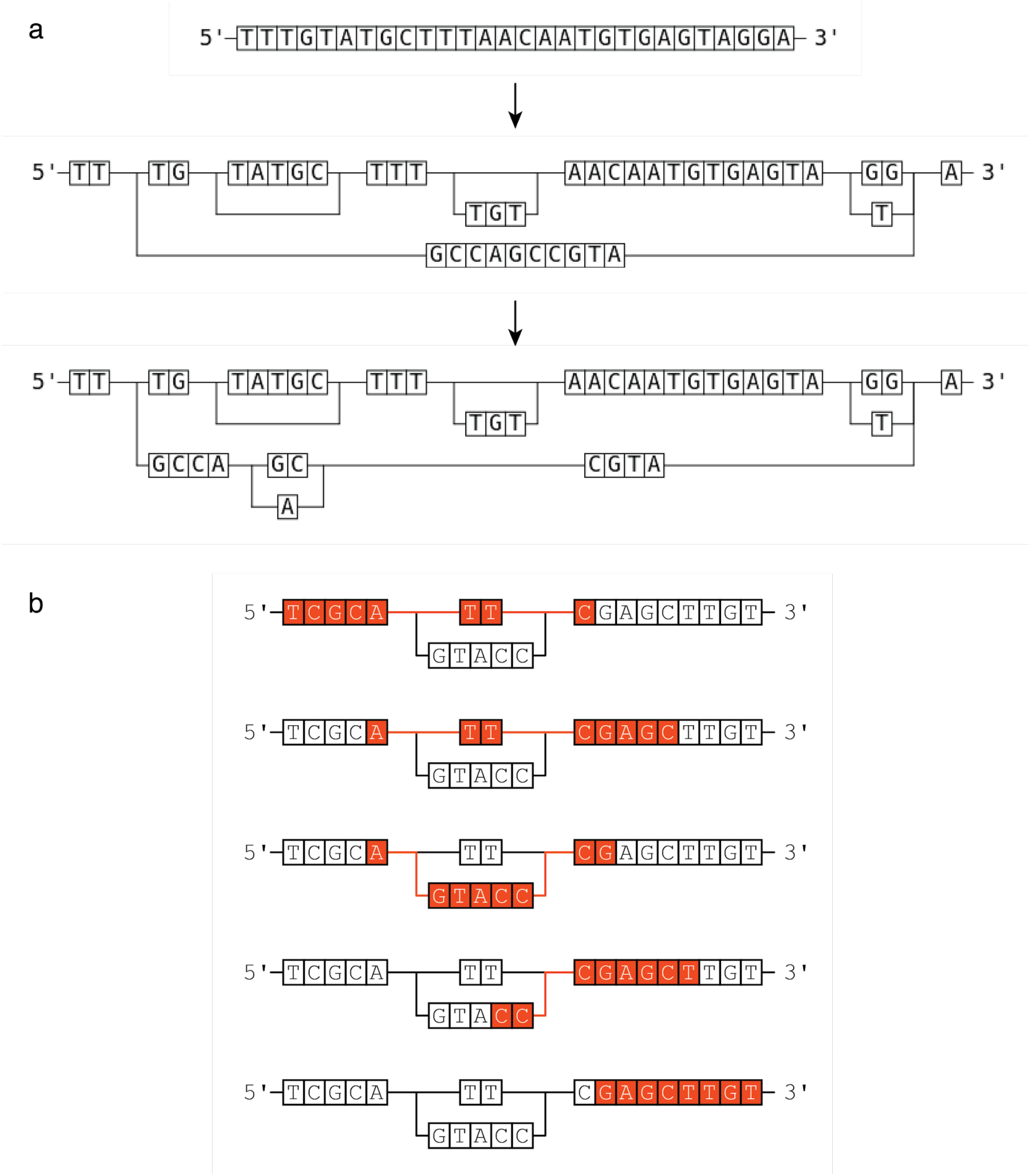
(a) Local graphs illustrated in Figure 1a were created and visualised using Graph Genome Pipeline from actual graphs. (b) k-mers (red bases) are indexed from every s-th position. If multiple k-mers can be generated from a given position due to an edge, all such k-mers are indexed. k- mers can also start on variant edges until they return to the parent edge. In this example, k=8 and s=4.

The search procedure is implemented as follows. For each *k* consecutive symbols in a read, a hash index is calculated by applying the hash function used during the search index construction; a list of *k*-mer loci corresponding to the calculated hash index is determined. We use sliding search window approach to locate substantial spatial clusters of loci belonging to an aggregate of *k*-mer lists determined for the read. These clusters represent candidate match regions. We allow for gaps between loci. A search window size is selected based on the read length and the upper limit for novel INDELs. Each located cluster is assigned a score which is calculated by analyzing matching *k*-mers, their positions in the graph and corresponding positions in the read. The higher the score, the larger the probability of match with the read. Clusters with scores exceeding a threshold are treated as seeds for local alignment. Each seed hence represents a region in the graph against which the read might align. Paired-end reads are treated as single search patterns with gaps. These are processed in the same way as single-end reads which significantly reduces the computational complexity of pair-read search.

#### 3.1.3. Local alignment

The majority of reads do not contain novel insertions or deletions. We exploit this by taking a hierarchical approach to local alignment: for each candidate seed we first attempt a fast gapless alignment algorithm and then, if required, a slower alignment algorithm which permits novel insertions and deletions. In both cases, the graph region is extended either side of the seed to accommodate cases where k-mers at each end of the read are not contained in the graph index or the read contains novel variations or comes from a repetitive region.

We use a graph-aware version of the bit-parallel approximate (BPA) string matching algorithm (also known as the bitap algorithm) for the first stage. Given a pattern P of size m and a text of size n, the score matrix, *R*, is defined as

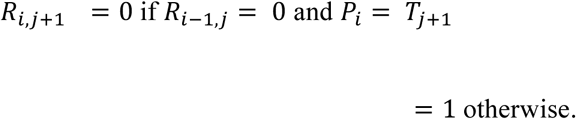

Therefore *R_i,i_* = 0represents the case where the first i characters in P match the last i characters in T that end at j. This can be efficiently implemented by creating a mask for each character in the alphabet spanned by P; the transition from j to j+1 can then be performed by applying a right shift to the j-th column (which is aided by using 0 to denote a match) and a bitwise AND with the character mask corresponding to the (j+1)-th character. The algorithm lends itself naturally to a highly optimised SIMD implementation. The extension to handling mismatches and gaps is relatively straightforward and has been previously described48–50.

Our extension of BPA to the graph exploits the bitwise nature of the algorithm: as the score at a query symbol depends upon the match from any incoming edge, incoming edges can be combined by a single bitwise OR operation.

The alignment can then be extracted by backtracking through the score matrices for each edge. Crucially the graph-aware BPA algorithm scales linearly with the number of edges in the region of interest and not the number of paths, allowing us to handle variation-dense regions of the genome.

Dynamic programming is motivated by Bellman’s Optimality Principle^51^ and remains applicable for finding the lowest cost traversal in a DAG52. The commonly-used Smith-Waterman algorithm therefore generalises to finding the best matching path in a directed acyclic graph where the number of possible incoming states for the alignment of a query symbol is a multiple of the number of edges merging at the locus being considered. We exploit this to correctly map reads in presence of novel indels and structural variations not contained in the graph, using a custom SIMD-optimized implementation of the Smith-Waterman algorithm against the graph reference53 in cases where BPA fails to find an alignment with fewer than a given number of (default: 4) mismatches for a seed.

The alignment with the highest mapping quality is selected and the alignment is returned as a path in the graph and a CIGAR string relative to that path (i.e. relative to the concatenated genomic sequence from the edges visited in the alignment).

#### 3.1.4. Projection to the linear reference

An alignment for a read against the Graph Genome Reference is converted to an alignment against the linear reference for output in the BAM. The alignment position is given as the first base in the graph backbone (i.e. linear reference) which the read aligns against. In the case of alignments entirely inside an edge, as can happen in long insertions, the nearest position on the graph backbone is reported. The CIGAR describing the alignment against the graph is converted to a CIGAR relative to the linear reference by projecting the alignment against each edge to the linear reference (e.g. an alignment match against an *n*-bp insertion edge is reported as *n*I and a novel 1-bp deletion against an *n*-bp. insertion edge is reported as (*n*-1)I). Consecutive insertions and deletions, whilst possible in graph-based alignments, are not accepted by conventional downstream tools based on a linear reference. Such paths are hence merged into a single insertion or deletion event followed by mismatches, i.e. a CIGAR of xIyD is converted to (x-y)IyM if the insertion is longer than the deletion and (y-x)DxM otherwise. The alignment is soft-clipped by applying a constant soft-clip penalty and Kadane’s algorithm for the maximum subarray problem^54^.

#### 3.1.5. Single-end mapping quality

Single-end mapping quality calculations closely follow the approach implemented in MAQ^55^ and further refined for the BWA-MEM aligner^27^. Mapping errors originating from the reads that do not come from the (known part of the) reference genome are not calculated (for a more elaborate discussion, readers are referred to the supplementary materials of Li and Durbin 2009^55^). This leaves two possible sources of mapping errors: 1) the true site is not considered (i.e. the correct seed is missed), and 2) the true site is considered, but not selected as the mapping location.

##### Estimating the probability of missing the correct seed

The primary reason a correct seed would be missed is that for performance reasons the graph aligner only considers a number of highest scoring seeds. The probability that the correct seed is not in the set of considered seeds (denoted as Ω) can be estimated by utilizing seed match scores (described above):

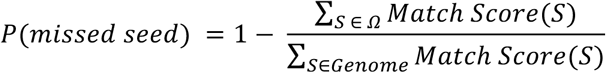

In practice, calculating the sum in the denominator over the entire reference structure would be prohibitive, so we approximate it with a number of top-scoring seeds, that is computationally feasible, but still significantly larger than the number of seeds processed by local alignment against the Graph Genome Reference.

##### Estimating the probability of selecting the wrong mapping location

This probability is the same as defined in the section 2.5.2 of the Supplementary Material of Li and Durbin 2009^55^:

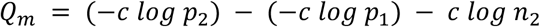

where *c* = 10/*log*10, *p*_1_ and *p*_2_ are the probabilities of the best and the second best hits, and *n* is the number of hits with the same score as the second best hit. The probability of a hit is estimated from the alignment scores, as done in Li 2013^27^. The major difference to linear aligner is that the alignment scores are calculated against the graph reference, i.e. reads are not penalized for passing through known variation.

#### 3.1.6. Paired-end mapping quality

In addition to the single end mapping quality described above, paired-end reads also receive a mapping quality by taking into the account both ends of a read pair. Calculation is similar to that for the single-end reads, but the alignment score for the pair is taken as the sum of the individual scores for the two mates. Before a read pair is considered “eligible” for the paired mapping quality value, two conditions must be met: (1) the read ends must be mapped in a proper pair, i.e. orientation must be correct and the estimated insert size must fall within the expected range for the library, and (2) at least one of the mates must be uniquely mapped (i.e. have a non-zero single end mapping quality). A read is assigned the greater of its single-end mapping quality and the quality calculated for the pair.

### 3.2. Reassembling Variant Caller

The Reassembling Variant Caller of the Graph Genome Pipeline takes elements from Samtools^55^, FreeBayes^56^ and HaplotypeCaller^57^. An important novel aspect of our approach is the use of the graph reference to help guide the assembly and genotyping phases (Supplementary Information 3.2.5), reducing the effects of reference bias.

We chose to use local reassembly over a simpler approach of direct genotyping of variants (i.e. attempting to maintain the representation of the called known variants as they are in the graph) for several reasons. First, it provides the same treatment for both known and novel events, while resolving the potential situation when reads originating from the same biological haplotype follow different paths on the graph depending on their different starting and ending positions and sequencing errors. Second, it resolves the situation where there is ambiguity over which branch a read should take, which is a common occurrence around the edges of repetitive sequences. Third, it provides a unified representation for the called variants even if graphs with different variant representations were used in the process. Finally, our approach allows the use of graphs with specialized structures optimized to improve the processing, such as hierarchical graphs to reduce sequence/k-mer repetition or short-range haplotype graphs that decrease the number of possible paths through the graph.

#### 3.2.1. Reassembly Window Formation

The reassembling variant caller (VC) starts on the reference at the beginning of the user specified interval. The VC makes one pass down the reference utilizing two pointers to call variants. The first pointer is the window formation pointer and the second pointer combines the variant windows into a reassembly window. The window formation pointer is nominally 300 bps ahead of the variant calling window. The first pointer is used to find parts of the genome that potentially contain variants. These possible variants are identified by the CIGAR string in each read. Each variant location is stored as a variant window. As the second pointer reaches each variant window it amalgamates all of the variant windows that are sufficiently close to each other into larger reassembly windows. Window amalgamation continues until all of the nearby variant windows are included or until the resulting reassembly window gets too large. In the latter case, window amalgamation ceases (even if there are other close variants) and the window is processed. The window is nominally 300 bps long, but can be extended further (we have tested up to 35 kb) to encompass larger events if they are present in the CIGAR. Once the reassembly window has been processed and any variants found, the second pointer continues one location past the last variant found.

#### 3.2.2. De Bruijn Graph Haplotype Generation

For each reassembly window we generate a set of putative haplotypes. We start by creating a De Bruijn-like graph (DBG)^58^, following the approach taken by GATK HaplotypeCaller^57^. The DBG is initially comprised of the k-mers extracted by running a sliding window over the reference sequence. In this process loops may arise in the DBG when the k-mers generated for graph creation are not unique. This leads to indeterminate sized haplotypes which should be avoided. Using a large k-mer size can eliminate the loops and solve this problem in many cases, but also exacerbate the effects of sequencing errors, low quality bases and limited read lengths. We therefore only attempt using moderate sized k-mers (by default we start with k-mers of size 15, and increase up to 85 in increments of 10). If there are still cycles at the maximum k-mer size, the reassembly window is skipped and no variants are called from it. Next, we also add the k-mers obtained by running a sliding window over all of the reads overlapping the reassembly window. While processing the reads, we look for non-unique k-mers, defined as k-mers that occur more than once within a single read. Such k-mers are added to the graph, but only connected to the previous and the next k-mer in the read. Non-unique k-mers will not show up in any subsequent k-mer searches and cannot be connected to, thus avoiding formation of loops.

For the k-mers extracted from reads, we estimate the probability that the k-mer is free of sequencing error by multiplying the per-base probabilities of correctness (derived from the base qualities produced by the sequencer) and discard the k-mers where this probability falls below a certain threshold (by default, the threshold is set to 0.991^k^, where k is the k-mer size). However, if a read’s k-mer aligns without mismatches against its corresponding sequence on the reference graph genome, this k-mer will not be discarded regardless of its error probability.

Once the DBG is created and pruned, haplotypes are derived via a depth-first search. By construction the DBG has one source node and one sink node. The source node corresponds to the first k-mer in the reassembly window and the sink node corresponds to the last *k*-mer in the window. In the absence of variants and errors, the DBG is just a single line of nodes from the beginning of the window to the end. Due to variations of the two haplotypes (in the diploid case) and read errors, there will be bubbles in the graph. Bubbles in the graph lead to multiple paths, each corresponding to a possible haplotype. Every path through the graph from source to sink node corresponds to a possible genome sequence that starts at the beginning of the reassembly window and stops at the end. This means all of the haplotypes start and stop at the same location on the reference which helps in identifying variant locations. These haplotypes represent estimates of sequences the original genome had in the reassembly window.

When there are many variants or errors in the data or the window is very long, many paths can result. This is greater than the number of paths that can be processed in a reasonable amount of time. We therefore process a subset (20 by default) of most likely haplotypes with the highest score computed as follows. For the first variant along the haplotype, we compute the ratio of the number of reads overlapping the variant and supporting the allele to the total number of reads overlapping the variant. Then, for each pair of alleles at consecutive variants along a haplotype, we compute the ratio of the number of reads that overlap with both variants and support both alleles, plus one, over the number of reads that overlap with both variants and support the first allele (without regard to whether they support the second allele), plus two. We multiply all such ratios along a haplotype to derive its score.

#### 3.2.3. Locating Potential Variants

The Needleman-Wunsch sequence alignment algorithm^59^ is used to map each haplotype in turn to the reference sequence within the reassembly window. The subset of loci in the reassembly window containing the locations of the potential variants is determined and processed further to find and genotype the variants.

#### 3.2.4 Likelihood Calculation, Variant Calling and Genotyping

The pair Hidden Markov Model (pHMM)^57,59^ calculates the likelihood of generating a pair of sequences according to a certain probabilistic model. For each haplotype, likelihoods are calculated for each read. If there are *m* haplotypes and *n* reads, a total of *m*×*n* likelihoods are calculated. These values are taken to be the conditional probabilities of the read given the haplotype, *p*(*r*|*h*). We use the base qualities to compute match and mismatch likelihoods, and use 10-4 and 10-1 for gap open and extension likelihoods, respectively.

To calculate the probability of a certain variant allele’s presence and its genotype, all reads that overlap a small window around the location of the variation are considered. For each read the likelihoods of the haplotypes are “marginalized” to derive likelihoods of the reference and each of the variant alleles present at the site on a per read basis. In this case, marginalization means picking the maximum of the likelihood values of all the haplotypes that contain an allele, *a*, so *P*(*r*|*a*) = *max_h containing a_p*(*r*|*h*).

The probability of the reads is considered conditionally independent given the allele, and so

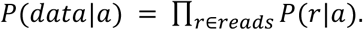

For the diploid case, the probability of the data for a given genotype <a,b> is shown by some reworking of the proof in supplementary material for Li et al.60 to be

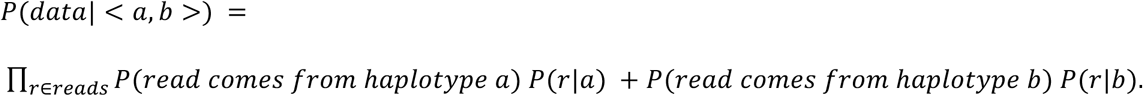

For variations where the length of variant a is va, the length of variant b is vb and the read length is r, let la = va + r - 1 and lb = vb + r - 1 the equation becomes

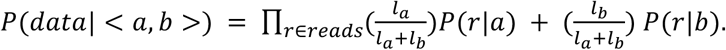

The probability of a variation in the case of one variant allele is

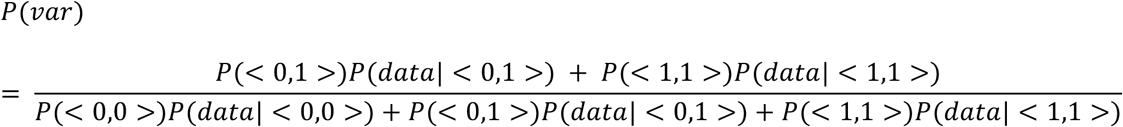

where the values *P*(<*a*,*b*>) are the assumed prior probabilities of the respective genotypes (see Supplementary Information 3.2.5).

If a variant is found, it is output in the VCF file.

#### 3.2.5. Utilizing Graph Information for Variant and Structural Variation Finding

The graph aligner outputs graph tags describing the alignment path through the graph for each read. This allows the VC to know which parts of each read represent the graph reference (either along the linear part of the reference or a branch) and which parts do not. This knowledge is used in by the VC in several ways.

First, when filtering the k-mers derived on reads based on base qualities, we avoid dropping k-mers which originate from portions of reads that are an exact match to the graph reference (whether to the backbone or some of the variant paths). The reasoning behind this choice is that a base call is unlikely to be erroneous in a way that would make it match to a known allele. Second, when calling the variant, we increase the prior probability of the site containing the alternative allele if that allele is present in the graph. By default, heterozygous alternate alleles are given a prior probability of 10^−2^ if in the graph or 10^−4^ if not in the graph. In both cases, the prior of a homozygous alternate is one-half that of a heterozygous alternate. In the future, we will adjust these priors based on population allele frequency.

Graph information is also used in finding structural variations (SVs) that are included in construction of the graph. When the VC comes across reads (by default, we require at least two reads mapping the same event) whose graph tags suggest it aligns across a long branch, the reassembly window is extended to encompass the entire branch. Such windows are then processed in the same manner as the other, shorter windows.

### 3.3. Standard Hard Variant Filters

We use a set of filtering criteria to filter out low quality (likely false positive) variant calls from the raw set of variants produced by the variant caller. The criteria are similar to the hard filtering best practices suggested by the Broad Institute for use with GATK tools^61^.

We filter out any SNP calls that fail one or more of the following tests:

- Alternate allele is supported by at least 20% of the reads.
- Mean Base Quality^61^ of the reads supporting the alternate allele is greater than or equal to 15.
- Variant quality divided by Depth^61^ is greater than or equal to 2.
- Rank sum test for mapping quality of reference vs. alternate allele supporting reads^61^ being different is greater than or equal to −12.5.
- Rank sum test for positions of reference vs. alternate allele supporting bases^61^ within reads being different is greater than or equal to −8.
- Fisher strand bias test^61^ is less than 60.

For INDEL variants we filter out any calls that where the variant quality divided by depth is less than 2.

### 3.4. Adaptive Autonomously Trained Variant Filter

We sought to compare Graph Genome Pipeline against a wide range of pipelines benchmarked in a systematic fashion and controlled experiment using the same input data and against the same truth set. The richest source of such benchmarks is Precision FDA Truth Challenge (https://precision.fda.gov/challenges/truth/), which includes over 30 benchmarked pipelines, many of which were employ supervised machine learning-based variant calling and filtration approaches.

As an alternative to our hard filters, we have developed an adaptive model trained on a validated set of variant calls supplied by GiaB. This filter is able to make decisions about the eligibility of genotypes based on joint consideration of various attributes calculated by the variant caller for each variant. It uses a simple logistic regression model trained on a subset of attributes from the VCF INFO field. We trained separate models for candidate SNP and INDEL variants. The features used for the adaptive filter are the following.

- Rank sum test for the mapping qualities of the reads supporting the reference vs. the alternate allele, as calculated by Van der Auwera et al.^61^
- Rank sum test for the base positions of the reads supporting the reference vs. the alternate allele, as calculated by Van der Auwera et al.^61^
- Rank sum test for the base qualities of the bases supporting the reference vs. the alternate allele. This metric is calculated in the same manner as the two above^61^, but using supporting read base qualities as input instead of positions and mapping qualities.
- Fisher’s exact test for strand bias of the reads at the variant site, as calculated by Van der Auwera et al.^61^
- Mean mapping quality of reads overlapping the variant site
- Variant quality divided by depth
- Symmetric odds ratio (SOR) test for the strands of the reads supporting the reference vs. the alternate allele, as calculated by the Broad Institute’s GATK StrandOddsRatio annotation, described at https://software.broadinstitute.org/gatk/documentation/tooldocs/current/org_broadinstitute_hellbender_tools_walkers_annotator_StrandOddsRatio.php
- Ratio of the number of reads supporting the reference allele and the total depth
- Ratio of the number of reads supporting the alternate allele and the total depth
- Variant quality
- Genotype quality
- Length of the homopolymer preceding the variant site, counted as the number of identical consecutive bases starting from the base preceding the variant position
- Mean base quality of bases supporting the reference and the alternate alleles separately
- Mean mapping quality of reads supporting the reference and the alternate alleles separately
- Number of soft-clipped reads supporting the reference and the alternate alleles, computed separately for each allele and each read orientation (forward vs. reverse)
- Mean number of alignment mismatches in reads supporting the reference allele and the alternate allele separately. Each deleted or inserted base is counted as a single mismatch. E.g. a read with the CIGAR “50=1×10=4I35=” would be considered to have 5 alignment mismatches.
- Mean number of mutation events in reads supporting the reference allele and the alternate allele, where a string of consecutive alignment differences is considered a single mutation event, thus a read with the CIGAR “50=1×10=4I35=” would have only 2 mutation events.
- The number of tandem repeat units in the reference allele and the alternate allele separately, if the variant in question is a tandem repeat expansion or contraction. Otherwise, this feature is set to 0. Repeat units are counted up to 50bp away from the variant reference coordinates, hence the maximum value this feature can get is (50 + max(0, <TANDEM repeat insertion length>))/<REPEAT unit length in bp>.

We replicated the experimental setup of the challenge, including the benchmarking software used, and used the version 3.2.2 of the GiaB truth sets^62^ for the GRCh37 human genome reference release. PrecisionFDA had provided two separate sequencing datasets for the Truth Challenge. The first was provided for the purpose of training variant calling pipelines and derives from sample HG001/NA12878 of Utah/European ancestry that NIST prepared using a PCR-free protocol and sequenced to approximately 50x coverage. For the actual evaluation, PrecisionFDA provided a dataset from HG002, the child of the Ashkenazim trio, generated using the same protocol and sequenced to approximately the same coverage. Performance is calculated against the aforementioned GiaB truth sets taking into the account only the sites specified as high confidence regions by GiaB. The adaptive filtration model was trained on the HG001 data and the trained model applied to the HG002 dataset.

After adaptive filtration the Graph Genome Pipeline achieves the best F1 score for both SNP and INDEL variants (Supplementary Fig. 6, Supplementary Table 7), and is the only pipeline to achieve high scores in both categories.

**Supplementary Figure 6.**
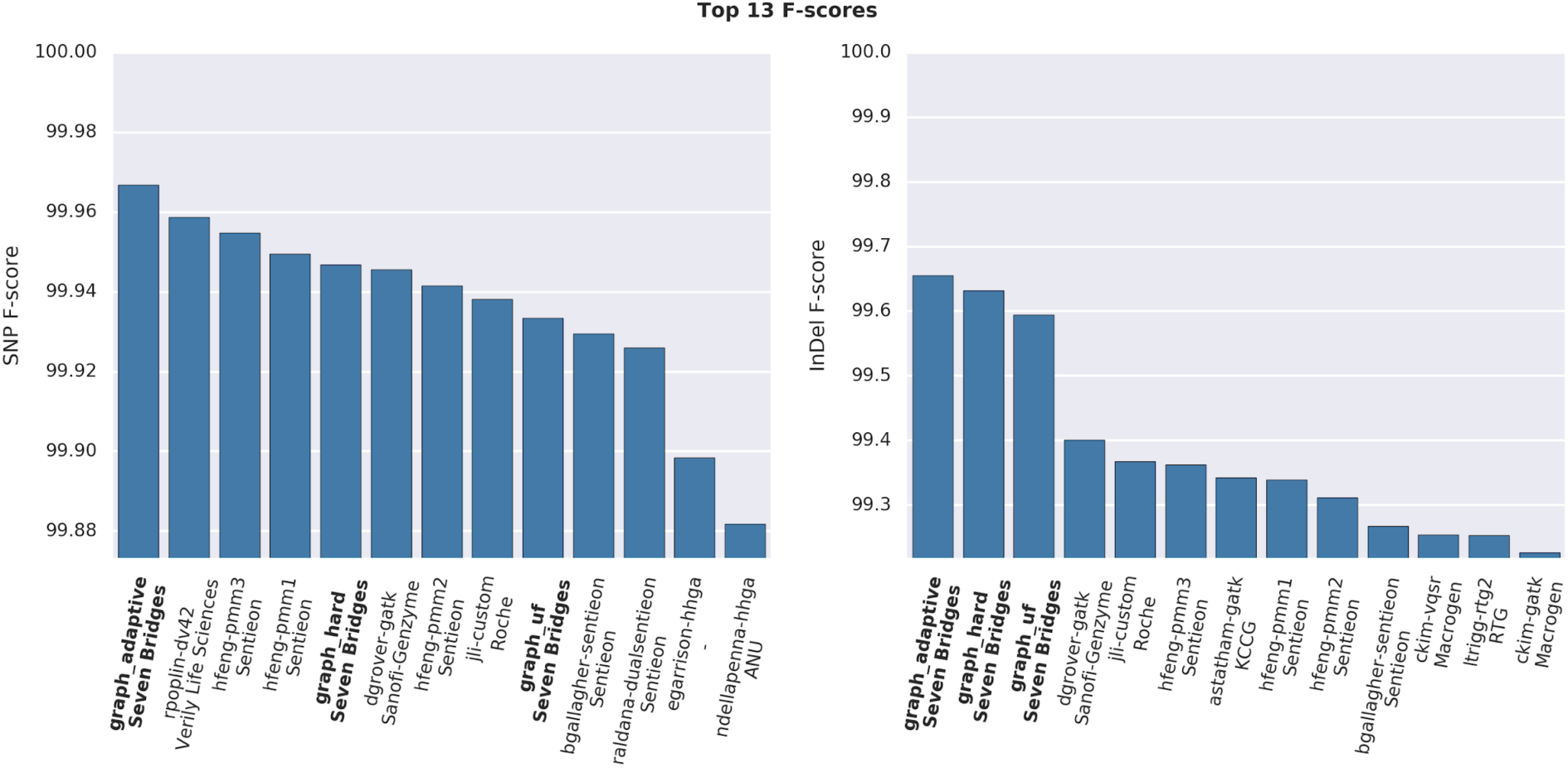
SNP/INDEL F1 score of the top ten scoring pipelines of Precision FDA Truth Challenge for both SNPs and INDELs along with three F-scores from the Graph Genome Pipeline. Graph Genome Pipeline Hard Filters were applied as outlined in Supplementary Information 3.3. The Graph Genome Pipeline using adaptive variant filtration achieves the best F-score in both categories.

A closer look at the both precision and recall (Supplementary Fig. 7) shows that the adaptive filtration method achieves a large increase in precision of SNP calls at a very low cost to recall. While the precision gain for INDELs is only moderate it still results in net positive increase in F1 score (Supplementary Figure 6).

**Supplementary Figure 7.**
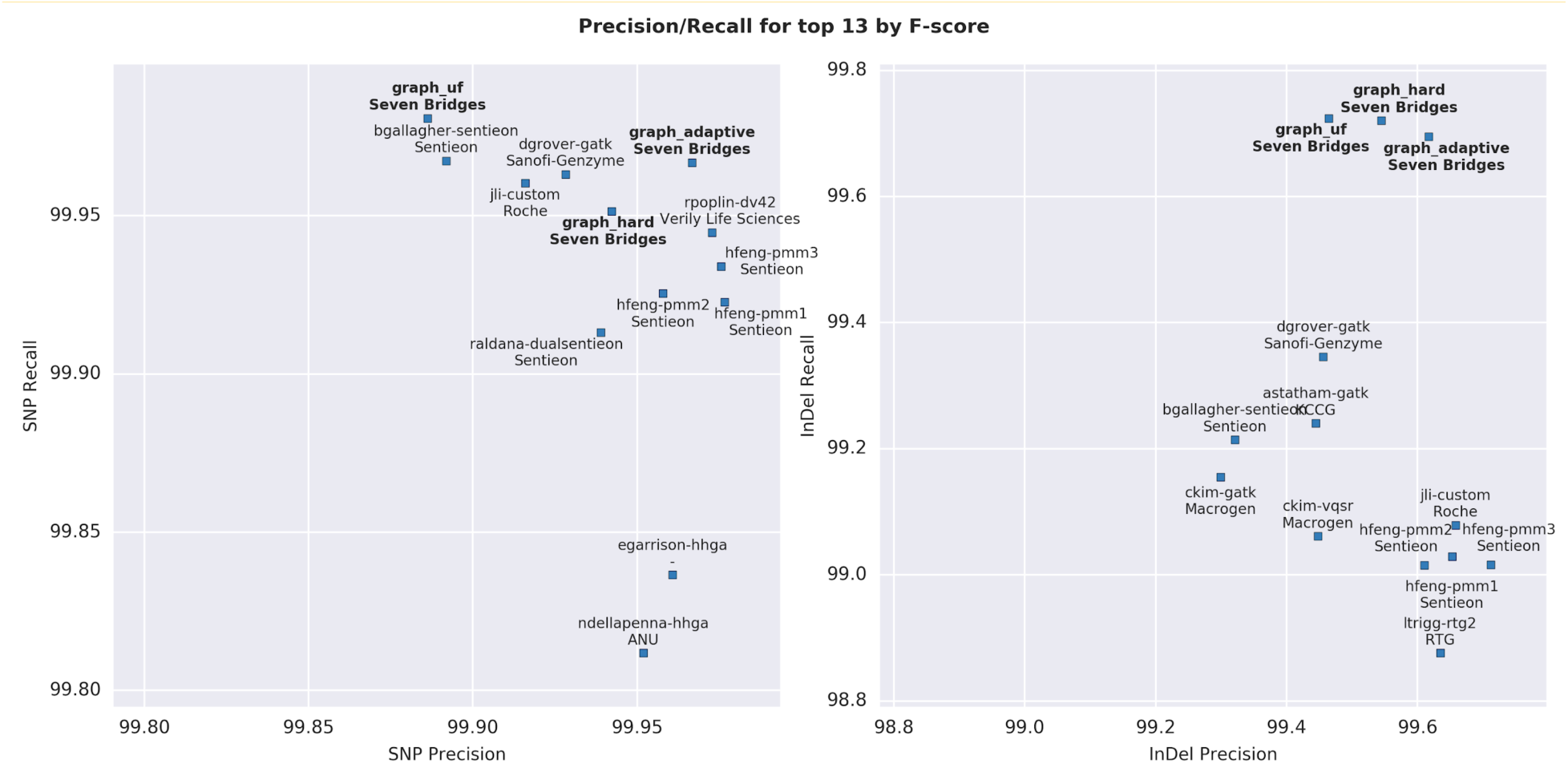
SNP/INDEL precision/recall score of the top ten scoring pipelines of Precision FDA Truth Challenge for both SNPs and INDELs with three Graph Genome Pipeline scores included as well. Graph Genome Pipeline Hard Filters were applied as outlined in Supplementary Information 3.3.

## 4. Testing and Benchmarking Methodology

### 4.1. Software, versions, and parameters for the aligner tools

#### 4.1.1. Graph Genome Aligner

Graph Genome Aligner version 0.9.1.1 was used in the experiments, by running the following command line:

*|| aligner --reference $FASTA --vcf $GRAPH -o $OUTPUT --threads $THREADS -q FQ1 -Q FQ2-- read_group_platform $PLT --read_group_sample $SAMPLE --read_group_library $LIBRARY*

#### 4.1.2. BWA-MEM

BWA-MEM27 version 0.7.13 was used in the experiments, by running the following command line:

*|| bwa mem -R $RG -t $THREADS $FASTA $FQ1 $FQ2 > $OUTPUT*

### 4.2. Software, versions, and parameters for the variant calling tools

Files produced by the two aligners were sorted using Sambamba^63^ version 0.5.9:

*|| sambamba sort --nthreads=$THREADS --memory-limit=2GiB --out=$OUTPUT $BAM*

#### 4.2.1. Graph Genome Pipeline

Graph Genome Pipeline consisted of Graph Genome Aligner version 0.9.1.1 and Graph Reassembling Variant Caller version 0.5.20. Variant calling was process-parallelized by running different instance of the caller to process different regions of the genome. Bcftools version 1.4 (http://www.htslib.org/) was used for merging the per-region VCF files, and applying the variant filters.

*|| caller -v $OUTPUT -i $REGION -b $BAM -f $FASTA -g $GRAPH -x all -s 10*

*|| bcftools concat -o $OUTPUT -Ov $VCF1 $VCF2…*

*|| bcftools filter -o $OUTPUT -e “(TYPE=’snp’ && (INFO/ADR[1] < 0.20 || INFO/MBQ[1] < 15 || INFO/QD < 2 || INFO/MQRankSum < −12.5 || ReadPosRankSum < −8 || INFO/FS > 60)) || (TYPE=’indel’ && INFO/QD < 2)” -Ov -s FP $INPUT*

#### 4.2.2. BWA-GATK

We used GATK tools (https://github.com/broadgsa/gatk) from version 3.7. Base quality score recalibration was set to calculate the calibration tables on chromosome 20. Base recalibration and variant calling was process-parallelized by running different instance of the caller to process different regions of the genome. HaplotypeCaller was set to output a gvcf file, which was followed but GenotypeGVCFs tools to produce the VCF.

*|| java -Xmx$XMX -jar $GATK --T BaseRecalibrator --out $OUTPUT -nct $THREADS -R $FASTA -- input_file $BAM --knownSites $DBSNP --intervals 20*

*|| java -Xmx$XMX -jar $GATK -T PrintReads -nct 4 --out $OUTPUT -R $FASTA --input_file $BAM -- intervals $REGION --BQSR $BQSR*

*|| java -Xmx$XMX -jar $GATK -T HaplotypeCaller --out $OUTPUT -R$FASTA --input_file $BAM -- pcr_indel_model NONE --intervals $REGION--emitRefConfidence $GVCF --dbsnp $DBSNP -- allowNonUniqueKmersInRef*

*|| java -Xmx$XMX -jar $GATK -T GenotypeGVCFs --out $VCF --variant $GVCF -R $FASTA*

*|| java -Xmx$XMX -jar $GATK -T CombineVariants --out $OUTPUT --variant $VCF1 --variant $VCF2…*

For analyses using hard filters we used the following command lines:

*|| java -Xmx$XMX -jar $GATK -T VariantFiltration --filterName “QD” --filterExpression “QD < 2.0” -- filterName “MQ” --filterExpression “MQ < 40.0” --filterName “FS” --filterExpression “FS > 60.0” -- filterName “MQRankSum” --filterExpression “MQRankSum < −12.5” --filterName “ReadPosRankSum” -- filterExpression “ReadPosRankSum < −8.0” --variant $VCF --validation_strictness SILENT --unsafe ALLOW_UNINDEXED_BAM -R $FASTA --out $OUTPUT*

*|| java -Xmx$XMX -jar $GATK -T VariantFiltration --filterName “QD” --filterExpression “QD < 2.0” -- filterName “FS” --filterExpression “FS > 200.0” --filterName “ReadPosRankSum” --filterExpression “ReadPosRankSum < −20.0” --variant --variant $VCF --validation_strictness SILENT --unsafe ALLOW_UNINDEXED_BAM -R $FASTA --out $OUTPUT*

For analyses using VQSR we used the following command lines:

*|| java -Xmx$XMX -jar $GATK -T VariantRecalibrator -nt $THREADS --recal_file $RECAL --rscript_file $RSCRIPT --tranches_file $TRANCHES --input $VCF --use_annotationQD --use_annotation MQRankSum --use_annotation FS --use_annotation DP --use_annotation ReadPosRankSum -- use_annotation MQ --use_annotation SOR -R $FASTA --mode SNP - resource:dbsnp,known=true,training=false,truth=false,prior=2 $DBSNP - resource:hapmap,known=false,training=true,truth=true,prior=15 $HAPMAP - resource:omni,known=false,training=true,truth=true,prior=12 $OMNI - resource:1000G,known=false,training=true,truth=true,prior=10 $KG*

*|| java -Xmx$XMX -jar $GATK -T ApplyRecalibration -nt $THREADS --out $OUTPUT --input VCF -- ts_filter_level 99.5 --tranches_file $TRANCH -R $FASTA --recal_file $RECAL --mode SNP*

*|| java -Xmx$XMX -jar $GATK -T VariantRecalibrator -nt $THREADS --recal_file $RECAL --rscript_file $RSCRIPT --tranches_file $TRANCHES --input --input $VCF --use_annotation DP --use_annotation FS --use_annotation ReadPosRankSum --use_annotation MQRankSum --use_annotation QD -- use_annotation SOR -R $FASTA --mode INDEL --maxGaussians 4 - resource:dbsnp,known=true,training=false,truth=false,prior=2 $DBSNP - resource:mills,known=false,training=true,truth=true,prior=12 $MILLS*

#### 4.2.3. Graph Aligner - HaplotypeCaller

We used the same version and the command lines HaplotypeCaller and the hard filter as described under BWA-GATK.

#### 4.2.4. BWA-Freebayes

Freebayes^56^ version 1.10 was process-parallelized by running different instance of the caller to process different regions of the genome. bcftools version 1.4 was used for merging the per-region VCF files, and applying the variant filters.

*|| freebayes -v $VCF -r $REGION -f $FASTA -= $BAM*

*|| bcftools concat -o $OUTPUT -Ov $VCF1 $VCF2…*

*|| bcftools view -o $VCF -Ov -e “QUAL < 30” $VCF*

#### 4.2.5. BWA-Graphtyper

We used Graphtyper^12^ version 1.3. Variant calling was process-parallelized by running different instance of the caller to process different regions of the genome. We used bcftools to merge the per-region VCF files and apply the suggested filters:

*|| bash make_graphtyper_pipeline.sh $BAM $CONFIG*

*|| bcftools concat -o $OUTPUT -Ov $VCF1 $VCF2…*

*|| bcftools view -o $OUTPUT -i “(ABHet < 0.0 | ABHet > 0.30) & MQ > 30 & QD > 6.0” $INPUT*

#### 4.2.6. BayesTyper

We used Bayestyper version 1.1 (https://github.com/bioinformatics-centre/BayesTyper) with 36 threads, followed by bcftools to exclude the uncalled sites from the VCF file. We used the following command lines:

*|| /bin/bash run.sh 55 $FQ1 $FQ2 $FASTA $OUTPUT $THREADS > terminal-out.log*

*|| bcftools view -o $OUTPUT --exclude-uncalled -e ‘GT=“0/0”’ $VCF*

### 2.3 Software, versions, and parameters for the variant calling benchmarks

For evaluating the accuracy of the variant calls (against simulated data, GiaB truth-sets, and the SNP Array truth-sets) we used vcfeval version 3.6, followed by hap.py version (this is the same setup as used by the recent PrecisionFDA Truth Challenge: https://precision.fda.gov/challenges/truth/results).

*|| rtg vcfeval -t $SDF -o $OUTPUT -T $THREADS --ref-overlap -m ga4gh -f “QUAL” -c $VCF -b $TRUTH --all-records*

*|| python qfy.py -o $OUTPUT --write-vcf --write-counts --verbose -t ga4gh --stratification $STRATS -- roc-delta 20 --roc $QQ -r $FASTA $VCF*

### 4.3. Aligner performance benchmarking and resource requirements

We used 10 randomly selected samples from the EUR population from the Coriell repository (http://www.ebi.ac.uk/ena/data/view/PRJEB20654 (Supplementary Table 5). We aligned these samples using 36 threads on AWS c4.8xlarge instances that have 36 CPU cores and 60 GB of random access memory. Tests with varying number of threads was done on a randomly selected sample, ERR1955509, (Supplementary Table 6). Supplementary Figure 8. depicts the genome coverage produced by the two aligners for these ten samples.

It should be noted that in our tests BWA-MEM used significantly less memory (5.3 GB) when aligning a small dataset of 10 thousand reads, pointing to a possible implementation issue with the tool.

**Supplementary Figure 8.**
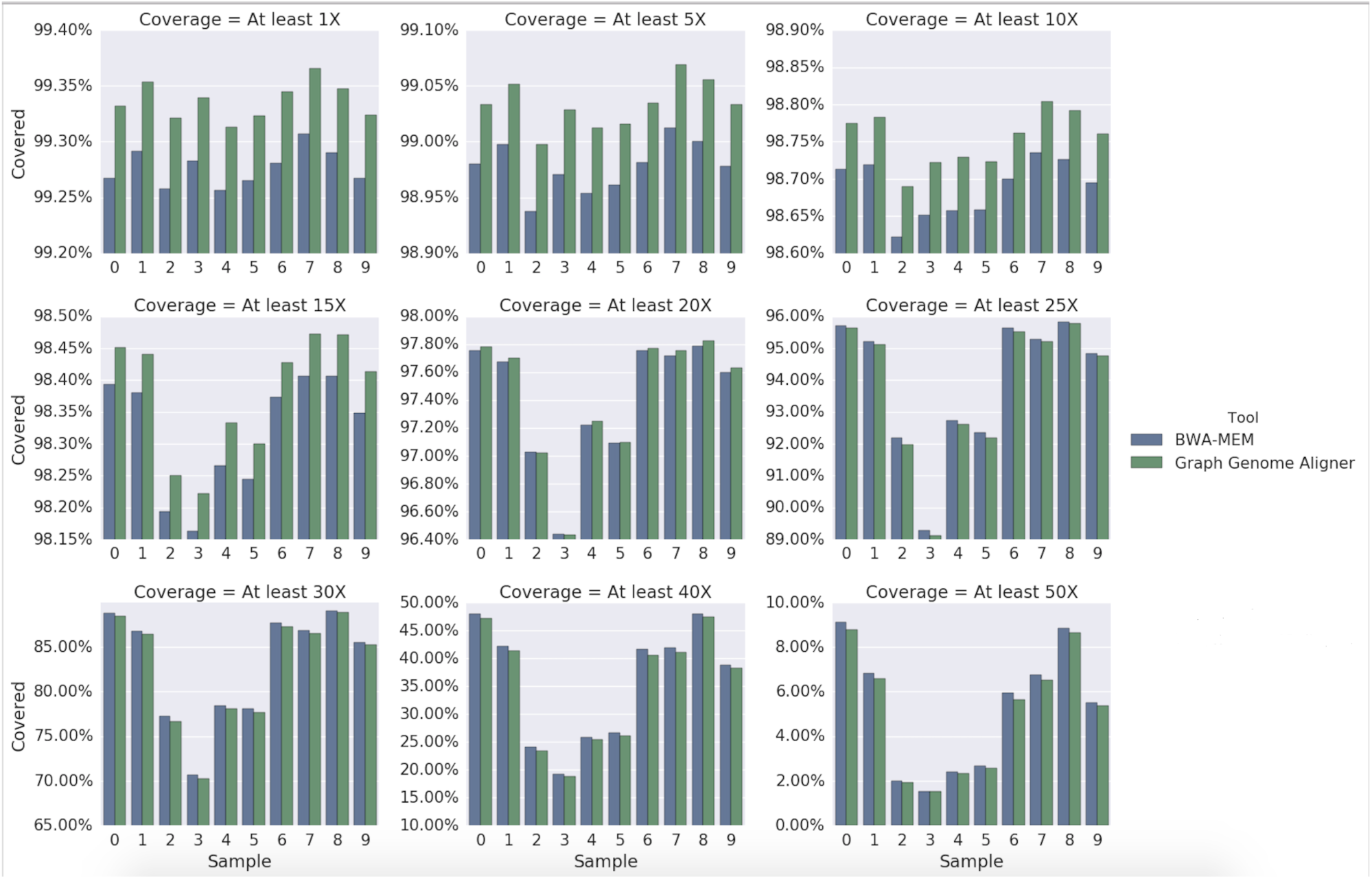
Coverage produced by the Graph Genome Aligner and BWA-MEM for ten 40X WGS samples from the Coriell dataset.

### 4.4. Benchmarking Experiments Using Simulated Datasets

We simulated reads for 10 genomes using the open source Mitty program (https://github.com/sbg/Mitty). As truth VCFs we used five samples from the GiAB project (HG001, HG002, HG003, HG004, HG005) and five samples from the 1000G Phase 3 release (HG00096, HG00551, HG03585, NA12878 and NA18488). The VCFs were filtered to remove complex variants, overlapping variant calls on the same chromosomal copy (e.g. a SNP placed in the middle of a deletion) and duplicate variants. The alleles specified by the genotype set for each genome were combined with the GRC37 reference FASTA to obtain sample-specific diploid DNA sequences. The read simulator is ploidy-aware, so where phasing information is available in the VCFs the variants are placed on corresponding chromosome copies. Where there is no phasing information, variants are placed on the first parental copy of each chromosome. Mitty generates Illumina-like reads in FASTQ format by stochastically sampling these DNA sequences, and inserts realistic base call errors (substitution only) generated from an error model derived by sampling base quality scores from a GiaB FASTQ file.

We generated 50x coverage reads from the whole genome excluding regions with masked reference sequence (i.e. those with ‘N’s). These FASTQ files were supplied to the pipelines under study and the alignments and variant calls were analyzed for correctness. To assert the alignment correctness of a read, we find its first base that was both derived from the linear reference (as opposed to an insertion branch) and aligned to the linear reference by the aligner. A read is considered correctly aligned if the alignment positions of these bases match (Supplementary Figure 9). Variant calls were assessed against the VCF used for simulation using vcfeval in the same way as the GiaB truth sets were assessed (Supplementary Information 4.5).

**Supplementary Figure 9.**
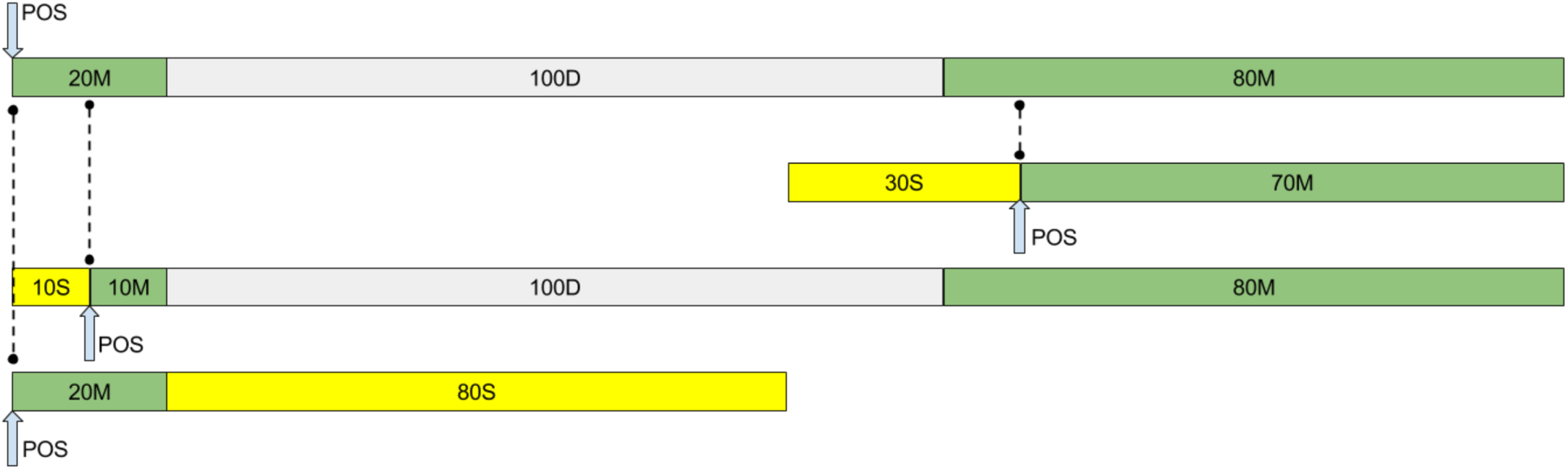
Simulated read alignment scoring. The top most diagram shows the alignment of the original, simulated read. Normal CIGAR nomenclature is shown. Our alignment scoring algorithm gives all the subsequent alignments a perfect score. In the first alignment (30S70M) the first reference matching base (indicated by the large arrow) is placed on the reference in the same position as that base in the simulated read (shown by the dotted line). This is also true for the second (10S10M100D80M) and third (20M80S) alignments.

#### 4.4.1. Simulated read alignment performance excluding microsatellite and segmental duplication regions

Here we show simulated read alignment performance stratified to exclude reads that were simulated from microsatellite and segmental duplication regions (Supplementary Figure 10). We can see that the difference between Ref and SNP reads is now much less pronounced. This is because the vast majority of variant bearing reads are from regions that exclude micro-satellites and segmental duplications while a certain fraction of Ref reads do also come from these hard-to-align-to regions. Excluding these difficult regions from the analysis changes the statistics for Ref read alignment, but not for variant bearing reads.

**Supplementary Figure 10.**
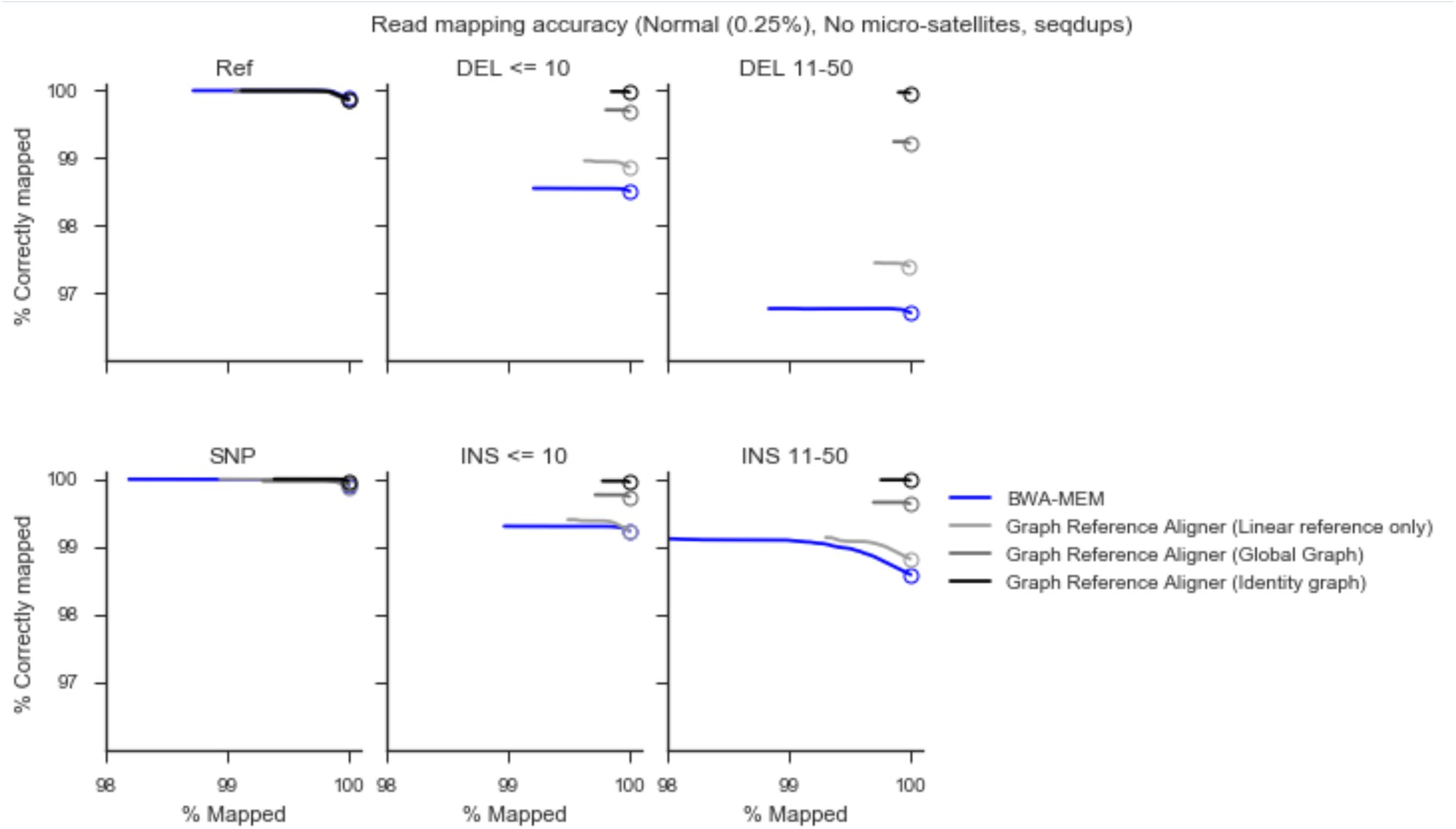
Alignment accuracy comparison stratified to exclude simulated reads whose true positions overlap with microsatellites or segmental duplications.

#### 4.4.2. Effect of sequencing error rates on simulation results

We performed the simulations presented in Fig 2. with three additional sequencing error rates while keeping all other parameters such as read length and template size distribution the same (Supplementary Figure 11). The impact of different sequencing error rates on read alignment results are shown in Supplementary Figure 12. In terms of alignment accuracy, the graph reference aligner performs better at all error levels. At the highest error level simulated, even though BWA-MEM maps a greater fraction of reads, the Graph reference aligner is more accurate. Simulated variant calling results at varying error rates are show in Supplementary Figure 13. The Graph Genome Pipeline and Graph Aligner + GATK-HC remain the best pipelines for INDELs in all error rates. In SNP calling, the Graph Genome Pipeline outperforms others when the error rate is less than 2%. At higher error rates the performance of BWA-GATK and Freebayes are the best.

**Supplementary Figure 11.**
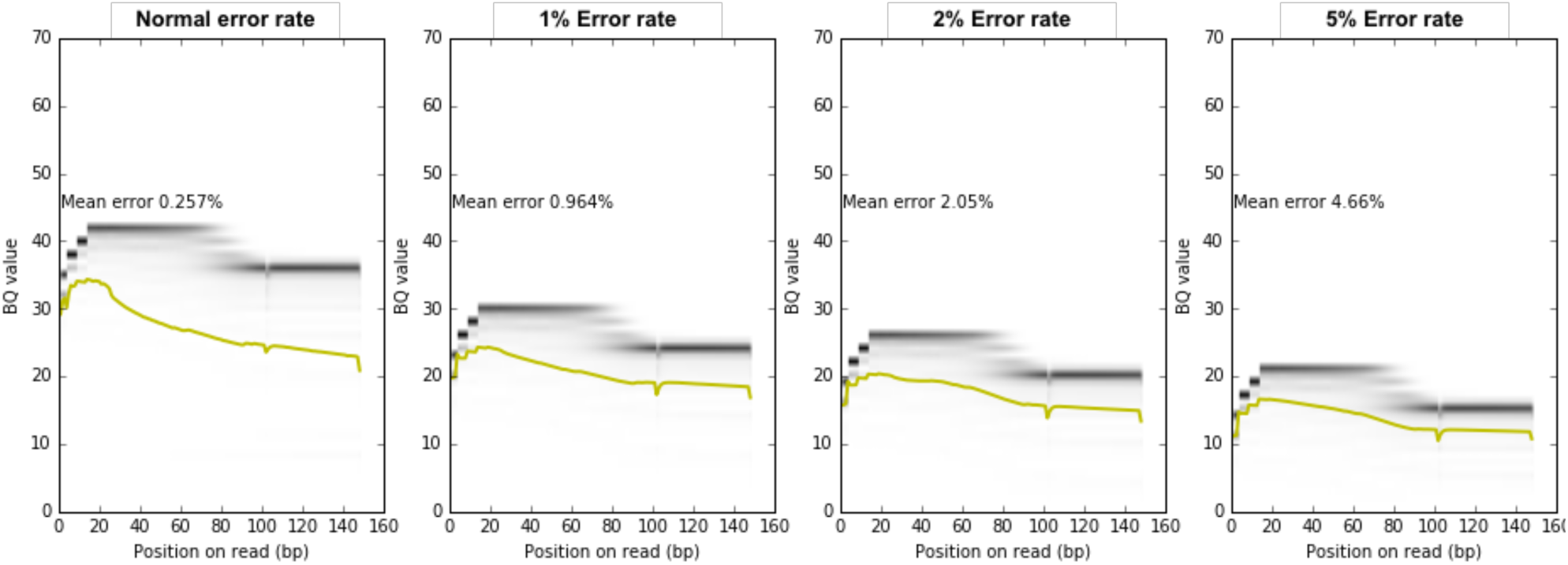
Base quality plots for the “normal” read model and the modified versions that gradually change sequencing error rate. The X-axis indicates the base of the read and the Y-axis represents the base quality value. The heat map shows the base quality distribution, while the yellow solid line shows the mean base quality values for each base. The “Normal” model is extracted from a GiaB FASTQ sample. The modified versions are generated by linearly shifting the base quality distributions down by one or more bins as required to get close to the target error rates.

**Supplementary Figure 12.**
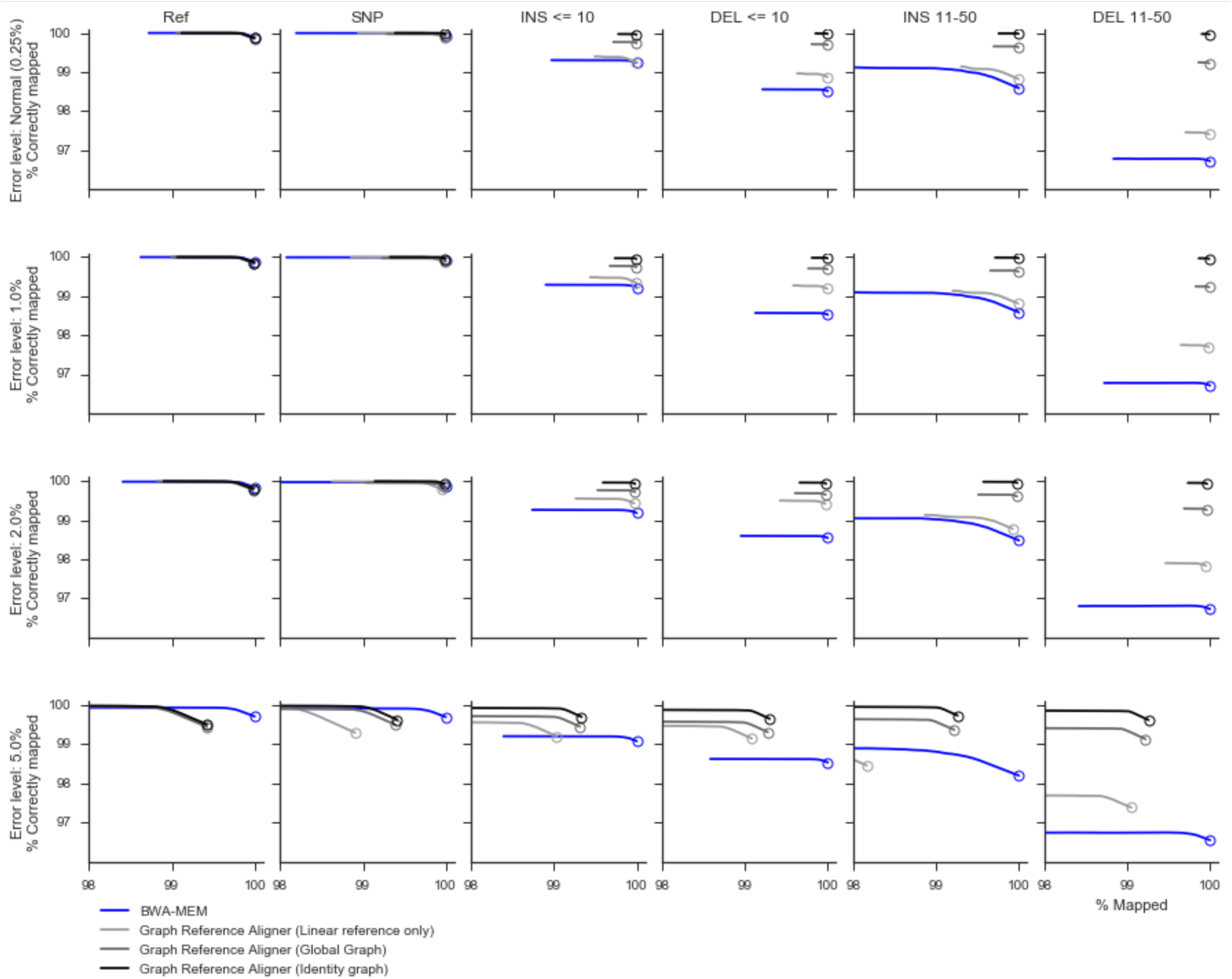
Alignment accuracy comparison at different simulated sequencing error rates. For each plot, the X-axis indicates the percentage of reads mapped, the Y-axis the corresponding percentage of reads correctly mapped. The curves are parametrized by MQ cutoff as detailed in the Figure 2 in the main text. The circle markers correspond to having no MQ cutoff. Each plot panel shows alignment accuracy for reads covering different classes of variants: Ref (no variant covered by the read), SNP (one SNP covered by the read), INS/DEL (one insertion/deletion of given size range covered by the read). The simulated sequencing error rate is indicated in the figure titles. The empirical sequencing error rate for this dataset is about 0.25%. At all sequencing error levels the graph aligner has a better accuracy, although BWA-MEM maps more reads at the highest error level tested.

**Supplementary Figure 13.**
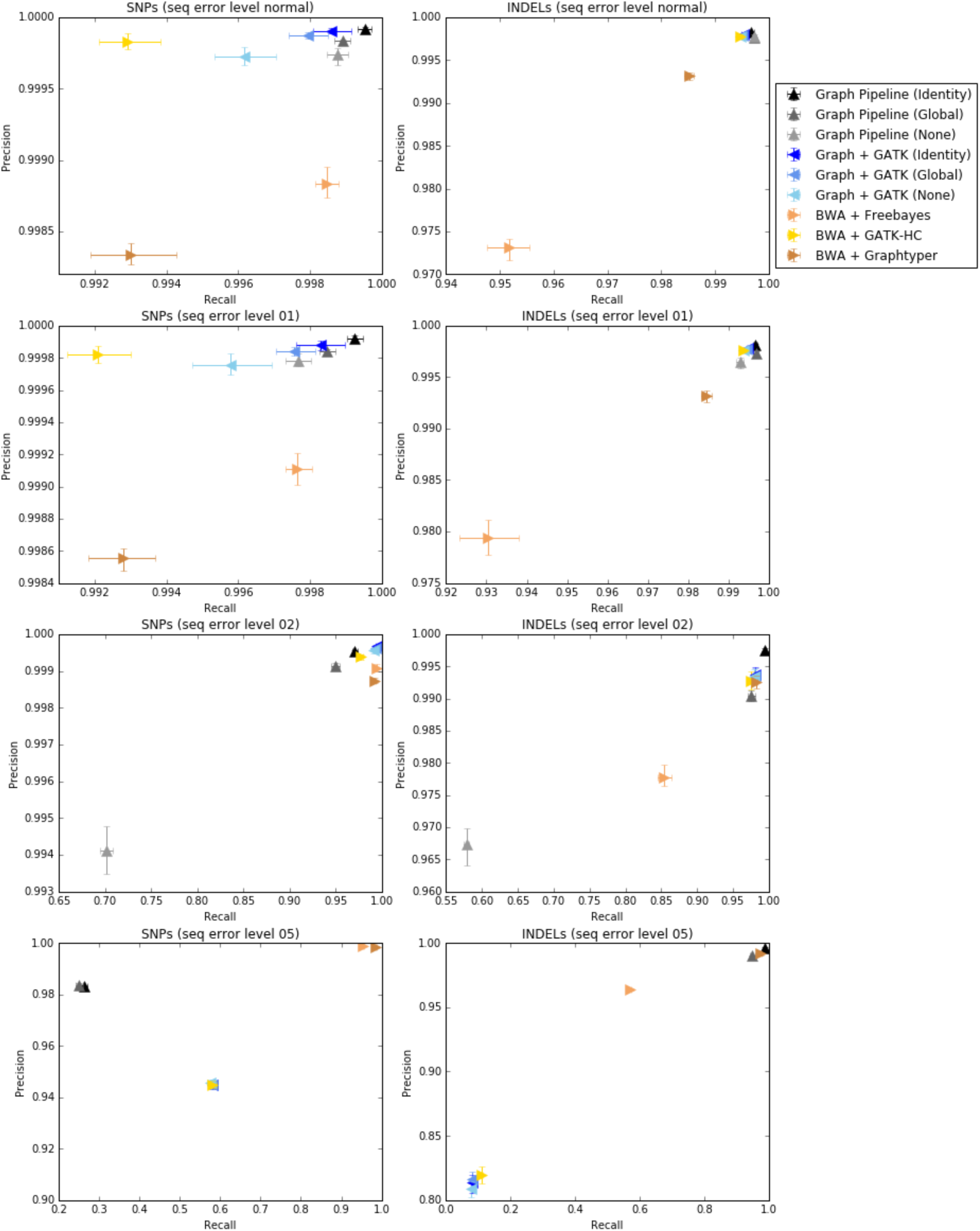
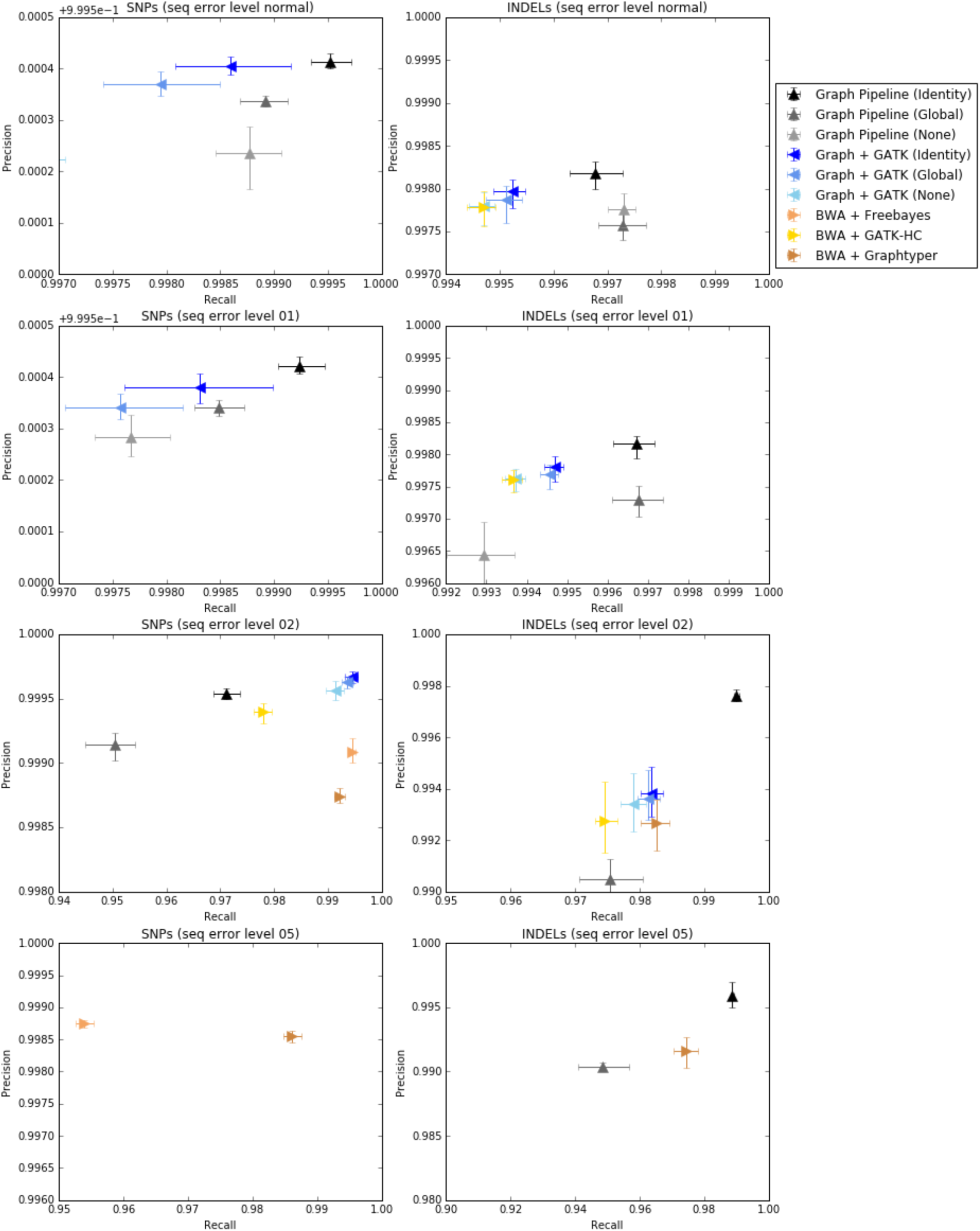
Variant calling accuracy comparison at different simulated sequencing error rates across different pipelines tested. For each plot, the X-axis indicates recall and Y-axis indicates precision. For each condition 10 simulated samples are run and the mean and bootstrapped 95% confidence intervals are shown. The top panel shows all the data while the bottom panel shows just the highest scoring pipelines for a better comparison

#### 4.4.1. Simulations demonstrate bias towards standard test samples

Simulation results with samples taken from 1000G VCFs and the five GIAB truth set VCFs suggest that pipelines have relatively high accuracy on the two standard samples (HG001, HG002), but not for other samples that are not commonly used to assess pipeline performance. It is apparent that the relative accuracy of the pipelines in the high-confidence regions is not a good predictor of the accuracy in more heterogeneous regions of the genome.

The results from the pipelines (Supplementary Figure 14) can be compared against the original genotypes, in specific regions, providing a reliable way to estimate sensitivity and specificity in any given region of the genome. The accuracy for HG001 and HG002 (Figure 3a in the main text) can be contrasted with the accuracy of the samples simulated from the published genotypes for these samples.

**Supplementary Figure 14.**
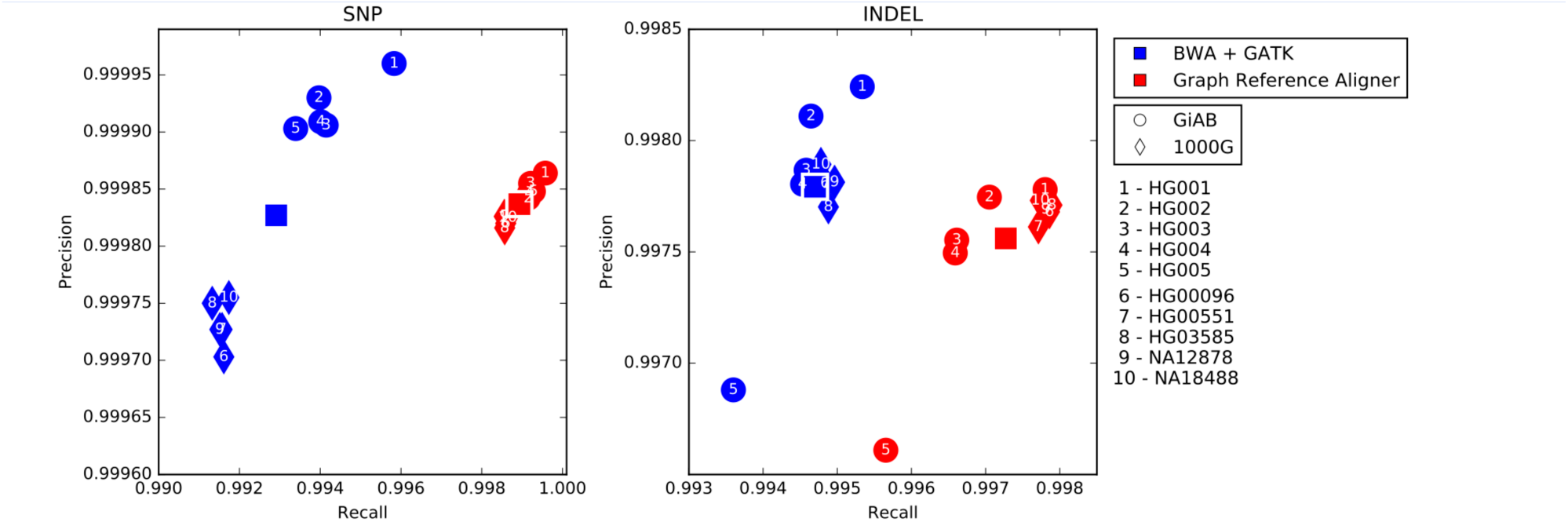
Precision and recall from 50x reads simulated using genomes from GiaB (points 1 - 5) and 1000G (points 6 - 10) samples for the BWA-GATK pipeline with hard filters and for the Graph Genome Pipeline. The squares indicate the average for all 10 samples for the respective pipelines. These are the same data points are those of BWA-GATK and Graph Genome Pipeline in Figure 3a.

### 4.5. Benchmarking against GiaB Truth-sets

Benchmarks (Figure 3b in the main text) were done on all 5 GiaB29,62 samples, HG001 (NA12878), HG002 (son of the Ashkenazi Jewish Trio), HG003 (father of the Ashkenazi Jewish Trio), HG004 (mother of the Ashkenazi Jewish Trio), and HG005 (son of the Chinese Trio). HG001-HG004 are sequenced to 50x coverage using PCR-free library preparation protocol and 2×150 sequencing reads, while HG005 is sequenced using 2×250 bp reads. All files were downloaded from the GiaB FTP site (ftp://ftp-trace.ncbi.nih.gov/giab/ftp/).

### 4.6. Benchmarking genome determination pipelines using family trios

Family trios have been used as truth-free benchmarks for variant callers by leveraging Mendelian inheritance rules and the low de novo germline mutation rate in humans^64–74^. We used three different methods for estimating the rate of Mendelian compliance rate in a family trio, where variant calling is performed independently for each member of a family, i.e. Mendelian inheritance information is not directly used in variant calling. The first method is the simplest and is based on detection Mendelian violations through line-by-line comparison over the VCF files of a family trio. Results from this method are shown in Supplementary Fig. 15.

**Supplementary Figure 15.**
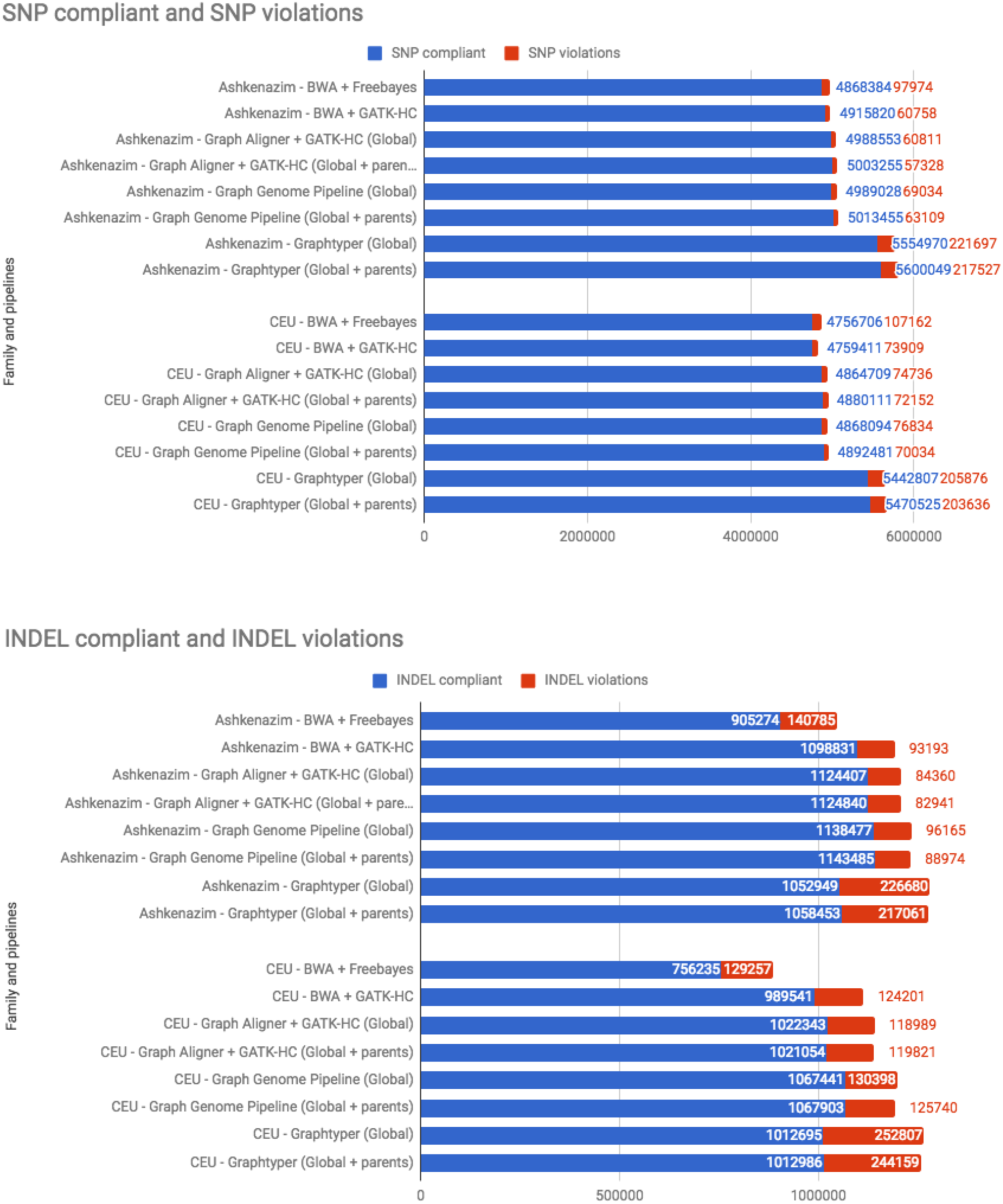
Simple Mendelian compliance and violation counts between pipelines.

The second trio comparison method is also based on counting Mendelian violations in a given family trio, but involves variant representation resolution. It has previously been reported that due to ambiguous representation of variants in the VCF format, Mendelian compliant variants in a family trio can be represented in an inconsistent manner, which leads to spurious Mendelian violation calls75. To address this issue, we extended the representation-agnostic variant comparison algorithm of vcfeval75 to family trio comparisons, with the goal of producing more accurate tallies of Mendelian violation rates. This method is detailed in section 4.6.1.

While straightforward tallies of Mendelian violation rates can be used to rank variant calling algorithms, they do not provide fine-grained information about the types of errors each algorithm makes (i.e. FP vs. FN rates). To this end, past studies have used the count distribution over the different types of Mendelian violations to statistically estimate false positive and false negative rates of a benchmarked algorithm^12,64–66,69^. We detail our methods for estimating these error rates using variant calls from family trios in section 4.6.2.

#### 4.6.1. Mendelian violation counting with variant representation resolution in family trios

The standard specifications of the VCF format is flexible and allows multiple different representations for one or multiple variants. It has been established that this flexibility causes the same variants to be matched between two VCFs have different representations for them^75^. Moreover, simple single-variant normalization approaches such as left-shifting are insufficient to making different representations fully consistent, but variant representations must be consolidated by considering adjacent variants in each VCF being compared. Indeed, a similar multi-variant representation consolidation scheme is the standard practice in the GA4GH guidelines for variant comparison^45^, and we have followed these guidelines in our single sample variant calling benchmarking experiments described in above sections 4.4-4.6.

Several tools, e.g. RTG Mendel^76^, GATK SelectVariants (https://software.broadinstitute.org/gatk/documentation/tooldocs/4.beta.6/org_broadinstitute_hellbender_tools_walkers_variantutils_SelectVariants.php) and Vcftools Mendel (https://vcftools.github.io/man_latest.html), exist for counting Mendelian violations in a given trio through naive variant comparison. In these approaches, each record in the merged trio VCF is processed independently and only variants with identical positions on the reference genome are analyzed together. This is analogous to the case when two VCFs are compared variant by variant without representation resolution and as a consequence, ambiguous variant representation also affects Mendelian violation analysis in family trios. Supplementary Figure 16 illustrates how inconsistent variant representation can cause a Mendelian compliant inheritance pattern to appear as multiple Mendelian violations.

**Supplementary Figure 16:**
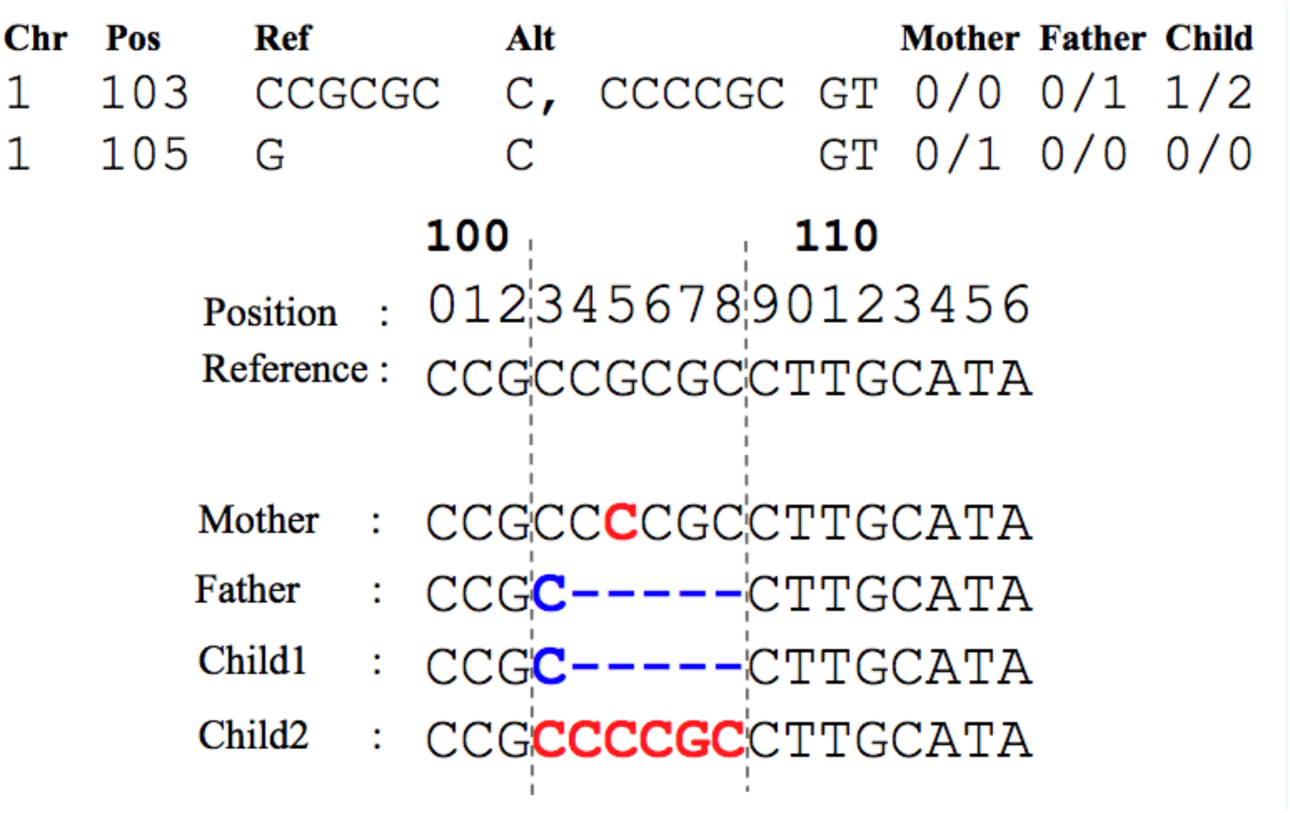
Example of variant representation difference. Naive trio comparison tools categorize the first row as a Mendelian violation since the second allele of the child does not exist in the mother. However both variants are actually Mendelian consistent when they are applied to the reference sequence simulteneously.

Vcfeval resolves the variant representations between two diploid VCFs, a baseline (gold standard) and a called (test) VCF, by assigning each allele in each variant into one of the two haplotypes such that the number of consistent variants and alleles between the haplotype sequences constructed from each VCF are maximized. To describe a haplotype of a sample formally, vcfeval uses a binary phasing vector PV = {p1, p2, … p|V|} ∈ {0,1}|V| to indicate the selection of the first, e.g. maternal, (if pi = 0) or the second, e.g. paternal (if pi = 1) sample allele of the i-th variant in a set of variants V. Note that the values of pi ∈ {0,1} refer to the two alleles in each diploid variant position of a sample, and therefore at a given genomic position the same phasing vector value may refer to different bases in different samples. In other words, given the two parental alleles that a sample has at each of its variant position Vi, PV chooses one allele of that sample at each variant and combines them into a haplotype vector. Since the human genome is diploid, PV has a complementary phasing vector P’V = {1-p1, 1-p2, …, 1-p|V|}, which selects each allele not selected by PV.

The objective function used by vcfeval is

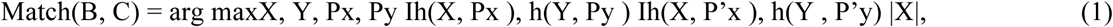

where B (C) denotes baseline (called) variant sets, X (Y) is the set of variants which maximizes the number of matching variants in the baseline (called) variant set and *X* ⊆ *B*, *Y* ⊆ *C*, PX∈ {0,1}|X|, PY∈{0,1}|Y|. Variant sets B and C are equivalent to the lines of VCF currently under comparison in the baseline and called samples, respectively. Similarly, variant sets X and Y are subsets of lines in B and C, respectively. Function h(X, PX) constructs a haplotype sequence by using the reference sequence and making the sequence substitutions dictated by phasing vector PX applied to variants X. Ih(X,Px),h(Y,Py) is the indicator function that is equal to 1 if h(X, Px) and h(Y, Py) are the same DNA sequence and 0 otherwise.

The indicator functions in Eq. 1 are multiplied by the number of variants in X, |X|, so Match(B, C) is maximised when the maximum number of variants in B are chosen such that a consistent pairing of haplotype sequences can still be found between the baseline and the called samples. The vcfeval manuscript details how Match(B, C) can be maximised with respect to variant subsets X and Y and their phasing vectors PX and PY.

We extend the vcfeval algorithm into a variant representation-robust Mendelian compliance comparison tool for family trios, consisting of variant set M from the mother, F from the father and C from the child. In this case, we seek to segregate the child alleles, including as many variant positions in the child as possible, into two haplotypes that can still be reconstructed from the alleles of the parents at each parent variant. Formally, our objective function following vcfeval’s notation is

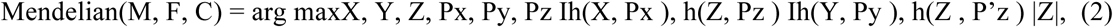

where *X* ⊆ *M*, *Y* ⊆ *F*, *Z* ⊆ *C*. The number of Mendelian compliant variants in the child can then be obtained by taking the number of variants in Z that maximizes the function Mendelian(M, F, C) (along with the other maximization variables X, Y, PX, PY and PZ). The remaining child variants, C\Z, are reported as Mendelian violations.

For each variant r ∈ M∪F∪C there are three possibilities: r is excluded from the maximized variant sets (i.e. r ∉X, Y, Z), r is included in the maximized variants sets (ie. r ∈X, Y or Z) with the first allele (pr = 0), or r is included with the second allele (pr = 1). Thus, given the trio variant sets M, F and C, there are 3|M∪F∪C| possible combinations of X, PX, Y, PY, Z and PZ. Exhaustive enumeration of all possible variants and phasing vectors would be computationally costly. Instead, we match the variants between the mother-child and father-child pairs independently and then combine the matching results. In our experiments, this heuristic approach arrives at the correct variant matching result 99.99% of the time compared with the exhaustive search. A manuscript for our method, VBT, detailing the algorithm and the related validation experiments is currently under preparation (preprint available at https://www.biorxiv.org/content/early/2018/01/24/253492). Source code for VBT is available at https://github.com/sbg/VBT-TrioAnalysis. Below we give an overview of the steps taken by VBT.

First, a full genotype matching step is performed between the child and the mother. This step is carried out using the standard vcfeval algorithm (reimplemented in VBT) and yields a set of child and mother variants Xboth and Zmboth that are consistent for both alleles.

The mother’s and child’s variants that are not matched for both alleles are sent for an allele matching step. In this step, the objective function to be maximized is defined as

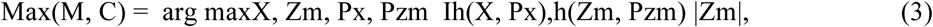

where Zm ⊆ C and PX∈ {0,1}|X|, PZm∈ {0,1}|Zm|. This equation has two differences to vcfeval’s objective function. First, there is only one indicator function, i.e. only one of the haplotypes is required to be matched between the child and the mother. Second, the phasing vectors Px and PZm here are only allowed to select non-reference alleles. Otherwise Eq. 3 could be maximised by including all the child’s heterozygous variants with a reference allele, and then having the phasing vector select the reference allele at each variant, effectively excluding each variant from haplotype comparison yet including it towards |Zm|. By only allowing the phasing vectors to include non-reference alleles ensures that all variants contributing to |Zm| actually contribute a non-reference allele in h(Zm, Pzm). Mendelian compliant heterozygous child variants with a reference allele are accounted for separately in step 6 below.

This step yields two different child variant sets: Zmnone, where none of the alleles can be matched with the mother’s alleles, and Zmone, where exactly one allele matches with the mother’s alleles. Note that mother and child variants sharing both alleles are already accounted for in step 1. Eq. 3 is maximized using vcfeval’s maximization algorithm repurposed for the changes in the objective function described above.

Steps 1 and 2 are repeated for the father-child pair, yielding Zfboth, Zfone and Zfnone.

Next, the variant sets Zmboth, Zmone, Zmnone, Zfboth, Zfone and Zfnone are checked for Mendelian violation. Child variants where both alleles can be matched with two alleles in both parents, i.e. Zmboth⋂Zfboth, are automatically considered Mendelian compliant. These are variants where the child has either a homozygous alternate genotype or heterozygous genotype with no reference alleles, and both parents have the same genotype as the child.

Of the remaining child variants, those that share both alleles with one parent and one allele with the other parents, i.e. (Zmboth⋂Zfone)∪(Zmone⋂Zfboth), are considered Mendelian compliant. These are the variants where the child’s genotype is heterozygous with no reference alleles, and one parent also has this same genotype while the other parent has one of the the child’s alleles.

A heterozygous child variant r ∈ ZHETREF where the child has a reference allele (i.e. excluding heterozygous variants composed of two non-reference alleles, or ZHETALT) must be treated separately, since the reference allele is excluded from the Max() function as described in step 2. For such variants to be rescued and considered Mendelian compliant, the following conditions must be satisfied. The variant must be part of the allele matching solution with one but not both of the parents, i.e. r must be in (Zmone⋂Zfnone)∪(Zmnone⋂Zfone). Now the non-reference allele is matched with one parent through Eq. 3, and the other parent’s variant set (i.e. VCF) is searched for a variant at the genomic position i of the variant r. If this parent has no variants at position i, or if the parent has a variant with a reference allele at position i, then and only then is the child variant r is considered Mendelian compliant and added to ZHETREF-CONSISTENT.

This reference allele matching step relies on naive position-based variant and allele matching. However, when combined with other steps, notably the full vcfeval-type sequence matches performed in steps 1 and 2, this step seems to cause relatively few Mendelian compliance checking mistakes compared with a full exhaustive sequence search that Eq. 2 is guaranteed to find, with the two methods agreeing 99.99% of the time.

Of the remaining child variants, those that do not have a matching allele with at least one of the parents (i.e. variants in Zmnone∪Zfnone) are considered Mendelian violations.

The only remaining unaccounted-for variants are ZHETALT = Zmone⋂Zfone, which are heterozygous child variants composed of two non-reference alleles. Among ZHETALT variants, those where PZm and PZf choose a different child allele are considered Mendelian compliant, since the two alleles can be separately matched with one or the other parent, and are thus added to the ZHETALT-CONSISTENT subset. Similarly, ZHETALT variants where the child’s phasing vectors choose the same child allele for both parents are added to ZHETALT-INCONSISTENT.

So far the steps above have focused on finding Mendelian violations at the child’s variants that are not homozygous reference. To test for Mendelian compliance at positions where the child is homozygous reference (i.e. 0/0 variants which are omitted from the child’s VCF), the following steps are taken. Any of the parents’ variants M\X and F\Y that were excluded from Eq. 3, but contain no homozygous alleles (i.e. heterozygous alternate or homozygous alternate variants) are automatically counted as Mendelian violations. These are in other words parent variants that are failed to matched with a child haplotype in Eq. 2. For heterozygous parent variants excluded in Eq. 2 where the parent has a reference allele, we naively check (akin to step 6) whether the other parent has a reference allele at this position, and whether the child is homozygous reference at this position. If both conditions are true, we consider the variant as Mendelian compliant assign it to MHETREF-CONSISTENT (or FHETREF-CONSISTENT). Otherwise the variant is considered a Mendelian violation. Otherwise the variant is considered a Mendelian violation and assigned to MHETREF-INCONSISTENT (or FHETREF-INCONSISTENT).

Finally, the number of Mendelian compliant variants is tallied as

\# Mendelian compliant = |Zmboth⋂Zfboth| + |(Zmboth⋂Zfone)∪(Zmone⋂Zfboth)| + |ZHETREF- CONSISTENT| + |ZHETALT-CONSISTENT| + |MHETREF-CONSISTENT ∪ FHETREF- CONSISTENT|.

\# Mendelian violations = |Zmnone ⋂ Zfnone| + |((Zmone⋂Zfnone)∪(Zmnone⋂Zfone)) / (ZHETREF- CONSISTENT) | + |ZHETALT-INCONSISTENT| + |(MHETREF-INCONSISTENT ∪ FHETREF- INCONSISTENT)|

#### 4.6.2. Inferred Genotype Accuracy

While Mendelian violation rate in trios can act as an indirect proxy for variant calling errors in variant caller benchmarking, previous studies have shown that the genotyping error rate of a variant calling pipeline can be statistically estimated based on variant calling data from a family trio64–69,77–79. Such approaches have been applied in practice previously80–82. We expand on previous work in two ways. First, we treat the frequencies of different parental genotype combinations as independent. As a result, our model is able to fit aggregate data of variants that are not necessarily in Hardy-Weinberg equilibrium. This parametrization of the true frequencies makes our model more flexible than previous approaches, which assume global allele frequencies^65–68,77,79,83^. Secondly, previously published models either assume an error rate independent of the true genotype^64,66,69,79^ or model the confusion matrix with only a few parameters^65,67,68,77,78^. In contrast, we estimate the frequencies of the trio genotypes and all individual genotyping error rates to infer an aggregate genotype confusion matrix over all variants for each pipeline. The resulting genotype confusion matrix can then be used to compute the genotyping accuracy of a pipeline for each of the three possible genotype values.

The implementation of the genotype accuracy estimation algorithm used in this study is implemented very similarly to that detailed in our preprint at bioRxiv^84^.

For simplicity, we only consider the case of unphased, bi-allelic genotypes, 0/0, 0/1 and 1/1. There are 33 = 27 possible trio genotype combinations, of which 15 are Mendelian compliant. We use T to denote the set of 27 observable genotypes and t to denote the set of 15 Mendelian compliant genotypes. Similarly, G is an element of T and represents an observed trio genotype, e.g. (0/0, 0/1, 0/1), which denotes that the mother is homozygous reference, while the father and the child are heterozygous alternate, and g is an element of t and denotes a Mendelian compliant trio genotype.

Our model is based on the following premises. We assume that all variant loci are subject to the same genotyping error profile, defined by an (unknown) 3-by-3 genotyping matrix E, where the rows (and columns) correspond to true (and called) genotypes, 0/0, 0/1, 1/1. Under this assumption, we are allowed to aggregate all called trio genotypes G, since the collection of their counts {NoG} is a sufficient statistic, where the o superscript indicates that these values are the observed counts. We disregard phred-scaled genotype likelihood (PL) values in this analysis because, although they convey information about the confidence of the variant caller, their limited availability for loci with homozygous reference alleles would prevent us from using the majority of the variant loci. This model explains the observed trio genotype G for a given variant called by the pipeline by positing a generative model comprising a mixture of 15 categorical distributions, each characterizing the chance of G being called in place of an unknown true trio genotype g (Supplementary Fig. 17).

**Supplementary Figure 17.**
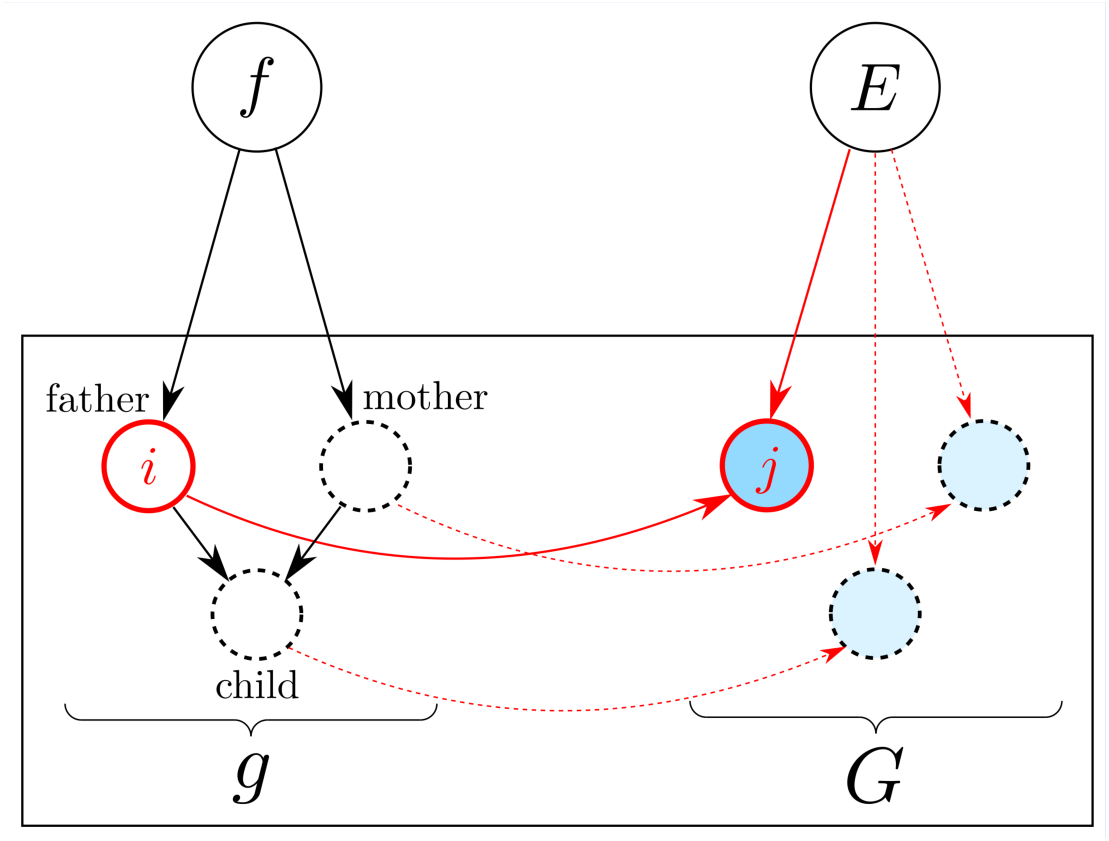
Graphical representation of the model. Parameters, f (parental genotype frequency), E (genotyping rates), determine the distribution of the hidden variable: the true genotype trio g, and the observed genotype trio G. The true genotype i ∈ I = {0/0, 0/1, 1/1} of a variant in the genome of each family member undergoes an independent transition, subject to the same genotyping rates E, resulting in called genotype j ∈ I.

To carry out the inference we use an Expectation-Maximization (EM) algorithm to fit the mixture model to {NoG}. In this model, the data is assumed to be a result of the following generative procedure. For each variant: 1) A true parental genotype combination is selected from a categorical distribution (defined by frequencies f), and a child genotype is picked according to the rules of Mendelian inheritance. This yields one of the 15 trio genotypes that are consistent with Mendelian inheritance. 2) All three individual true genotypes in the trio are separately subjected to the “observation process” (represented by the matrix E), which maps it to an observed genotype. This either changes the genotype (introducing an error, e.g. 0/1→1/1) or leaves it unchanged (making it a correct call, e.g. 0/1→0/1). After we obtain maximum likelihood estimates of f and E, we calculate the estimated genotype confusion matrix, and additional derived metrics. Below, we give a detailed description of our model and the corresponding EM algorithm.

##### Model details

###### Parental genotype frequencies

Let f = {fpar} denote the frequencies with which each parental genotype combination par, is present in the set of variants under investigation, where par is a sorted combination of the parents’ genotypes, i.e. one of (0/0, 0/0), (0/0, 0/1), (0/0, 1/1), (0/1, 0/1), (0/1, 1/1) and (1/1, 1/1). This means the probability of the mother having g1 and the father having g2 genotypes is P ((g1, g2) | f) = fpar(g), where par(g) = sorted(g1, g2) selects the genotypes of the parents and sorts them.

Here f is subject to normalization, i.e. Σ*_par_ w_par_ f_par_* = 1, where the normalization weight *w_par_* is 1 if *g*_1_= *g*_2_, and 2 if *g*_1_≠ *g*_2_, in *par* = (*g*_1_, *g*_2_). In order words, we assume that the frequency of (g, g’) is equal to the frequency of (g’, g), meaning that we implicitly assume that the two parents come from the same population.

###### Child genotype frequencies

Given the genotypes of the two parents, (g1, g2), the genotype of the child g3 has a well defined categorical distribution, dictated by the rules of Mendelian inheritance. Let Cg denote this probability.

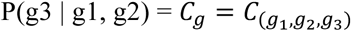

The values of Cg are known, e.g. C_(0/0, 0/1, 0/1)_ = 0.5, because parents with 0/0 and 0/1 genotypes produce a child with 0/1 genotype 50% of the time.

###### Trio frequencies

Combining the parental and child genotype distributions allows us to write the distribution of the trio genotype g as

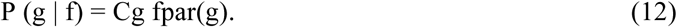

###### Error rates

Let E = {Ei,j : i, j ∈ I} denote the probabilities of calling the genotype j when the true genotype is i for a variant. We assume that genotyping errors have identical rates but happen independently for the three members (p = 1, 2, 3) of the trio. This allows us to write the probability of each type of (mis-)genotyping event as the product

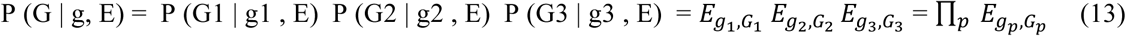

where E is subject to the normalization Σj Ei,j = 1, ∀ i.

##### Inference

###### EM solution

From the observable data we can extract the total counts of each observed genotype triplet regardless of Mendelian compliance:

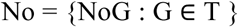

Explicit information about the true trio genotypes g ∈ t is missing from the observed counts No. Our ultimate goal is to estimate this hidden data, i.e. the joint counts of true and called genotype trios

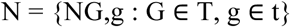

such that ΣgNG, g = NoG, ∀ G

Multiplying the conditional probabilities of Eq. 12, 13 yields the joint and conditional distributions of observed (G ∈ T) and hidden (g ∈ t) variables,

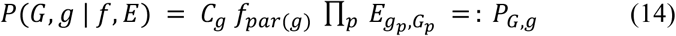

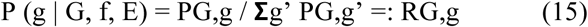

where RG,g can be understood as the “responsibility” of the hidden component corresponding to the true genotype trio g for the observed genotype trio G.

We use an EM approach to fit the model parameters (f and E) to the observed data No. In the E-step, we calculate R, using the formulas in Eq. 14 and 15. This is followed by the M-step, where we update the values of f and E using the formulas:

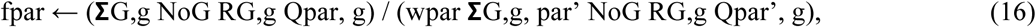

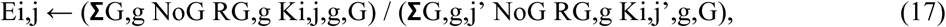

where Qpar, g is 1 only if par = par(g) and 0 otherwise, and Ki,j,g,G is the number of times the genotype transition i → j appears in the genotype trio transition g → G; for example K0/0, 0/1, (0/0, 0/0, 0/1), (0/1, 0/1, 1/1) = 2 because both the mother’s and the father’s genotypes went through the mis-genotyping event 0/0 → 0/1 but not the child’s.

###### Benchmarking metrics

The EM algorithm converges to a local optimum. In practice, when running our method on trio datasets from different initialization points, we always reach the same solution. We use the MLE solutions fMLE and EMLE to calculate the most likely joint histogram N, using

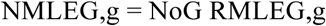

where RMLEG,g is a function of No, fMLE and EMLE, calculated from Eq 14 and 15. We define the estimated genotype confusion matrix for each family member p as

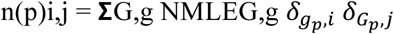

where *δ* is the Kronecker-delta, evaluating to 1 only if the quantities in its subscript are equal, and giving 0 otherwise.

From these confusion matrices, we define the true positive, false, negative, false positive and “mixed-up positive” counts as

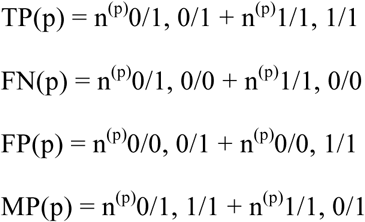

for each family member p, from which any benchmarking metric can be directly evaluated, including precision, recall, and F score, with the following formulas:

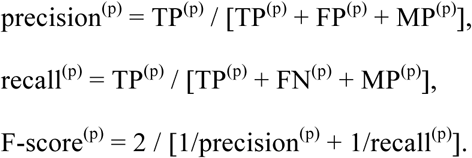

###### Input data

The required input data for our Genotype Accuracy Inference method are the raw frequencies (counts) of the variants with different observed trio genotypes. After calling the variants of the three members of the trio separately, any aggregation method that produces such counts is admissible. For this study, we decided to use our variant representation consolidation method, VBT, for counting Mendelian violations, as described in section 4.6.1. The detailed output logs produced by this method allows us to extract the number of variants with different observed trio genotypes in the following way. The pipeline used is illustrated in Supplementary Figure 18.

**Supplementary Figure 18:**
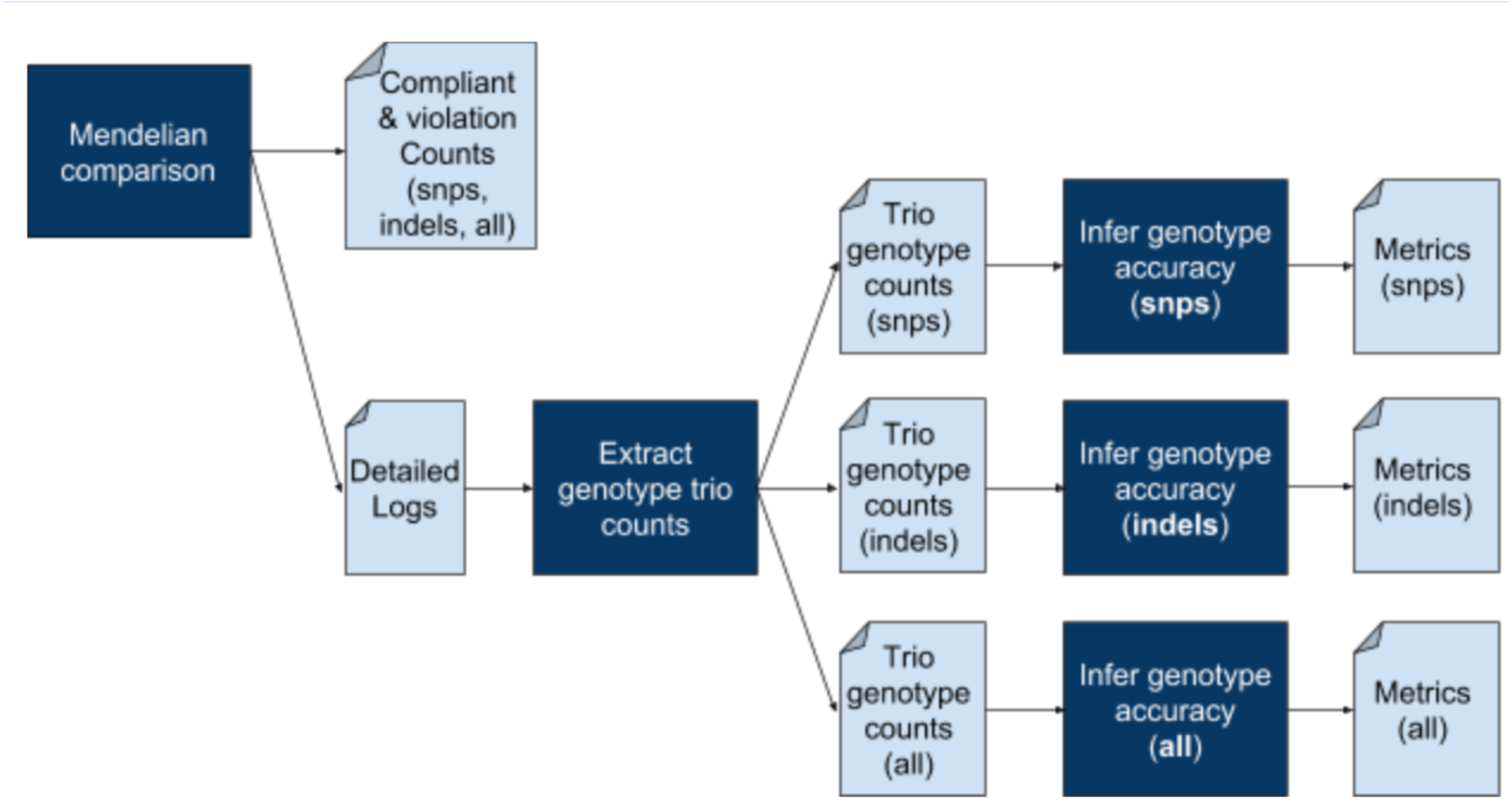
Illustration of the input preprocessing pipeline for our Genotype Accuracy Inference tool, run separately for snps, indels and all variants combined.

The output of our Mendelian violation checking tool produces 27 (and +1 for multi-allelic sites) raw frequencies (counts) for each of the following type of variants: snps, insertions, deletions, complex variants. (We combine the two tables in the output by taking only the counts from the “compliant” table that correspond to the 15 Mendelian compliant trios and only the counts from the “violation” table that correspond to 12 trios that violate Mendelian inheritance rules. Entries outside of these selected ones are non-zero only because of the variant merging step. Excluding them from the downstream analysis avoids overcounting them.) We add the counts of insertions, deletions and complex variants to obtain the counts of indels.

The variant callers we employed did not report homozygous reference variants. As a result, we had no information about the number of variants sites present in all three people with homozygous reference genotype, i.e. N°G is unknown for G = (0/0, 0/0, 0/0). To avoid this missing data biasing our results, we used 1 million in place of the counts of these uniformly homozygous “variants”.

#### 4.6.3. Datasets for benchmarking using related genomes

We used two sets of trios in the experiments based on the related genomes, the CEU Trio: NA12878 (daughter), NA12891 (father), and NA12892 (mother), and the Ashkenazi Jewish trio: HG002 (son), HG003 (father), and HG004 (mother). Data for the CEU trio are 40x coverage 2×100bp (available from the 1000 Genomes FTP site, http://ftp.1000genomes.ebi.ac.uk/), while the Ashkenazi Jewish trio is based on 50x coverage 2×150bp data obtained from the GiaB FTP site (ftp://ftp-trace.ncbi.nih.gov/giab/ftp/).

### 4.7. Benchmarking Against SNP Array Genotyping

Benchmarks were done on 3 samples (NA12878, NA12891, and NA12982) from the CEU trio sequenced to 40x coverage using 2×100bp reads. SNP array data was taken from 1000 Genomes Phase 3 dataset, obtained from both the Illumina Omni platform (ALL.chip.omni_broad_sanger_combined.20140818.snps.genotypes.vcf.gz) and the Affymetrix SNP6.0 platform (ALL.wgs.nhgri_coriell_affy_6.20140825.genotypes_no_ped.vcf.gz). The two platforms indicated very discrepant results, both in terms of absolute accuracy levels, and relative performance of the tools, with the Freebayes results showing the largest amount of variation (Supplementary Table 8).

However, both platforms across all 3 samples suggest that Graph Genome Pipeline achieves the highest F-score, with the exception of the sample NA12878 and the Omni platform, which give a small advantage to the combination pipeline of Graph Aligner and GATK HaplotypeCaller (Supplementary Fig. 19, Supplementary Table 8).

**Supplementary Figure 19.**
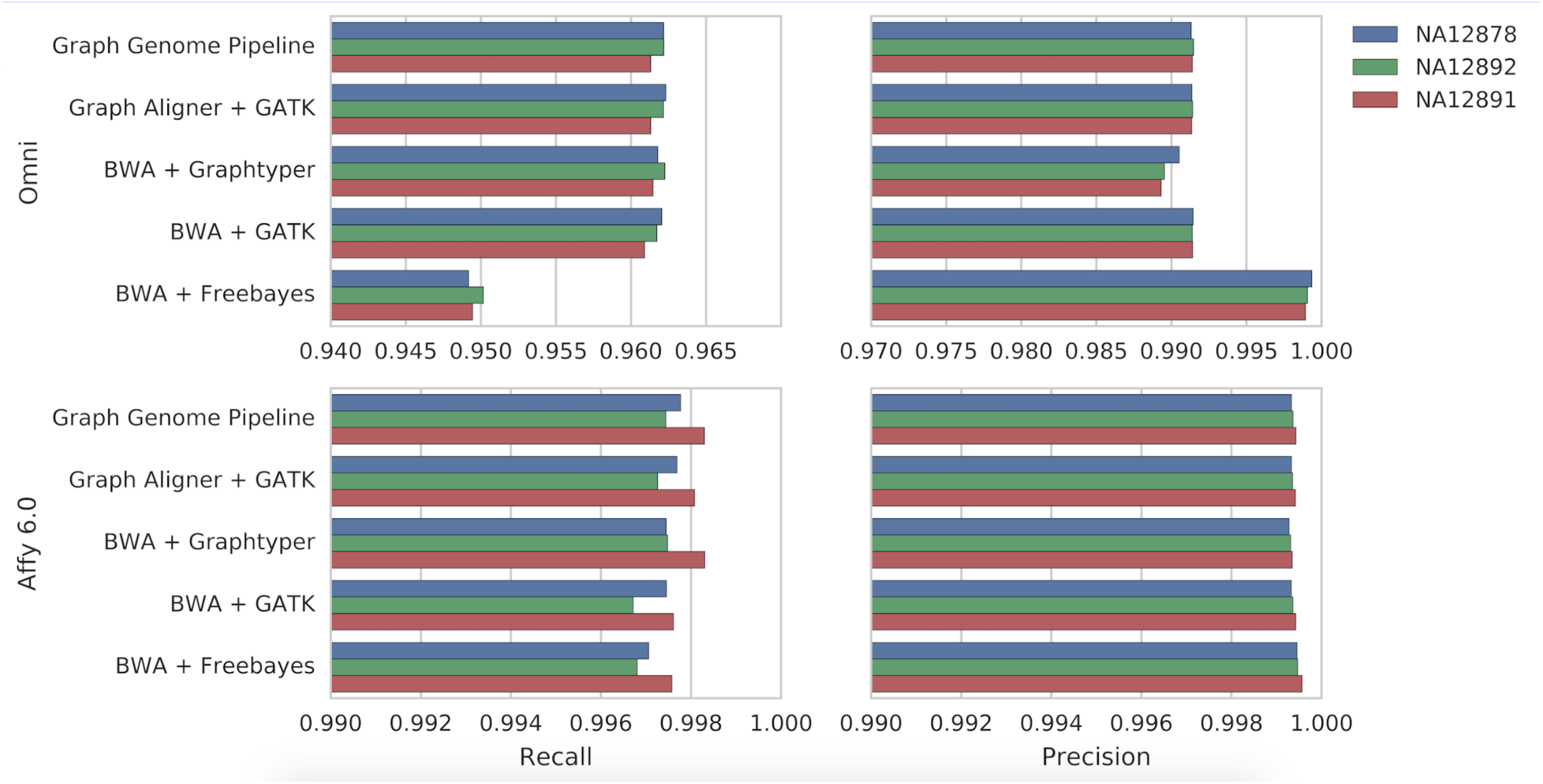
Comparison of the variant calling results against SNP Array data.

## 5. True Variants Missing from GiaB Truth-sets

Manual inspection of a random subset of the false positive variants identified by the Graph Pipeline revealed that a significant portion of those calls exhibit patterns that suggest them to be true variant sites, missed by the GiaB truth set due to reference bias affecting the alignment step of the truth set generation. Namely, the alignments produced by the linear aligner used for the truth set show a gradual drop in coverage as the variant site is approached (Supplementary Fig. 20). Furthermore, the linear aligner often places a few reads in support of the variant, sometimes with a soft-clip covering the variant site. Affected regions (in which the coverage drop is evident) are mostly one hundred to several hundred bases long, and are mutation dense (usually < 30 variants). In contrast, graph-based alignments tend to exhibit a more even coverage patterns across those regions (Supplementary Fig. 20). All of the considered variants were genotyped as being heterozygous reference/alternate alleles by the Graph Genome Pipeline (Supplementary Table 14).

**Supplementary Figure 20.**
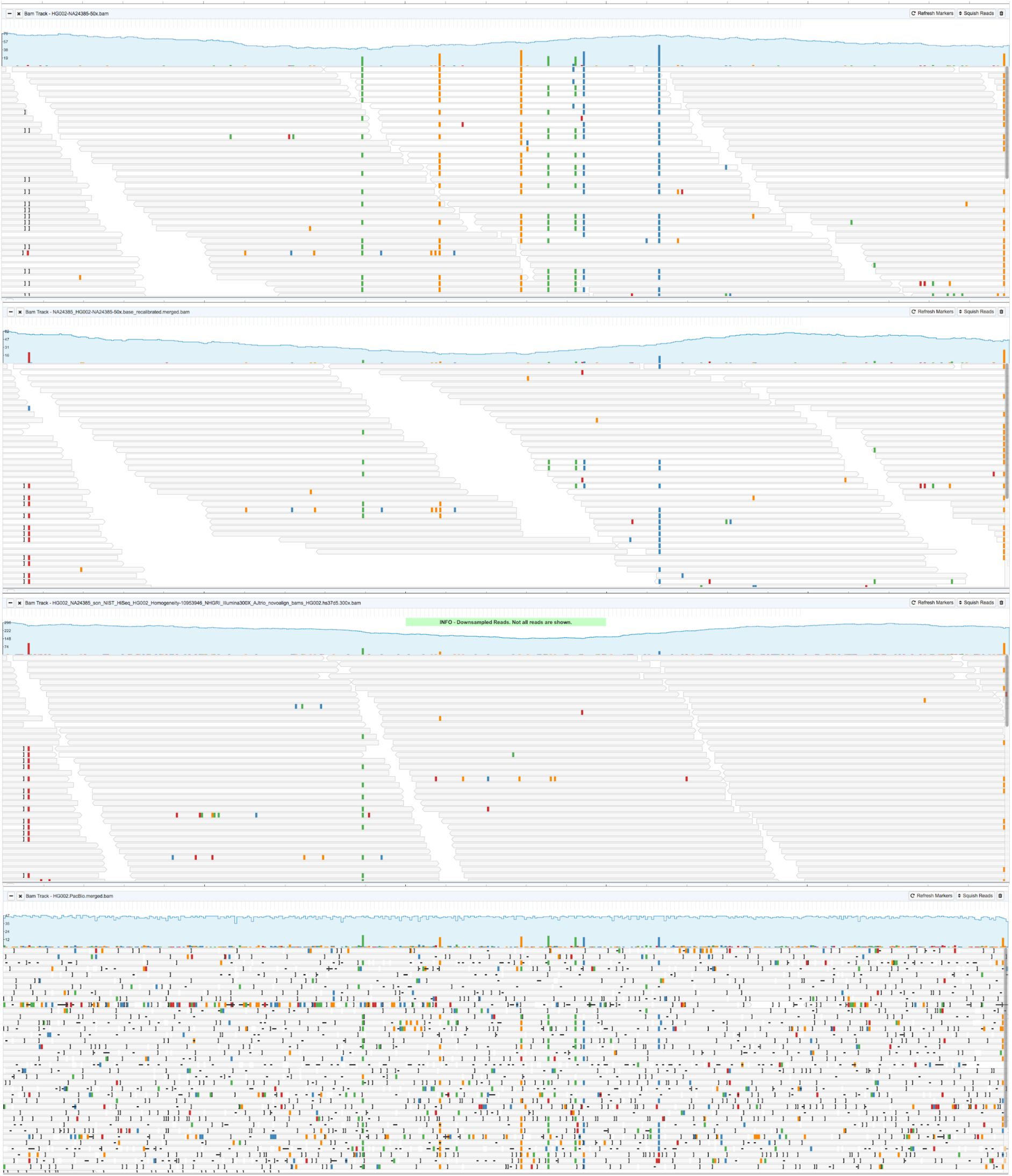
Alignments in a highly mutated region. Image shows Illumina 2×148bp read alignments by Graph Aligner (first from the top), BWA-MEM^27^ (second from the top), Novoalign (http://www.novocraft.com/products/novoalign/) (third from the top), and PacBio reads aligned by BWA-MEM (bottom), in HG002 sample (region 1:99,567,636-99,568,117). PacBio and Novoalign data was obtained from Zook *et al.^29^*. Linear aligner with short reads show a drop in coverage, indicating a loss of an allele, while Graph aligner, and long reads show that the region is dense with mutations.

The observed pattern can be explained by the effects of reference bias. Most short read mappers prioritize read alignment based on the alignment scores, which reflect the distance between the read’s sequence and reference genome. When the edit distance is large, or when a read aligns to multiple reference genome loci with high similarity, standard read mappers have problems placing the reads correctly. In some cases the reads may be placed correctly, but are assigned a low mapping quality score, as they match (almost) equally well at several genomic loci. Mapping against a graph genome reference has the advantage of avoiding mismatch penalties for all variants which are already known and are a part of the graph. This effect is amplified in regions dense with variants on the same haplotype, i.e. where a read contains several variants (Supplementary Fig. 20). As Graph Aligner utilizes paired end information (Supplementary Information 3.1), the range in which a known variant affects the mapping is extended beyond the length of a single read to the full insert size of the read pair.

There are several different mechanisms by which a graph genome reference assists the discovery of a variant:

- If the variant is present in the graph, reads containing this variants have an increased alignment score (a would-be mismatch against a linear reference becomes a match)
- If the variant V_1_ is on the same haplotype with other adjacent variants V_2_, V_3_, … V_n_, which are known and placed in the graph, reads containing both the variant V_1_ and the variants V_2_, V_3_, … V_n_ have increased alignment scores
- Increased alignment score can change the choice of the alignment location, allow mapping of reads that would otherwise go unmapped, or boost the mapping quality estimate for a read.

### 5.1. Automated annotation of potentially missed calls

Four of the five GiaB samples have been sequenced using orthogonal technologies, namely by PacBio Long Reads and 10x Genomics Linked Reads sequencing^29^. Both of these technologies can provide additional insight regarding the validity of these variant calls.

PacBio Long Read sequencing^29,62^ produces reads that are several kilobases long, making it easier to match a read across a short, highly divergent region. Furthermore, alignment methods for these reads tend to be less sensitive to differences between the read and the reference, in order to compensate for the typically higher error rates. Consequently, long read alignments are less affected by reference bias at the length scales studied above (regions of a hundred to several hundred bases) and often support the existence of the same variants as our graph based method.

The 10x Genomics Linked Reads technology (http://www.10xgenomics.com) utilizes molecular barcoding to mark short reads originating from the same DNA molecule of up to several hundred kilobases long. Once aligned, the reads can be accurately phased and tagged with the information regarding the copy of the chromosome they originated from. A site with the homozygous reference genotype can be expected to be covered with a mixture of reads tagged as belonging to either of the chromosomes. Conversely, if all or a large majority of reads covering a site all phase to the same copy of the chromosome, it is likely that one of the alleles is missing. Such a pattern would be consistent with mapping algorithm failing to map, as would be caused by reference bias.

We have utilized PacBio and 10x data to devise a strategy for identifying variant calls made by our graph-backed pipeline that are marked as false positive calls against the GiaB truth sets, and which may potentially be real variants missed during the truth set construction. We defined the following criteria:

- Long reads show strong support for the variant being present (at least 20% of the reads support the variant allele)
- 10x reads fail to properly support homozygous reference genotype (more than 90% of the reads have the same phase or are unphased)
- Coverage at the variant site in the short read alignments produced by the Graph Aligner is within four median absolute deviations from the median across the entire set
- Coverage at the variant site in the long read alignments produced by the BWA-MEM is within two median absolute deviations from the median across the entire set.

Coverage limitations were introduced to exclude sites that may contain an extra allele due to a potential copy number variation event. We have limited the analyses to SNPs, where we can directly calculate the support in PacBio reads, as opposed to INDELs that would require a more elaborate calling method. Following this approach, 26-42% of calls marked as false positive calls made by our pipeline are identified as potentially correct calls (Supplementary Table 2).

We selected a subset of these variants for a Sanger sequencing validation experiment (Supplementary Tables 2 and 14). The criterion for exclude of variants was proximity to heterozygous INDEL variants, which cause difficulties in the process of Sanger sequencing^85^. For each variant site we set a region of +/- 100bp, and merged all overlapping regions originating from different variants. We further extended each of these merged regions by 200bp each side, followed by another round of merging of any overlapping regions. From this set, we exclude all regions that contain multiple indel variants. Results of the Sanger sequencing experiment are provided in Supplementary Table 14 and summarized in Supplementary Table 2

**Supplementary Table 2.** Counts of calls marked as false positives made by the graph genome pipeline against GIAB High Confidence truth-sets, calls identified as potential true variation sites, calls selected for Sanger sequencing validation, and the calls that passed the Sanger sequencing validation.

| Sample | Total FP calls | Suspect FP calls | Calls selected for validation | Validation sequencing successful | Validated calls |
| --- | --- | --- | --- | --- | --- |
| HG001 | 2,044 | 531 | 351 | 294 | 187 (63,6%) |
| HG002 | 2,107 | 888 | 598 | 510 | 306 (60%) |
| HG003 | 2,396 | 826 | 0 | - | - |
| HG004 | 1,852 | 653 | 0 | - | - |

GiaB samples HG002, HG003, and HG004 form a family trio (the GiaB Ashkenazi Jewish trio). We made use of this to perform a Mendelian inheritance consistency check on the SNP calls we believe to be real variation missing from the truth-set. Out of 888 such calls, 816 follow a Mendelian inheritance pattern, while 72 are violations, further supporting the notion that the majority of these calls are indeed real variants.

Finally, we have examined the variants and the reads supporting them in order to deduce if their discovery was, as we suspected, assisted by the graph genome reference, and if so by which mechanism (Supplementary Table 3). About half of the calls (47%) are previously known variants contained in the Global Graph Reference. The other half is mostly made up of variants adjacent to known ones (31.4%) or assisted by paired end mapping (17%), while few percent could not be directly linked to the graph (3.7%). Of all the reads supporting these variants, 60% would have been mapped to a different location without the effect of the graph, while for the rest the effect from the graph was related to the mapping quality.

We did not attempt the converse experiment, i.e. validating the existence of any of the variants from the GiaB truth-set that were not called by the Graph Genome Pipeline, as we have not observed any systematic patterns of false positive calls in the GIAB truth sets that are correctly avoided by the Graph Genome Pipeline.

**Supplementary Table 3.** Likely true variants missing from the GiaB truth-set and their relationship to the Global Reference Graph.

| Sample | Variants in the graph | Nearby variants in the graph | Mates map across variants in the graph | Other |
| --- | --- | --- | --- | --- |
| HG001 | 305 | 157 | 55 | 14 |
| HG002 | 392 | 299 | 165 | 32 |
| HG003 | 363 | 274 | 161 | 28 |
| HG004 | 280 | 189 | 151 | 33 |

### 5.2. Sanger Sequencing Validation Experiment

Primer design, PCR amplifications, Sanger sequencing and variant calling were performed by Quintara Biosciences, CA using standard protocols. Primers were designed using Primerquest (http://www.idtdna.com/Primerquest/Home/Index) around 200-250bp away from the variants of interest. Q5 High-fidelity 2x master mix from NEB (CAT# M0492L) was used for PCR (1 cycle:95C 3 mins; 35 cycles: 95C 30s, 57C 30s, 72C 30s; 1 cycle: 72C 10 min and then 4C) with 25ng genomic DNA in 25ul reaction. BigDye Terminator v3.1 cycle sequencing kit from ThermoFisher Scientific (CAT# 4337455) was used for Sanger sequencing. Sequencing analysis 5.2 software from ABI was used for base calling.

## 6. Structural variation (SV) analysis

### 6.1 Genotyping Through Read Counting

Graph Genome Pipeline’s variation-aware structure allows reads originating from non-reference contigs to map directly onto alternate paths of the human genome reference. When a read is mapped to an alternate path, additional information regarding graph alignment is reported using custom annotation tags, since built-in fields of the BAM file format (https://github.com/samtools/hts-specs) are insufficient to accurately represent the alignment of that read. Alignment paths of reads can simply be read from these custom annotation tags for each region of interest. Then, by just counting the number of reads aligning to the reference vs. the alternate path at the breakpoints of the variant of interest, the genotype of the variant can be intuitively inferred from the ratio of reads (reads mapping to the alternate path/total reads). For the three clusters of true SV genotypes - homozygous reference, heterozygous, homozygous alternate - the expected proportions of reads aligning to the alternate path are 0, 0.5 and 1, respectively. Therefore, the genotyping thresholds were set right in the center between the expected cluster means, i.e. at 0.25 and 0.75. As shown in Figure 4b in the main manuscript, there are three genotype clusters as expected, which enables us to genotype SV events using a threshold-based clustering algorithm.

In practice, the read-based SV genotyper uses a graph alignment BAM file, iterates over a set of events from a VCF file and it infers the genotype of each event based on the observed mapping ratio of reads using 0.25 and 0.75 as thresholds. Since the aligner provides all this information already, genotyping is basically just retrieving reads and reading their tags from a graph-aligned BAM, which requires very little overhead.

### 6.2. SV reconciliation

Comparing variants between two different VCFs is a difficult problem because a variant can be represented in multiple different ways86. For example, a single SV event can be represented as a combination of several simpler INDELs due to the local phasing of subcalls into different positions. The naive way of comparison, which is to declare variants with identical position, reference and alternate alleles as the same, fails to identify equivalent variants with non-identical representations. Vcfeval^75^ can resolve variant representation differences but is unable to handle larger variants that span larger regions due to the exponential time and space complexity of these comparison algorithms. Longer variants such as SVs are either filtered out from comparison or the sites that contain SVs are skipped. In addition, vcfeval requires an exact match (100% identity) between the two variant sets. Variants are considered to be mismatching and marked as FP/FN even with a single base pair difference. For these reasons, we developed the following sequence comparison-based mismatch-aware SV reconciliation strategy to evaluate the performance of SV callers.

From a VCF produced by an SV caller, we attempt to infer the genotypes for a set of known SVs by considering all possible paired paths through the VCF. For each known SV, we first define the region boundaries by adding 100bp of padding around the SV breakpoints. Then for each region, we clip the reference contig from the GRCh37 human reference and build multiple sequence pairs by applying all possible variant phasing combinations that reside within the region boundaries. A total of 2^Number of small^ variants pairs of haplotype sequences (sequence tuples) can be generated within a region, since for each called variant in the sample, you can either apply its given phase or invert it. We then compare each haplotype sequence tuple against a single sequence produced by applying the ALT allele of the known SV within the region boundaries. However, it is not always expected that we observe a contig that is 100% identical to the known SV we are trying to genotype. This can be due to the known SV having an error, the SV caller making a base calling error (but calling the presence of the SV correctly), or the individual harboring a true variant within the SV. Therefore, as opposed to strict matching, we allow inexact matching by calculating the Longest Common Subsequence (LCS) between them and applying an identity cutoff of 95%. The match ratio is then calculated by the LCS of the two sequences divided by the maximum length between them. Finally, we identify the most likely genotype for a given SV event using a 2-step maximization. Step 1: we maximize the count of matches per sequence tuple. If both sequences in the tuple match with the known SV sequence, we treat its genotype as homozygous for the SV. If there is only a single match, we estimate its genotype as heterozygous for the SV. Step 2: when multiple sequence tuples homozygous or heterozygous for a known SV can be found, we maximize the match ratio of single sequences. For example, if the match ratio of alleles are (95%, 40%) and (99%, 20%), we select the latter since the ALT allele match ratio (99%) is higher. In the end, we obtain the genotypes for the SV event with maximized match ratios for each region.

### 6.3. SV Benchmarking

We compared the performance of Graph Genome Pipeline and the read counting method (explained in Section 6.1) that we developed against four SV genotyping pipelines: the genotyping module of Delly287, BayesTyper^11^, Graphtyper^12^ and HaplotypeCaller^57^. Delly2, BayesTyper, Graphtyper and both of our methods make use of known SVs, whereas HaplotypeCaller cannot. All tools were run with default parameters.

SV benchmarking was conducted using 49 samples from the EUR superpopulation of the 1000 Genomes Project. The 49 samples were selected because high coverage sequencing data from them is available from the Coriell study and their SVs have been individually genotyped5. In addition to that, we repeated our experiments using low coverage data on the same 49 individuals but only with methods that performed well on high coverage data. The results from each SV caller were compared against the truth genotype reported by the 1000 Genomes SV project5.

In the case where the tool reports variants in different representations from the input VCF, we used the variant reconciliation method detailed Supplementary Information 6.2. Our results show that the read counting method and Graph Genome Pipeline have the highest F-score (96.3% and 96.4% respectively) and concordance (96.6% and 96.7%) followed by BayesTyper with 95.3% F-score and 95.6% concordance (Fig. 4c in the main text; Supplementary Table 4). Although Graphtyper has a very high precision (94.5%), it suffers from low recall (26%) because of its limitation regarding variant length. The size of the longest SV, either gain or loss, genotyped as heterozygous or homozygous alternative by Graphtyper is 8,181bp, while our dataset contains events larger than 30,000bp. If we group SVs with non-reference genotypes in the truth set into three bins such that each one is placed into one of 100-1,000bp, 1,000-8,000bp, 8,000-33,000bp ranges, the concordance value of Graphtyper is 0.497, 0.076, and 0.006 respectively for each bin. This shows that the concordance value decreases constantly as the SV size increases. Also, it is worth mentioning that HaplotypeCaller and Graphtyper clearly state that they are not designed to handle large events.

**Supplementary Table 4.** Precision and F-score could not be computed for HaplotypeCaller since it called no SVs among the SV truth set.

| Method | Coverage | Alignment | Benchmarked with | Precision | Recall | F-score |
| --- | --- | --- | --- | --- | --- | --- |
| Delly | Low | Graph | Genotype comparison | 0.958 | 0.451 | 0.613 |
| BayesTyper | Low | Raw reads | Genotype comparison | 0.651 | 0.638 | 0.645 |
| Read Counting | Low | Graph | Genotype comparison | 0.890 | 0.791 | 0.837 |
| Graph Pipeline | Low | Graph | SV reconciliation | 0.976 | 0.627 | 0.763 |
| Delly | High | Graph | Genotype comparison | 0.990 | 0.568 | 0.722 |
| Delly | High | BWA | Genotype comparison | 0.994 | 0.567 | 0.722 |
| BayesTyper | High | Raw reads | Genotype comparison | 0.954 | 0.952 | 0.953 |
| Read Counting | High | Graph | Genotype comparison | 0.971 | 0.956 | 0.963 |
| Read Counting | High | SV-only graph | Genotype comparison | 0.980 | 0.966 | 0.973 |
| Graph Pipeline | High | Graph | SV reconciliation | 0.992 | 0.947 | 0.969 |
| GraphTyper | High | BWA | SV reconciliation | 0.945 | 0.260 | 0.407 |
| Haplotype Caller | High | BWA | SV reconciliation | - | 0.000 | - |

In addition, Delly2, BayesTyper, Graph Genome Pipeline and the read counting method was run on low coverage data from 1000 Genomes Project Phase 3 on the same individuals. In the low-coverage data, our approach is clearly superior in terms of both F-score and concordance (Supplementary Table 4). This indicates that alignment across the full variant with respect to the human reference is especially useful for low coverage data.

### 6.4 PacBio alignment across SVs (HG002)

We downloaded a set of 30x PacBio-sequenced reads from the son of the Ashkenazi trio (HG002), which was aligned using BWA-MEM by the Genome in a Bottle Consortium. The 50x Illumina reads used for the PrecisionFDA Truth Challenge were realigned using our graph pipeline and genotyped for existence of the 230 SVs. We were not able to observe a single locus where PacBio reads align clearly across the SV event. Reads were either completely soft-clipped around the deletion (Supplementary Figure 21, top), or fragments of reads were intermittently aligned across the event with inconsistent breakpoints with high variance (Supplementary Figure 21, bottom). In contrast, when an SV is known and included in the graph, Graph Genome Pipeline can readily align Illumina reads across them (Supplementary Fig. 22).

**Supplementary Figure 21.**
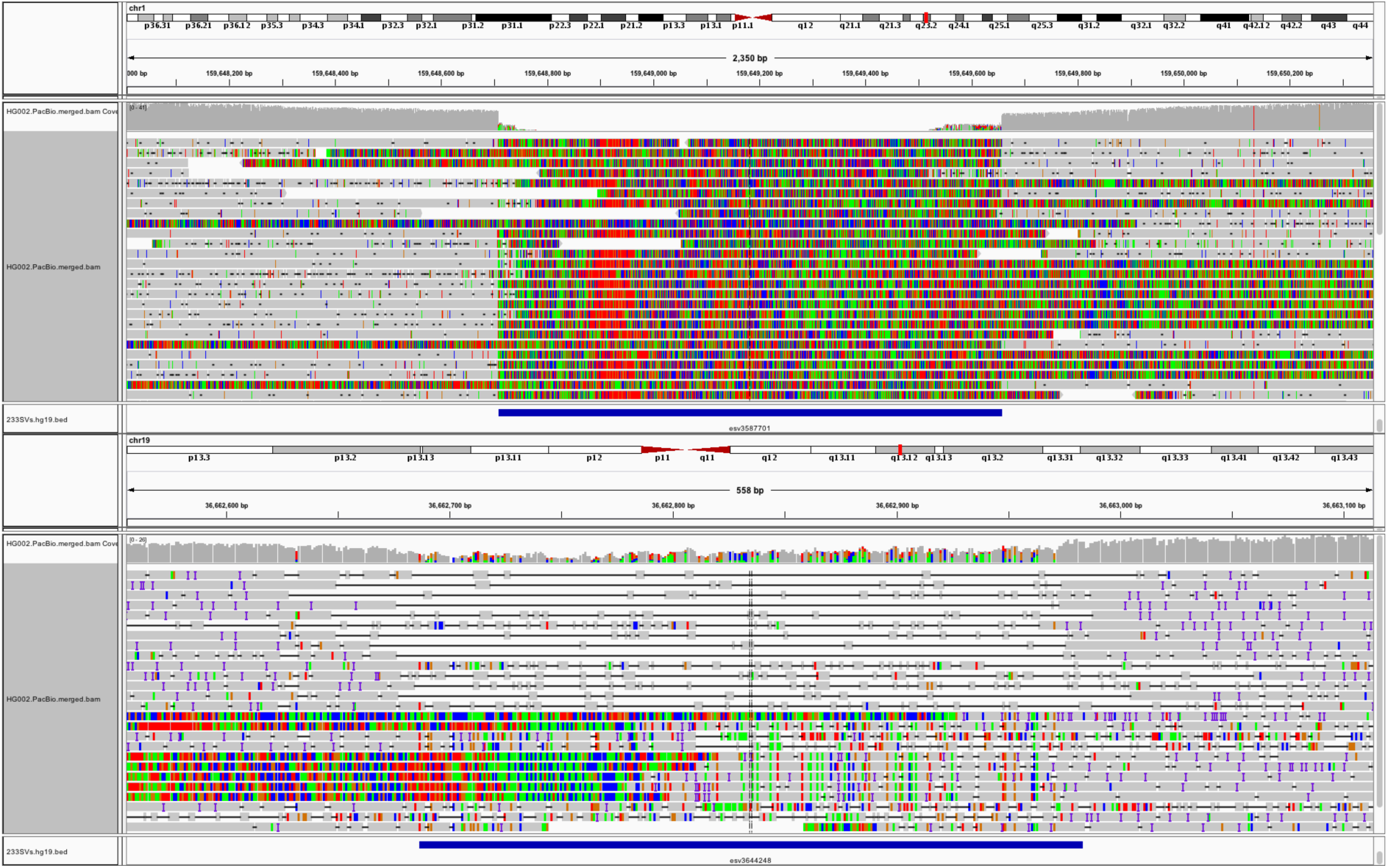
Representative examples of PacBio reads aligned using BWA-MEM with parameters tuned for PacBio reads. All PacBio alignments across the SVs fail in one of these two ways. Top: The portion of all reads that map across the deletion event are soft-clipped (esv3587701, GRCh37, chr1:159648708-159649658). Bottom: Some reads suggest the presence of a deletion event, but is fragmented by small stretches of aligned sequences, and suggest inconsistent breakpoints (esv3644248, GRCh37, chr19:36662810-36663112).

**Supplementary Figure 22.**
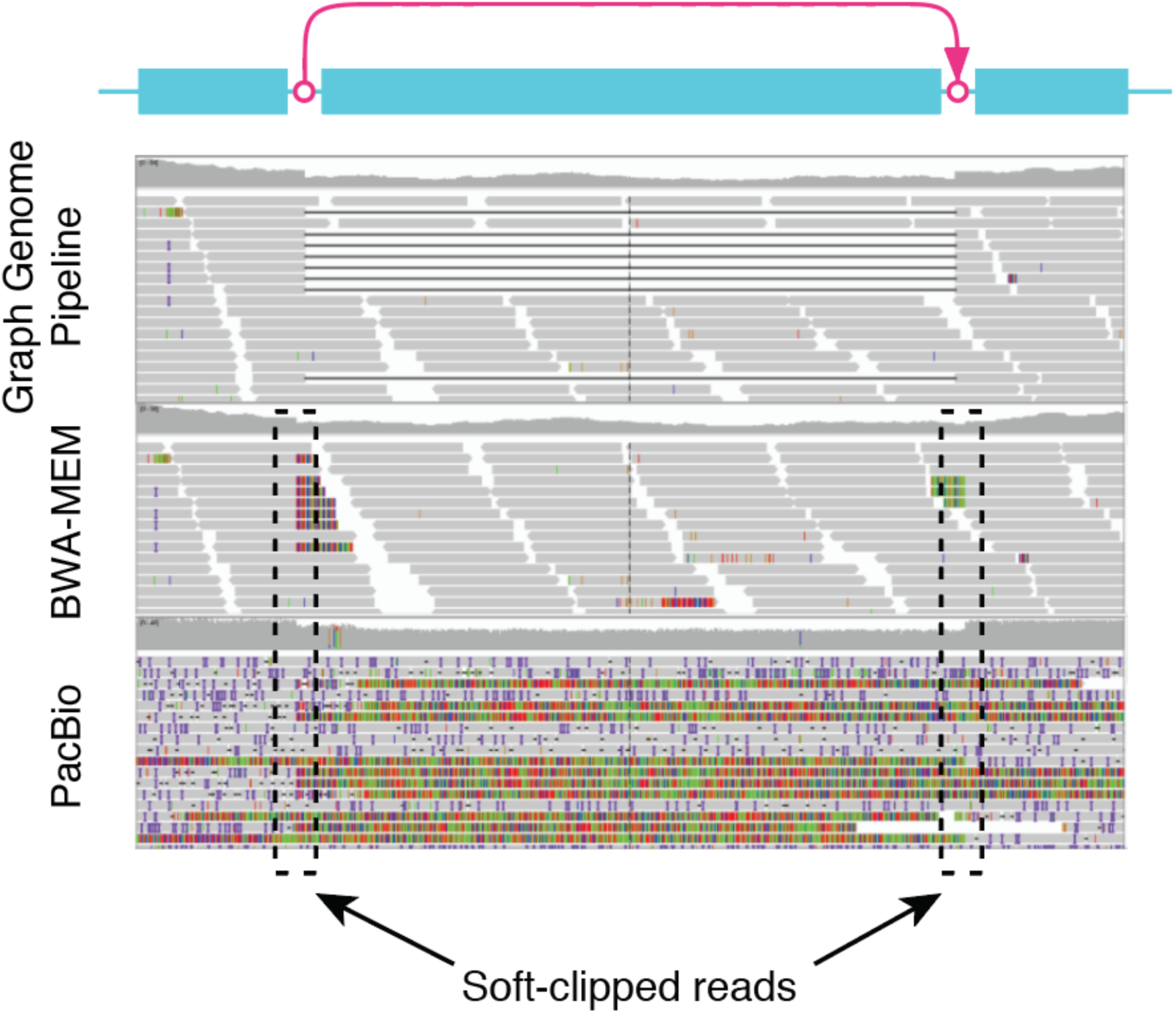
Example read alignment at an SV region (esv3642033/sbgsv207; 18:24571620-24572341 in GRCh37) in sample HG002 by different technologies. The top two panels show Illumina read alignment by Graph Aligner and BWA-MEM. The bottom panel shows the alignment of PacBio reads by BWA-MEM options recommended for PacBio data. This SV is in complete linkage disequilibrium with SNP rs1436904, which is significantly associated with breast cancer risk. Colored vertical bars at the tips of reads indicate soft-clipping. Whereas Graph Aligner (top panel) is able to correctly align reads across the SV, BWA-MEM fails to do so (middle panel). Similarly, BWA-MEM-aligned PacBio reads (bottom panel) using parameters optimized for PacBio (option ‘-x pacbio’ in BWA-MEM) also fail to align across the SV.

### 6.5. Ti/Tv of SNPs at SV breakpoints

SV breakpoints for Ti/Tv calculation were defined by setting read length to 0 (Supplementary Information 2.1.1). This chooses the “smallest” SV representation for each SV, leaving any potential flanking SNPs outside the SV branch. The rate of false positive SNPs at the 10bp closest to the SV breakpoint was estimated by assuming that randomly false positive SNPs have an expected Ti/Tv of 0.5, and solving the equation for false positive SNP rate *f* such that 0.5×*f* + (1 − *f*)×*2*.1 = 0.*79*.

